# SafeLens: A Statistically Controlled Nonlinear Machine Learning Framework for Characterizing Hidden Dynamics in Event-Locked Neural Activity

**DOI:** 10.64898/2026.10.31.763367

**Authors:** Christopher B. Minasi, Umair Hassan, Christopher C. Cline, Corey J. Keller

## Abstract

Machine learning has advanced rapidly over the past decade, yet event-locked neural activity is still analyzed predominantly with traditional statistics and linear models. While traditional linear methods provide computationally tractable statistical control for pre-selected features or trial-averaged waveforms, they miss subtle nonlinear and trial-varying neural dynamics. Nonlinear models can capture these complex patterns, but their feature reliance is opaque and notoriously difficult to threshold statistically. We introduce SafeLens (Searchlight Analysis of Fluctuation Events, Leveraging Extensive Null Sampling) to combine the power of nonlinear machine learning with the rigorous false-positive control of traditional statistical inference. At each position of a sliding window, a random forest learns to discriminate event-locked activity from interleaved resting-state neural activity using multiscale temporal derivatives. Randomized feature ablation inverts slope polarity in held-out trials to map the random forest models’ reliance on specific rising versus falling excursions across the time-frequency space. SafeLens then establishes an empirical null by running the same pipeline on separate resting-state recordings, training forests to discriminate rest from rest. This empirical null translates each channel’s complex nonlinear resting dynamics into a model-matched noise floor on the map’s linear reliance scale, allowing a single threshold to control the family-wise error rate against the channel’s own background structure rather than a summary statistic of its baseline. When we injected simulated cortico-cortical evoked potentials (CCEPs) into resting intracranial EEG recordings, SafeLens recovered ~2.5 times as many polarity-resolved deflections as trial-average thresholding and the closest data-driven method, maintaining its advantage under trial-to-trial variability while yielding ~5–11 times fewer false components, ~2–5 times lower mean boundary error, and superior single-trial latency tracking as a timing prior. In eight patients, CCEPs remained detectable above chance until 1,040 ms after stimulation, compared with 570 and 455 ms for those benchmarks. Together, these findings establish SafeLens as a novel solution for enabling nonlinear models to uncover subtle, trial-varying neural dynamics under strict statistical control.

**Author Summary:** Much of what we know about brain function comes from measuring how neural activity responds to specific events, from sensory stimuli and cognitive demands to targeted pulses of brain stimulation. Tracking these responses is fundamental for mapping how brain networks communicate in health and disease and for guiding interventions that modulate neural circuits. Because each response is buried in ongoing background activity and changes across repetitions, isolating it is difficult. Existing methods average repetitions inside short, hand-picked time windows, restricting researchers to seeing only the earliest, most stereotyped part of a response while leaving later, fluctuating activity invisible. Here, we introduce a new approach that combines the pattern-finding sensitivity of modern machine learning with the strict false-positive control of traditional statistics. Our method searches the period after each event without a preset window, using recordings of the brain at rest both to learn how a response differs from normal background activity and to measure what the model detects by chance alone. Using this tool, we show that human brain responses to electrical stimulation last over twice as long as previously recognized, revealing that brief perturbations trigger a longer cascade of neural activity than previously thought.

## I. Introduction

Evoked responses are measured by repeating a time-locked *event* and analyzing a physiological *recording* relative to the event’s onset. Event-locked studies account for more than thirty-three thousand publications indexed since 2015 (S1 Table) and roughly two thirds of the research articles in systems and cognitive neuroscience journals (S2 Table). The *event* can be anything from an external sensory stimulus or cognitive demand to a more direct perturbation such as a pharmacological challenge or an electrical or magnetic pulse delivered to the brain itself [1–5]. The *recording* is just as varied, ranging from direct captures of electrical or magnetic activity, such as scalp electroencephalography (EEG), magnetoencephalography (MEG), and intracranial EEG (iEEG), to slower indirect readouts, such as functional magnetic resonance imaging (fMRI), functional near-infrared spectroscopy, calcium imaging, and peripheral measures (e.g. pupil, muscle, or skin-conductance). Across these diverse event and recording modalities, the central challenge is the same, in that the response of interest is typically obscured by ongoing background activity that is often many times larger than the response itself [6] and must therefore be recovered from repeated trials. The standard practice is therefore to average many repetitions, since background activity uncorrelated with the event shrinks as 1/N over N trials while a response that repeats identically is preserved [7, 8]. But that arithmetic holds only if the underlying response is the same in every trial. When trials differ, the average is a composite that may not resemble any individual trial [9–11], and the cost falls on the most variable (often the latest and weakest) parts of the response, which the assumption of homogeneity dilutes.

Despite how frequently neural data violates this often implicit *assumption of invariance across trials* [8–14], much of the field can afford the cost of averaging since many neural recordings license further assumptions that buy them statistical power by pooling data across additional dimensions. Perhaps the most common of these is the *assumption of a known waveform shape*. The most widely known example of this is in the analysis of the hemodynamic response in fMRI studies. Since the recording is a slow indirect physiological consequence of neural activity, its stereotyped shape allows the many samples of a trial to be collapsed onto a single number representing the amplitude of that waveform in each trial, pooling the samples within a trial the way averaging pools across the trials themselves [15–17]. Even so, the hemodynamic response varies enough between subjects and between brain regions that estimating it for each subject outperforms assuming it [18, 19], and the assumed waveforms of other modalities vary between subjects as well [20–22]. Nearly as common is the *assumption of a shared sensor layout*. A regular imaging lattice or a conventional montage places neighboring positions on overlapping tissue, and the same position on the same anatomy in every subject, allowing evidence to be pooled across positions and subjects. This pooling strategy is standard in fMRI and in group-level EEG and MEG analysis [23–29]. However, functional roles assigned to specific anatomical structures vary across individuals [30, 31]. Consequently, spatial pooling accommodates this functional heterogeneity only by coarsening the resolution of an effect to the broader pooled region [32, 33]. These compensating assumptions involve inherent trade-offs, and empirical evidence suggests that an analytical framework minimizing such dependencies achieves superior sensitivity and spatial accuracy.

Here we focus on evoked responses in the form of cortico-cortical evoked potentials (CCEPs), recorded intracranially in patients with medically intractable epilepsy via stereoelectroencephalog-raphy (SEEG). A CCEP is classically the response evoked at one cortical site by an electrical pulse delivered at another site [34–37], and we use the term in the broader sense defined in Lyu et al. [38], for the response evoked at any site in the brain. In SEEG it is recorded from depth electrodes implanted to localize the seizure onset zone in patients with drug-resistant epilepsy [7]. CCEPs map the directed connections of an individual brain across distributed networks within a single session, with high spatial and temporal precision [37]. The intracranial contacts record the response directly and locally [7, 39–42], yielding a signal-to-noise ratio that, at the most strongly connected sites, is high enough for individual trials to be evaluated independently [43, 44]. CCEPs, however, have significant limitations. The recording is direct, so no slow indirect readout, such as the hemodynamic response of fMRI, imposes a stereotyped shape on the response [7], and as such the CCEP waveform varies with the stimulating and recording pair as well as with the patient’s age, state, and pathology [44–52]. The electrode locations are chosen based on clinical care, so no two patients share a layout and positions can be matched across patients only at the coarse level of an anatomical parcel [7, 52]. Within a patient, much of the signal that neighboring contacts on one shaft share is a volume-conducted copy of the same response [7, 53, 54]. The response also varies from trial to trial in amplitude and latency [43, 44], and later responses appear on only a subset of trials [55, 56]. In any case, the number of trials that can be collected is typically limited, because pulses are spaced about a few seconds apart to avoid altering the measured excitability [37, 57] and only a small amount of time is allotted for research during the patient’s hospital stay [7]. For CCEPs, then, each of the three aforementioned assumptions holds only loosely, if at all.

Despite this, the field’s approaches to detecting CCEPs rely on these same assumptions. According to a recent systematic review of 187 CCEP studies [58], the majority of those that report a detection method simply threshold the trial average inside a window fixed in advance [34]. Trial averaging makes the invariance assumption, and the window makes a weaker form of the known-waveform assumption because it specifies only the interval in which a response is expected, an interval the field has not even agreed on [58]. The other approaches in use gain statistical power by pooling across recording sites or by learning from human labels. Approaches that pool across recording sites combine them into one statistic through a model of how channels relate [23–28, 38, 59], and for CCEPs such a model would have to assume the connectivity the experiment set out to measure, beyond the volume conduction between neighboring contacts. These approaches can detect a response in a set of sites collectively but limit what can be inferred about which of the sites respond. Supervised detectors learn from human labels [60–62], and for one such detector, four raters judged the trial average by eye inside a fixed early window and agreed at a Cohen’s kappa near 0.6 [63]. A response outside that window is therefore never labeled, and the detector inherits the trial average, the window and the raters’ disagreement. Neither of these two approaches returns a measurement of the response at a single contact, since each makes the detection at a contact depend on more than that contact’s own recording: pooling on other contacts’ recordings, and a supervised detector on labels made from other recordings. SafeLens was designed to return a measurement of the response at each contact from that contact’s own recording, so the comparisons made below are with the approaches that do the same.

The approaches for CCEP analysis that return a measurement at each stimulating and recording pair assume a canonical response shape. They differ in how much trial-to-trial variation they allow. The trial average allows none, and canonical response parameterization (CRP) allows the amplitude to vary and nothing else [46]. It learns one shape and its duration per stimulating and recording pair from that pair’s own trials and so returns a waveform with a duration at each pair [46], and as the most developed of the methods that do, it is the primary benchmark here. Its authors state that one shape scaled by one amplitude per trial cannot represent the timing of individual features, so a component that shifts or disappears from trial to trial independently of the others has no parameter in CRP to represent it [46]. Models of per-trial timing give each component its own latency on each trial, so a component that shifts can be represented [14, 64–67], and some also give it its own amplitude [14, 67]. But these models require the number of components and their approximate positions to be specified in advance [11, 66, 67]. Standard practice typically extracts the number and positions of components directly from the trial average. As a consequence, these models inevitably inherit the very biases they aim to eliminate. No current method estimates the number and positions of components under rigorous statistical control.

Because every approach above either restricts its search to a preset window or derives its representation from the trial average, reported CCEP durations may reflect the sensitivity limits of those methods rather than the true end of the response. This limitation is greatest for later CCEP components, which shift in latency or appear on only a subset of trials [55, 56]. Trial averaging attenuates these deflections, and the single trial-invariant waveform of CRP cannot represent them [46], so duration estimates from either method can end too early. For instance, in a study applying CRP across the first second after stimulation, the averaged waveforms of significant responses lasted up to about 500 ms [68], and the paper introducing CRP showed example trial averages from five stimulation sites in two patients lasting up to 600–700 ms [46]. Delayed responses occurring up to 1 s after stimulation have been observed on single trials in seizure-generating cortex [55, 56], but whether CCEP activity outside such cortex persists at these latencies has never been statistically established. Determining where a CCEP actually ends therefore requires a detector that remains sensitive to late, trial-varying deflections across the full post-stimulus epoch.

Without assuming a waveform or a window in advance, no available method returns for each CCEP deflection a statistically controlled decision that it is present, when it began and ended, and with what polarity. SafeLens is designed to return that decision. By training random forests across a sliding window to discriminate single trials from interleaved rest and calibrating their feature reliance on separate resting recordings, SafeLens searches the whole epoch without requiring human labels, channel-pooling models, or assumptions of trial invariance. Because deflection-level decisions can only be verified where ground truth is known, we first validated SafeLens on simulated responses injected into real resting intracranial recordings. While holding its false-positive rate within the range expected at the nominal 5% level (5.4%), SafeLens resolved more than twice as many deflections as trial-average thresholding [69] or canonical response parameterization [46], with two-to fivefold lower mean boundary error and five-to elevenfold fewer false components. That advantage held under strong trial-to-trial variability, and the resulting map provided a data-driven prior that improved single-trial latency tracking over published analysis windows. Applied to CCEPs from eight patients, SafeLens resolved response structure extending beyond one second post-stimulus, whereas either benchmark recovered roughly half a second on the same data. Together, this work demonstrates that this framework for calibrating nonlinear classifiers on resting brain activity accurately recovers weak, trial-varying neural dynamics under family-wise error control, revealing that evoked cortical responses persist approximately twice as long as conventional analyses have been able to detect.

## M. Methodological Framework

### M.0 Overview

SafeLens takes an event that recurs at known times and a recording that supplies one time series per measurement site, and analyzes each site on its own. A site’s result depends on no other site and on no other subject. The output for each site is the SafeLens significance map, which describes the site’s response as deflections, each localized in time and assigned a frequency band and a polarity. The SafeLens significance map is thresholded so that the probability of reporting even one false deflection anywhere in a site’s significance map is held at a fixed level, 5% throughout this paper.

The recording is supplied as blocks (Section M.1). Each block is a continuous segment that contains the onset times of the events of one type recorded in it, and the resting data are supplied as blocks of the same form. From these blocks the analysis runs as one automated chain through to the SafeLens significance map (Fig 1). The event-locked and resting recordings enter as labeled examples of the two classes, event and rest (A). A searchlight window of fixed length slides across the peri-event epoch (C). The signal in each window is then converted into features that each report a rise or fall at one time in one frequency band (D). A random forest is trained at each searchlight position to distinguish the event windows from the resting windows (E), and every later stage operates on the trained forests. Ablation measures how much each forest relies on each feature (F, G), and a linear model condenses the ablation measurements into the integrated reliance surface, reliance over time and frequency (H). A deconvolution removes the smearing across time and bands that the searchlight window and the pyramid impose (I), and thresholding the deconvolved reliance surface combined over blocks yields the SafeLens significance map (J, M). The threshold on the deconvolved reliance surface comes from permutations of the same chain run on the secondary recordings with faux event times (B).

**Fig 1.**
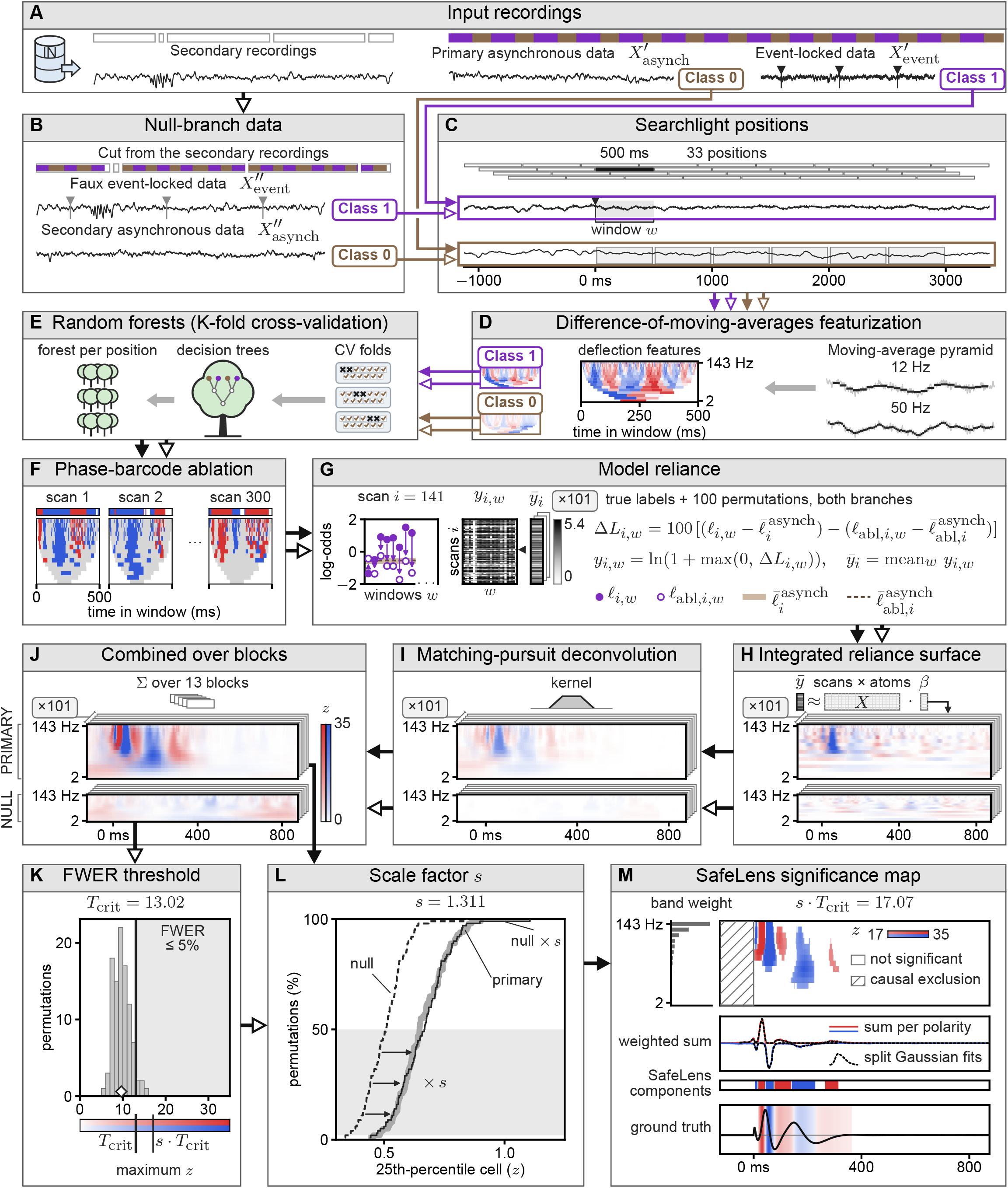
Overview of the SafeLens pipeline. All panels show the example channel of Fig 3A. **A**. The input is a recording in which blocks of event-locked data (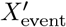, violet tiles) are interlaced with resting data called the primary asynchronous data (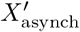, brown tiles), together with separate resting data, the secondary recordings. A block is one segment of event-locked data together with the adjacent primary asynchronous data. Violet lines represent class 1 and brown lines class 0. The black line from (A) to (B) represents the secondary recordings, and from the forests onward, one black line per branch represents that branch’s results. Arrowheads are filled in the primary branch and hollow in the null branch. Black triangles on the traces mark event times. **B**. The null branch replicates the blocks of the primary branch with faux event-locked data 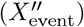 and secondary asynchronous data 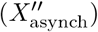. Each block’s faux event times (gray triangles, Section M.9) are the relative event times of the same block in the primary branch. Each box above the secondary recordings, in (A) and in the strip of (B), is one recording, and the unfilled box segments of (B) correspond to data that was not used. **C**. At each searchlight position across the peri-event epoch, one window w is extracted from every event-locked trial (class 1). The asynchronous data are segmented into asynchronous trials, as many per block as there are events, and each of their windows (gray boxes on the class 0 trace) is used as a class 0 sample at every position (Section M.3). The bars above the traces are the searchlight positions, and the black bar and the shaded window mark the one, starting at 0 ms, that (D to G) show. **D**. Difference-of-moving-averages featurization converts each window into deflection features that each report a rise (red) or a fall (blue) at one time in one frequency band, by a logarithmic pyramid of moving averages on a centered grid followed by first-order temporal differencing (Section M.3). The large image shows the features of the class 1 window, the small images each class’s mean features, and on the right two example bands’ moving averages (black) over the window (gray). A band is named by its frequency, 1 divided by its features’ support width. The white regions in the feature images of (D) and (F) mark times at which a band has no feature. **E**. A class-balanced random forest is trained at each searchlight position within a K-fold cross-validation whose test sets consist of whole blocks (Sections M.2 and M.5). Each card is one fold, with a cross on each held-out block. **F**. On the held-out windows of each fold, phase-barcode ablation measures how much each forest relies on its features (Section M.6). Each scan is defined by a signed barcode over time, red and blue bars with white gaps between them, and flips the sign of the features fully under bars of their own sign, and of partly covered features probabilistically. Three scans at one position are shown, each with its barcode above the window’s features, the flipped features in red or blue to highlight their polarity and the others in gray. **G**. Model reliance (Section M.6). For one scan i, the left plot shows the forest’s class 1 log-odds *ℓ* for class 1 windows before the ablation (filled) and after it (hollow), each arrow pointing from before to after, with the mean over the class 0 windows before (wide light line) and after (dashed), here nearly equal. ΔL is a window’s drop beyond the drop of the class 0 mean, and the model reliance y is zero wherever ΔL ≤ 0. The matrix shows y for every scan, with scan i’s row marked by the triangle beside it. One gray scale, saturating at the 99th percentile of y, colors both y and 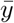. The same ×101 sheets are drawn behind every surface in (H to J), with the class labels permuted within each block. (G to I) show one block, from the held-out windows of its fold (Section M.2). **H**. A regularized general linear model (GLM) explains each scan’s 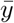 by the atoms (band × time × polarity) that the scan ablated, pooling the scans of every searchlight position onto one time axis. *β*, averaged over 50 bags of resampled trials and arranged over time and band, is the integrated reliance surface 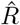 (Section M.7), drawn after the mean of its permuted surfaces is subtracted (Section M.8). X and *β* are drawn as schematics. The primary rows of (H to J) show the true labels and the null rows one permutation. The two rows of (H) share one color scale, in arbitrary units, saturating at the 99.7th percentile of the primary row, and in (H to J) and (M) blue is drawn over red where both are nonzero. **I**. The surfaces of both branches are standardized on the null branch’s permutations and then deconvolved by matching pursuit, which removes the temporal smearing of the pyramid (the kernel drawn), to yield each block’s deconvolved reliance surface (Section M.8). **J**. The deconvolved reliance surfaces are combined over blocks into one surface per channel (Section M.2). The color bar of (J), in z, also colors (I). **K**. The histogram counts the maxima of the null branch’s 100 permuted surfaces over the cells from 0 ms on, and *T*_crit_ is set so that the family-wise error rate (FWER) over those cells is at most 5% (Section M.9). The diamond marks the null branch’s unpermuted maximum, which is excluded. The strip shows the color scale of (J) on the same axis, with *T*_crit_ and, from (L), s *T*_crit_ marked. **L**. The scale factor s compares the permuted surfaces of the primary branch with those of the null branch (the arrow from (K), with *T*_crit_), each permuted surface summarized by its 25th-percentile cell (Section M.9). The curves show the cumulative share of the 100 permutations, the primary branch’s drawn as the wide gray line and the null branch’s, scaled by s, as the thin line on it. s is the median ratio of the primary curve to the null curve over the shaded lower half, and is at least 1. **M**. The SafeLens significance map is the primary branch’s combined surface (the line from (J) through (L)), thresholded at s·*T*_crit_ from 0 ms on. The hatched cells before 0 ms are the causal exclusion (Section M.8). The component read-out weights each polarity’s combined surface by the band weights shown and sums it over bands into the weighted sum, both in arbitrary units (Section M.10). Contiguous time points of one polarity that are significant in the map form a component, whose edges are trimmed by its split Gaussian fit or shared with a neighbor across a short gap. The ground truth, the channel’s injected CCEP averaged over trials, is highlighted over each deflection that SafeLens detected, more deeply where the curve is steeper.

Beyond the recording, the user supplies the event onset times, the resting data and the searchlight window length. The event onset times are needed because every window is placed relative to them. Enough resting data must be recorded that a portion can be set aside from the event-versus-rest comparison to build the null (Section M.1 and M.9). Rest recorded close in time to the events, ideally interleaved with them, serves the comparison best. The set-aside portion may come from farther away in the session. The searchlight window length *ω* is chosen to span the longest deflection expected, within the bound that the inter-event interval sets (500 ms throughout this paper, Section M.3). Every other setting, from the pyramid’s width schedule to the fold and permutation counts, has a reference default that was applied unchanged to every dataset reported here. S3 Table lists the remaining constants of the reference configuration.

The SafeLens significance map of a site is built from every trial of the event, in every experimental condition, pooled against the site’s resting record. The forests learn the response from every trial and the absence of a response from the resting record, which contains far more windows than the trials, and the sensitivity to weak deflections comes from that contrast. The SafeLens significance map can then inform the analyses that compare conditions, as a data-driven prior for per-trial and windowed methods (Section R.6). A direct contrast between two conditions within the same permutation machinery is possible in principle and is discussed in Section D.3.

On a workstation with 32 cores (AMD Threadripper), 128 GB of host memory and a 24 GB GPU (RTX 3090), a montage of 100 to 200 sites takes about 9 to 18 hours at the reference settings (S4 Table). Forest training accounts for about a quarter of that time and runs in parallel across the cores, so its speed follows the core count. The ablation and the solves for the permuted integrated reliance surfaces run on the GPU, and host memory sets how many sites one card processes at once. Sites are processed independently, so on a cluster the throughput grows with the number of GPU nodes. The host memory a site needs is set by the evidence tensor of the solve for the integrated reliance surface, which grows with the ablation scans, the searchlight positions, the trials of each block and the permutations. At the settings used here the analysis of one site peaks near 10 GB of host memory, though its raw recording is under 100 MB, and S5 Table gives the expression from which to scale that peak. Run time scales with the number of events and with the number of searchlight positions the inter-event interval admits, and permutations add memory far more than time (S5 Table).

### M.1 Data Specifications and Preparatory Procedures

SafeLens requires two categories of time-series data. The first is event-locked data 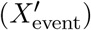, recordings with accurate onset markers of discrete events such as electrical-stimulation pulses, sensory presentations or task cues. The second is asynchronous resting data, recorded during periods with no events of interest, and it forms the rest class. Enough resting data must be acquired for two subsets. The primary asynchronous data 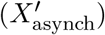 are compared directly against the event-locked data in the main analysis, and the secondary recordings are reserved for constructing the empirical null, from which the faux event-locked data 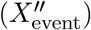 and the secondary asynchronous data 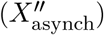 are cut (Section M.9).

A block is a set of trials that the user regards as part of the same condition, supplied as one recording segment with the list of their onsets (Fig 1A). In the datasets reported here, each block is one recording run of about 26 pulses delivered at one site and one intensity. The analysis holds out whole blocks, pools the trials within each and combines across them at the end (Sections M.2, M.8 and M.9). A difference among the trials of one block therefore goes unresolved, and the block’s estimate is formed from those trials alone. The smallest block tested contained 13 trials. Nothing requires the trials of a block to be consecutive events of the recording, and one run may be supplied as multiple blocks.

An event type is one kind of event delivered under one protocol, such as a pulse at one stimulation site, a tone or a task cue. Pulses at two different sites are two event types, and so are a go cue and a no-go cue in the same task. The events of a condition are repeats of a single event type, and SafeLens tests whether that event type is distinguishable from rest. A condition may span any number of blocks.

A block has one list of onsets and no label per event, so two event types recorded in one segment are supplied as two blocks (the same segment twice, once with each type’s onsets). Each type is then a condition of its own with its own SafeLens significance map, as stimulation from two sites yields two significance maps at one recording contact. SafeLens is designed for responses that have ended before the next event begins, since the searchlight positions of an event run from about a second before its onset to the following onset (Section M.3). It was not tested on responses that overlap in time. Two event types interleaved so closely that the response to one has not ended when the other begins are therefore not separated, since the windows of each type contain both responses. Separating them requires the types to be spaced so that one type’s response has returned to baseline before the other’s windows begin. Two event types whose responses are expected to share a morphology can equally be kept as two conditions of one event type and compared as such (Section D.3), for example when a change of state shifts only the response’s latency.

Each block has an acquisition timestamp, by which the blocks are ordered chronologically. The cross-validation assigns whole blocks to folds in recording order (Section M.2), and the null construction copies each block’s onset pattern into the secondary recordings in chronological order (Section M.9). Inter-event intervals are measured within a block and never across a block boundary, so a recording cut to excise artifact, epileptiform periods or other events is admissible when every cut falls at a block boundary.

Certain preprocessing steps are recommended before SafeLens is applied. The released analysis pipeline applies no referencing, filtering or artifact substitution of its own and operates on the signal as supplied, so these steps are performed beforehand. Re-referencing is strongly advised [7], and the choice among common-average referencing, a bipolar derivation and a Laplacian derivation depends on what the analysis treats as noise and what as signal [7]. For the stereo-EEG data reported here we used a one-dimensional Laplacian, which references each contact to the mean of its two neighbors on the electrode and localizes CCEPs to gray matter better than the referential or common-average montage does [53]. The Laplacian was chosen over the bipolar derivation because SafeLens reports a signed deflection at each contact and the sign of a bipolar derivation depends on the order of subtraction [7], and it is the reference recommended for local population-level and low-frequency oscillatory activity in stereo-EEG [54]. The choice of referencing matters to the result, since the re-referenced signal is what the pipeline analyzes and the polarity of every deflection in the SafeLens significance map is read from it (Section V.2).

A known stereotyped artifact, such as the brief high-amplitude stimulation artifact 1 to 15 ms after a pulse in CCEP recordings, may be remediated before feature extraction and need not be. The simulation results reported here were run with the generator’s stimulation artifact left in place, an alpha-function pulse of 1 ms rise and 10 ms duration (Section V.1). SafeLens localizes the artifact as a component like any other, and because each tree sees only a subset of the features, the reliance on the response survives alongside the reliance on the artifact (Section D.2). We did not test a transient that overlaps a component of interest in time, so whether the two would be resolved as separate components is unknown. We also did not test whether removing the artifact first would improve the results. Contaminated samples were not substituted with interpolated or synthetic values, since substitution can introduce structure that differs from rest. A user who does want to remediate can run the analysis once to locate the contaminated interval, treat that interval with the tool of their choice and run the analysis again.

Explicit correction for slow baseline drift by high-pass filtering is generally unnecessary, since the differencing features of every frequency band are blind to a constant offset and robust to slow trends (Section M.3). On one simulated dataset, we tested a simple per-trial detrend and found that the number of deflections SafeLens resolved was unchanged (493 of 846 with and without it, a paired difference of 0 percentage points with a 95% confidence interval of −1.4 to 1.4). So, we avoided it, since drift correction can itself introduce artifacts [70].

### M.2 Cross-Validation Architecture

The classifier is trained and scored inside a K-fold cross-validation over recording blocks, to prevent overfitting (Fig 1E). The simulated validation and the real-data application both used seven folds over all of a subject’s blocks (thirteen or fourteen in the simulated data, eleven to fourteen in the real data), each fold holding out one block or two consecutive blocks.

Folds may follow a seeded random partition of the blocks, contiguous runs of them in recording order, or an explicit list of blocks given per fold. The analyses reported here used contiguous runs. The resting data follow the same cut, one resting block beside each event-locked block (Section M.1). In each iteration the blocks of one fold are held out, and the rest form the training pool. Everything the classifier learns is learned on that pool, from the scaling applied to each feature to the trees of the ensemble and the score threshold that separates the two classes (Sections M.4 and M.5). The ensemble then scores the held-out windows, and every event-locked trial is scored once, by the fold that held out its block. Every resting window is kept, and the trees’ bootstrap draws equal numbers from the two classes (Section M.4).

The remaining stages, from the ablation to the deconvolution and its null, then run on each block’s held-out trials (Sections M.6 to M.9). Each block b therefore yields a deconvolved reliance surface 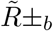 of its own, and the blocks are brought together only at the last step, where their surfaces are summed and divided by the square root of the number of blocks B (Section M.8). Which blocks are summed together is the user’s choice, and it is made after the forests have been trained. A condition is one such grouping, so the folds may cut across the conditions and one set of forests serves them all. Each condition is read from its own held-out trials, scored by forests that trained on the rest of the blocks, of that condition and of the others. Regrouping the blocks changes only that last step, so one run yields a SafeLens significance map for every grouping the user forms. Each grouping is a test of its own, with its null combined by the same rule. The threshold read from that null turns the combined deconvolved reliance surface 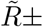 into that grouping’s SafeLens significance map (Section M.9). The held-out trials are also the evaluation set on which SafeLens, CRP and the z-scored detector are compared (Section V.3).

### M.3 Temporal Windowing and Difference-of-Moving-Averages Featurization

Within each cross-validation iteration, a sliding window of fixed duration *ω* (500 ms at the 1 kHz sampling rate, so 500 samples) is analyzed at every position of a grid across the epoch (Fig 1C). Event-locked positions advance in steps of Δ_event_ = round(*ω*/4) = 125 ms, so every time in the interior of the epoch lies within four windows. The grid begins a full window plus one window- and-a-step before the event, at −*ω* − (*ω* + Δ_event_) = 1125 ms, so that every pre-event time from −750 ms to just before −375 ms lies within four windows, none of which reaches the event. It ends where a full window would reach the mean inter-event interval, so that no window straddles the next event. The pre-event positions are trained and read like every other position, and their reliance enters the model of the integrated reliance surface with the rest (Section M.7). The read-out thereby sees positions with no response at the same coverage as the positions with one. The period before the event is neither subtracted as a baseline nor assumed to be quiet. The reported SafeLens significance map begins at the event onset (the causal mask in Section M.8), which is the interval on which the benchmark methods are scored (Section V.3).

The widest level of the pyramid described below is bounded by the window, so the window length sets the longest deflection the method can represent. The window *ω* is therefore chosen to span the longest deflection expected, and the inter-event interval only caps that choice. A deflection is one leg of an excursion that must also return before the next event, so no deflection can exceed half the interval, and a design should keep the deflections it expects well inside that bound. The reference window of 500 ms is a seventh of the 3.4 s interval used here. A fixed window also keeps each forest’s feature count bounded however long the epoch, so training grows with the number of positions and never with the epoch length. The step trades resolution for computation. A quarter window separates the components of a cortico-cortical evoked potential, whose durations run from tens to a few hundred milliseconds, and a smaller step yields more forests per time point and a finer estimate at proportionally greater cost.

Time is resolved at two levels. Within a window, every feature has its own time and frequency band, and the ablation in Section M.6 reads which of them the forest relies on, so reliance is placed on the features’ own grid, finer than the window. That placement has a limit, since a forest can substitute one feature for a correlated neighbor, so two times inside a window are separated only as far as the forest treats their features as distinct. Across windows there is no such limit. A forest trained at one position has never seen the signal outside its window, so two times more than a window apart are read from forests whose training windows never overlapped, and their evidence is independent by construction. Each interior time is read from the four forests covering it, and the model of the integrated reliance surface in Section M.7 admits reliance only where those positions agree.

Resting windows are drawn on a grid of their own, which begins at the start of each asynchronous trial of the primary asynchronous data (Section M.9) and advances by Δ_rest_ = *ω* = 500 ms, so that consecutive resting windows do not overlap. Every resting window serves as a comparison sample at every event-locked position, so resting samples already far outnumber event samples, and overlapping resting windows would add only correlated copies of them. A window that would overrun a recording edge is discarded, and a channel that cannot supply ten windows per class is skipped.

For each window, a feature vector is computed from a difference-of-moving-averages pyramid (Fig 1D). The signal is smoothed by boxcar moving averages of increasing width, and each smoothed level is then differenced in time, which estimates a derivative at that level [71, 72]. We call the resulting features deflection features, since each reports the local rise or fall of the signal in one frequency band. Level k of the pyramid uses filter length *W*_*k*_ and step *S*_*k*_, fixed by a logarithmic schedule. Starting from a base length *W*_base_, successive lengths grow geometrically by a factor *γ*, with each increment capped, snapped to a multiple of the base length, and floored to advance by at least one base length,

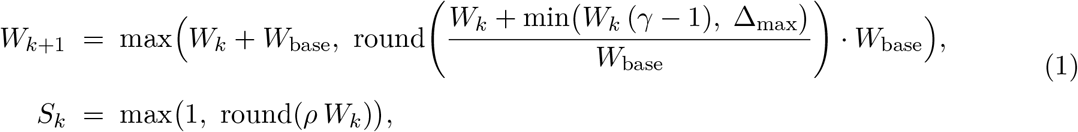

with reference values *W*_base_ = 5 samples, growth factor *γ* = 1.33, per-step cap Δ_max_ = 50 samples and step fraction *ρ* = 0.33. The pyramid grows geometrically while the increment *W*_*k*_(*γ* − 1) remains below Δ_max_ (up to about 150 samples) and linearly in steps of Δ_max_ beyond that. The linear regime keeps the widest levels sampled densely enough to cover the middle bands. The narrowest width is the sampling limit, and the widest is bounded by the window, whose length is set by the longest deflection expected. Adjacent bands overlap, so their features are correlated, and the forests use that redundancy. Each tree is grown on a third of the features (Section M.5), so a deflection that one tree cannot see in one band it finds in the next. The ensemble’s reliance on a deflection therefore spreads over neighboring bands, in a shape the read-out knows and inverts (Section M.8). A larger growth factor spaces the bands more sparsely and costs less, and a smaller one covers more finely at a cost that rises with the window width.

Because *W*_*k*_ and *S*_*k*_ need not tile the window evenly, each level’s grid is offset inward by half the residual *R*_*k*_ = mod(*ω* − *W*_*k*_, *S*_*k*_), which centers its sampled positions within the window. Every level then has a support width *d*_*k*_ = *W*_*k*_ + *S*_*k*_, the width that the ablation, the timestamp of the integrated reliance surface, the steady-state bound and the deconvolution operator all use. The centered moving average at level k is

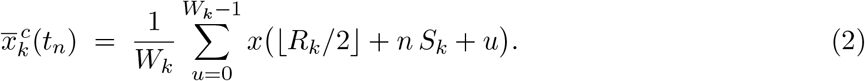

and the feature is its first-order temporal difference,

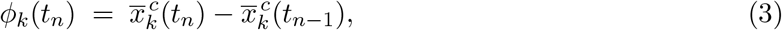

where the *t*_*n*_ are spaced by the step *S*_*k*_. The difference is a local slope, so the features are blind to any constant offset and robust to baseline shift and to variation in absolute amplitude, and the concatenation of {*ϕ*_*k*_} over all levels is the window’s feature vector. Because the averages are centered, a sharp excursion after the onset enters the wide levels up to half a box width before its own time. The causal mask in Section M.8 removes from the SafeLens significance map the part of that backward leakage that falls before the event, and the guard is not complete for extreme signals (Section D.6).

Two implementation choices make the dense featurization tractable. First, every level’s box average is obtained in O(1) per output sample from a single O(L) prefix sum over the windowed signal, the integral-image construction of computer vision [73]. The whole pyramid therefore costs one cumulative sum and no convolution. Second, the featurization runs on the GPU. Wavelet transforms [74] and FFT-based filtering can also separate a signal into bands, but a wavelet analysis of a great many overlapping windows is computationally burdensome and FFT-based filtering is more susceptible to the leakage and edge artifacts of short segments [75]. A moving average is a finite-impulse-response low-pass filter, and differencing a smoothed signal multiplies that low-pass response by a high-pass one. Each level of the pyramid is therefore a crude band-pass filter whose center frequency falls as the box widens, the one-dimensional analogue of a difference-of-lowpass image pyramid [71]. The ladder of widths tiles frequency, and the integrated reliance surface is indexed by time and frequency accordingly, each level k named by the frequency *f*_*k*_ = 1000/*d*_*k*_ of its support width *d*_*k*_ in ms. These frequencies fall from 143 Hz at level 1 to 2.2 Hz at level 15. The pyramid is fixed in advance and identical for every fold and every label permutation, so the feature basis contains no information about the labels of the data it is tested on [76]. Featurization is applied after windowing, so that no feature at a position draws on a sample outside that position’s window (Section M.8).

### M.4 Feature Standardization and Class Balancing

The ensemble is trained on the full feature vector of every window, with no dimensionality reduction and no feature selection. At the reference settings, that vector comprises the 769 deflection features of the pyramid (Section M.3), the Class 0 and Class 1 feature maps of Fig 1D. Two operations prepare the features for the ensemble.

First, the features are standardized per searchlight position. The mean and standard deviation of every feature are learned from the fold’s training pool and applied to the held-out windows. A feature that is constant across the training pool has its standard deviation set to one, so that nothing divides by zero.

Second, the classes are balanced at training time. Resting windows typically far outnumber event-locked windows at any searchlight position, and a forest trained on that imbalance tends to favor the resting class and to classify the event-locked class poorly [77]. Each tree is therefore grown on a balanced bootstrap, which is the draw of the balanced random forest of Chen, Liaw and Breiman [77]. The bootstrap draws, with replacement, as many event-locked windows as the tree’s training partition contains and an equal number of resting windows. Each tree draws a different resample of the resting pool, so the ensemble learns the resting class from far more windows than any one tree sees. The forest learns from the trials what an event-locked window looks like, and from the far larger resting record it learns what one does not look like. A misclassification-cost matrix whose off-diagonal entries are the two bag sizes is also supplied to every tree. At the one-to-one ratio used throughout, its two costs are equal, so the matrix reduces to the standard loss and the balancing is performed by the bootstrap alone.

### M.5 Random-Forest Classification and Per-Trial Scoring

A random forest [78] is trained for each searchlight position of each fold (Fig 1E). The forest consists of 300 trees, the reference count applied to every dataset reported here (Section M.0). Each tree is grown on a class-balanced bootstrap of the training pool’s windows (Section M.4), sees a random subset of the features, and has one of the pool’s blocks set aside from its draw. The trees therefore differ in their trials, their features and their blocks. No single block dominates the aggregated vote. The block exclusions are dealt evenly and systematically over the pool. Each event-locked trial has its own realization of the background activity and its own amplitude. A score averaged over trees grown on different resamples of trials and of blocks therefore reflects the response common to the trials and is robust to the particular noise draw and amplitude of any one of them. Leakage from the fold’s held-out blocks is already precluded by the fold partition (Section M.2).

The subset each tree sees is a third of the features, drawn once per tree (256 of the 769), a random subspace draw that prevents the trees from growing alike and is suited to redundant feature sets [79]. At each split, 28 of the tree’s visible features are tried as candidates, the square root of the full feature count and the usual default for a classification forest [80, 81]. Splits use the Gini criterion [82]. Growth stops at a minimum leaf size of one twentieth of the event-locked windows in the tree’s training partition (max(1, round(0.05 *n*_event_))). The fraction sets the floor at a single window when trials are few and at several when trials number in the hundreds, so a large training set cannot be split down to single-trial leaves. The field’s default of one observation per leaf grows a tree to full depth [80], and the larger minimum here limits the depth to which a tree fits its bootstrap. Neither the third of the features nor the leaf floor is expected to bear much on the result, and the reasoning behind both choices and the alternatives is given above and in Section D.2. Each tree is compacted as soon as it is trained, to cap peak memory, and the searchlight positions are trained in parallel.

For every searchlight window, the forest returns a continuous score, the average over all its trees of the posterior probability of the event-locked class. Every later stage reads this score (Sections M.6 to M.9), and it is recorded for every held-out trial at every searchlight position (Section M.11).

Two products of this stage are kept for the stages that follow. The first is the trained forest, flattened from tree objects into arrays of split variables, split points, child pointers, and leaf probabilities of the event-locked class. A per-tree offset table indexes the arrays, and a context table keyed by searchlight position and set-aside block stores the retained feature indices and the standardization constants. The flattening reduces storage, and because the reliance interrogation re-scores the forest hundreds of times per channel (Section M.6), it lets every tree be descended in one vectorized pass. A saved forest is therefore self-contained for re-scoring. The second product is the score of every held-out event-locked trial at every searchlight position, recorded together with its trial identifier and label, its searchlight position and its fold. These scores are the leakage-free record that the reliance and significance analyses read (Sections M.6 to M.9), and can be used for single-trial analysis (Section M.11). Because every pool block has trees that never drew on it, a decision threshold can be calibrated on out-of-block predictions at no extra training cost, and one is recorded with each forest, with accuracy and ranking measures beside it. These are capabilities beside the pipeline reported here, and no stage of it reads them. They give a quick check from training alone that the two classes separate on a channel, and a per-trial metric representing each trial’s likeness to the event-locked class (Section M.11 and D.3).

### M.6 Feature-Reliance Interrogation via Phase-Barcode Ablation

A random forest’s score at a searchlight position does not by itself reveal which features contain the event-related evidence or when within the searchlight window that evidence lies. As with the weight vector of a linear decoder [83], the forest’s internal parameters cannot be read directly as a feature-importance map. Because temporal localization is a primary objective, SafeLens adds a dedicated interrogation stage that measures the forest’s reliance on each feature by controlled ablation (Fig 1F, G). This interrogation belongs to the removal family of model explanations, in which a feature’s influence is measured by removing it and re-scoring the fitted model [84]. The quantity estimated is the fitted model’s reliance on the feature, and the reliance is the fall in the model’s performance when the feature’s information is withheld [85]. The interrogation’s nearest ancestor is the permutation importance of random forests, which breaks a variable’s association with the outcome and reads the loss in out-of-bag accuracy [78, 86, 87]. While Fisher, Rudin, and Dominici formally define model reliance as a ratio of expected losses, MR(f) := *e*_switch_(*f*)*/e*_orig_(*f*) [85], we adapt this framework to the groups of features that each ablation scan flips. SafeLens quantifies the model reliance on a group not as a loss ratio, but as the drop in log-odds separation that sign-flipping the group causes (Δ*L*_*b,i,w*_, Eq (6)). The integrated reliance surface (Section M.7) combines these group reliances into a reliance at each time and frequency. Occlusion probes apply the removal to a localized region of an input [88]. A SafeLens scan perturbs a temporally coherent, polarity-alternating region of the feature space by sign-flipping features within it, so the scan tests the direction of the reliance as well as its presence.

Each scan applies its own phase barcode to the searchlight window. The barcode is a row of contiguous bars, each of value +1, 0 or −1, and a bar of value 0 is a gap. The barcode’s generator emits a pair of bars of opposite polarity, and a gap follows each pair. Bar and gap widths are drawn log-uniformly between about half the width of the highest-frequency features and half the width of the lowest-frequency features. The uniform draw that sets each width is squared toward the narrow end, since the number of time-by-frequency locations at a width falls in inverse proportion to that width, and the scans together tile every location. A gap’s width is zero with probability 0.25, and the zero-width gap is the special case that lets a multiphasic pattern develop. A zero-width gap extends the alternation by a further pair of deflections, so the chance of a multiphasic pattern falls geometrically, fourfold with every added pair. A gap of nonzero width ends the pattern, and the next pattern begins with a fresh random polarity. The modal output is therefore a single biphasic pair between gaps, at a random phase. The alternation interrogates the rising and falling edges together, so the measured reliance has the biphasic form the deconvolution recovers (Section M.8). A neural response that depolarizes always rebounds, so the biphasic form is the response’s own shape. The features are correlated across time and across frequency bands, so an ablation drawn independently for every feature would waste most of its scans on removals in which the features left in place fill in for the features removed. The barcode biases the ablation toward removing temporally related features together.

On each of the 300 scans (the reference count, Section M.0), the barcode is read as two ablation masks over the searchlight window, a positive mask made of its positive bars and a negative mask made of its negative bars. For each feature *φ*, 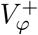 is the fraction of its time span in the positive mask, and 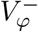 the fraction in the negative mask. Each feature’s own discriminative direction is summarized by a signed tree-vote consensus *ψ*_*φ*_ over a random sub-bag of trees. The consensus accumulates, over every split on that feature in the sub-bag, the difference in event-class probability between the split’s right and left children. A positive consensus therefore means that a larger value of the feature raises the event-class probability. The sub-bag is a random 63.2% of the forest, the fraction of a bootstrap’s draws that are unique (1 − *e*^−1^). On every scan the sub-bag is redrawn, so the votes’ own uncertainty is sampled across the scans. The polarity-matched fraction of feature 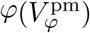 is then its fraction in the mask whose polarity matches its consensus,

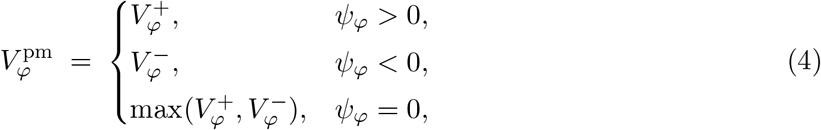

Ablation is enacted by sign-flipping a feature’s values in the fold’s held-out windows and rescoring the trained forest. The features are z-scored slopes, so negation turns a rise in one frequency band into a fall, setting the flipped feature’s own learned direction against it. The flips follow the polarity-matched fraction, so the positive mask selects only features the forest reads as rising, and the negative mask only features it reads as falling. Each scan therefore tests features of one learned direction at a time, and rising and falling deflections that overlap in time are told apart (Section M.7). A re-scoring is only a re-walk of the compact trees (Section M.5), so the many re-scorings are cheap. Feature *φ* is ablated on scan i with probability:

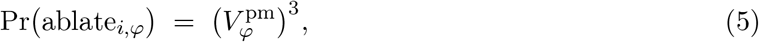

realized as an all-or-nothing Bernoulli draw for every feature on every scan. A feature with a polarity-matched fraction 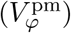 of 1 is ablated on every scan, one with 0.5 on about one scan in eight (0.5 cubed is 0.125), and one with 0.2 on almost none (0.2 cubed is 0.008). Grading the probability by the polarity-matched fraction lets the later steps resolve frequency band as well as time (Sections M.7 and M.8). The difficulty in resolving frequency band is that the barcode is one-dimensional, spanning time, so every bar that overlaps with a high-frequency feature also overlaps each lower-frequency feature whose time span is a superset of the high-frequency feature’s. This is why we decided against the simpler approach of ablating every feature with a polarity-matched fraction above zero (Pr(ablate *φ* on scan i) = 1 if 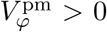), which would confound the ablation of a high-frequency feature with the ablation of its superset features. Instead, the cubic term in Eq (5) was chosen as a middle ground between the linear draw, 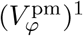, and the limit of higher exponents, 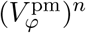 with n → ∞. A high-frequency feature has superset features in nearly every lower-frequency band, so a linear draw would ablate too many of them together with it to tell their effects apart, while higher exponents would ablate the lowest-frequency bands ever more rarely.

Throughout Sections M.6 to M.9, b = 1, …, B indexes the blocks of a recording (Section M.2), and every quantity before the combination of Section M.8 is computed separately for each block. Each is also computed in both branches, the primary branch and the null branch (subscript null). Every quantity is computed under the true labels and again under each of the *N*_perm_ label permutations of Section M.9. A superscript (j) marks the version under permutation j, and a symbol without it is computed with the true labels. t is the time from the event onset in ms, negative before it. It runs on the 5 ms knots *t*_*κ*_ of Section M.7 until the standardization places each surface on the 1 ms grid. f is the frequency that names a band, 1000 divided by its support width in ms, one of the 15 pyramid levels of Section M.3. A band’s support width is the time span of its features, the moving-average width plus the step. The superscript ± marks the rising or falling polarity, a hat an estimate and a tilde a deconvolved surface.

The model reliance of a scan is the reduction it induces in the forest’s class separation on the fold’s held-out windows. It is built from the effect of scan i’s ablation on each held-out event-locked trial w of block b, measured in the log-odds of the forest scores. The effect is the log-odds drop:

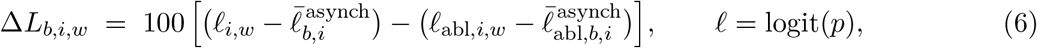

with *ℓ*_*i,w*_ and *ℓ*_abl,*i,w*_ the log-odds of trial w’s score before and after scan i’s ablation, and 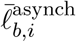 and 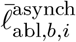 the mean log-odds over every window of block b’s held-out resting trials before and after it. Observation i is one ablation scan at one searchlight position, and w is a held-out event-locked trial of block b, scored through its window at that position. Every score is clipped to the interval from 10^−4^ to 1 10^−4^ before the logit. Δ*L*_*b,i,w*_ is the drop that the ablation causes in trial w’s log-odds separation from the resting windows. The factor 100 sets where the logarithm of the next equation begins to compress, at a drop of about 0.01 in log-odds separation. The model reliance *y*_*b,i,w*_ is Δ*L*_*b,i,w*_ floored at zero and compressed by a logarithm to stabilize heavy tails,

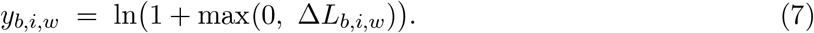

Over the 300 scans, the per-scan per-trial ablated scores record where and in which band the forest’s evidence lies, and the model of the integrated reliance surface reads them as model reliance (Section M.7). The whole procedure is run identically for the real models and for the null models (Section M.9).

### M.7 The General Linear Model of the Integrated Reliance Surface

The ablation scans (Section M.6) sample the forest’s reliance stochastically, and a general linear model (GLM) condenses them into the integrated reliance surface, the GLM’s coefficients over time and frequency band (Fig 1H). The GLM’s response variable is built from the effect of scan i’s ablation on each held-out event-locked trial w, measured in the log-odds of the forest scores. It is the model reliance *y*_*b,i,w*_ of Section M.6 (Eq (7)).

The design matrix has one column for each atom in each of two polarities, where an atom is one frequency band at one time knot and the time knots *t*_*κ*_ are points spaced 5 ms apart across the epoch. A feature belongs to the band of its pyramid level and to the time knot nearest the center of its support in the epoch (Section M.3). The feature’s polarity is positive when the forest at its searchlight position reads it as rising and negative when that forest reads it as falling, by the sign of its tree-vote consensus taken over all the forest’s trees (Section M.6). Each row i of the design is one scan at one searchlight position, so the atoms at a time knot are informed by the scans of every searchlight position whose window covers that knot.

Each tree’s vote on a feature is the tree-vote consensus (Section M.6) accumulated over that tree’s splits alone, and the trees’ votes are gathered onto the atoms, positive votes as rising vote mass and negative votes as falling vote mass. Each band’s vote mass is smoothed along time by a Gaussian with the standard deviation of a box as wide as the band’s support, and the grids *P*_pos_ and *P*_neg_ are the rising and falling vote mass averaged over the trees. Each feature enters *P*_pos_ or *P*_neg_ by its polarity, with the value of the grid at its atom. The design block is sign([A *P*_pos_, A *P*_neg_]), where A is the matrix of the per-scan ablation indicators *A*_*φ*_ ∈ {0, 1} (the realized Bernoulli draws of Eq (5)). An entry of the block is therefore 1 when the scan ablated at least one of the atom’s features of that polarity and the grid is nonzero at the atom, and 0 otherwise. A coefficient thus measures the response to the atom’s ablation, independent of how many of the atom’s features the scan ablated and of their vote mass. The design also has one nuisance column for each searchlight position and polarity, equal to 1 on the position’s scans with a 1 anywhere in that polarity’s half of the design block, to absorb offsets at the level of the searchlight position.

The coefficients are estimated by a bagged ridge regression [89, 90], in which each bag q solves the normal equations,

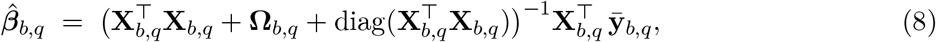

where *X*_*b*_ is the binary design of block b with the nuisance columns appended, *X*_*b,q*_ contains the columns of *X*_*b*_ that bag q solves over, 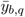 contains the mean of *y*_*b,i,w*_ over the bag’s drawn trials for each observation i, the third term is a baseline ridge equal to the diagonal of the Gram matrix (applied to all atoms, including the nuisance columns), and Ω_*b,q*_ is an epistemic penalty on the same columns that applies to the feature atoms alone and grows as the trees’ votes at an atom disagree,

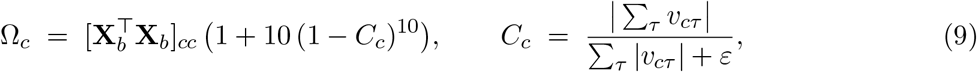

with *C*_*c*_ ∈ [0, 1] the tree-vote agreement at atom c, *v*_*cτ*_ the net vote at atom c of the *τ*-th tree of each forest (its rising vote mass less its falling vote mass, averaged over the forests that vote at the atom), and *ε* a small stabilizing constant. *C*_*c*_ is 1 when these votes share a sign, falls toward 0 as they cancel and is 0 at an atom without votes. The baseline ridge scales with the number of scans on which an atom was ablated, and the epistemic penalty [91] adds a second shrinkage set by the forest’s votes. Atoms on which the votes agree (*C*_*c*_ → 1) receive a diagonal loading of 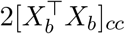, the baseline ridge plus the penalty’s floor. Atoms on which the votes cancel (*C*_*c*_ → 0) receive up to 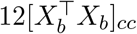 and are shrunk heavily. Because of the tenth power, the loading stays within 1% of its floor wherever *C*_*c*_ is at least 0.5, and the penalty grows appreciably only where the votes nearly cancel. The votes in *v*_*cτ*_ are pooled over every searchlight position whose window covers the atom’s time, so the penalty shrinks an atom on which the forests of those positions cancel one another’s votes.

We chose to stabilize the bagged ridge regression with 50 bags, in which bag q resamples *n*_*b*_ trials with replacement from block b’s *n*_*b*_ held-out event-locked trials to form 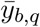 and draws a random 63.2% of the tree indices. The trial draw is shared by every observation, and the tree indices are the same at every searchlight position. A bag leaves out of its solve every atom at which the mean vote of the bag’s trees is below 0.01 in magnitude, and that atom’s coefficient in the bag is zero. Negative coefficients are set to zero within each bag, and the integrated reliance surface is the average of the coefficients over the 50 bags. Bags fitted in the same way to ln(1 + max(0, −Δ*L*_*b,i,w*_)), from ablations that raised an event-locked trial’s log-odds relative to rest, give a separate surface that no later stage reads.

The averaged coefficients form an integrated reliance surface for each polarity over time knot and frequency band. The rising surface 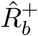 comes from the positive-polarity half of the design block and the falling surface 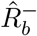 from the negative-polarity half. With 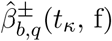 bag q’s coefficient for the atom at time knot *t*_*κ*_ and frequency f in the rising (+) or falling (−) half of the design, zero where the bag left the atom out,

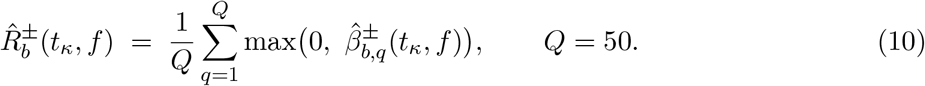

The integrated reliance surface is computed in two branches, the primary branch from the forests of Section M.5 and the null branch from the null forests of Section M.9, drawn as the primary and null rows of Fig 1H to J. Each branch’s surface is computed under its unpermuted labels and under 100 label permutations, and the forests are not retrained for a permutation. A permutation reassigns which of the branch’s held-out windows form the event-locked and resting sets of the GLM’s response variable. The null branch’s permuted surfaces form the null distribution of the significance test (Section M.9), and the permuted surfaces of both branches are used to match the two branches’ levels and spreads (Sections M.8 and M.9).

### M.8 Matching-Pursuit Deconvolution and the SafeLens Map

The integrated reliance surface 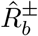 (Section M.7) is indexed by the time knots and frequency bands of the deflection features. Because each time point lies inside several moving-average boxes of every band (Section M.3), a deflection is averaged into features at several times and bands, and reliance on it appears smeared in both time and frequency. A cell-wise threshold read directly from the smeared integrated reliance surface would report the extent of a deflection as the extent of its smear. To recover the underlying response structure, SafeLens deconvolves the standardized integrated reliance surface by a matching pursuit [92] (Fig 1I).

The forests of the primary and null branches are trained against resting data from different recordings (Section M.1), so the two branches’ integrated reliance surfaces can differ in level and spread. The centering and standardization below and a factor s applied at the threshold (Section M.9) match the two branches’ surfaces, so that a threshold taken from the null branch applies to the primary branch’s deconvolved reliance surface. For centering, the integrated reliance surfaces of each branch are centered at each time knot on that branch’s own label permutations, the mean of its permuted surfaces being subtracted from its unpermuted surface and the mean of the other permuted surfaces from each permuted surface. Centering removes what all of a branch’s surfaces share regardless of the labels, which comes from that branch’s forests and resting recording, so the unpermuted surface measures the change that the true labels make. Negative centered values are set to zero. Each band is then expanded from its time knots onto the 1 ms sample grid of the data by convolution with a Gaussian whose standard deviation is that of a uniform distribution over the band’s support width. The Gaussian has peak 1 and is truncated to that width. For standardization, the rising and falling polarities of each band are expressed as z-scores relative to the band’s mean and standard deviation in the null branch’s permuted surfaces. Both statistics are taken over the interior of the epoch and averaged over the polarities, with a percentile-based noise floor on the null standard deviation. The z-scores are half-wave rectified, so that the standardized integrated reliance surface the pursuit operates on is a non-negative measure of reliance in excess of what the branch’s permuted labels produce on average,

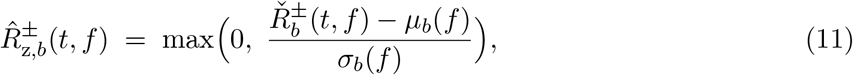

where 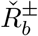 is 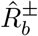 after the centering, the zeroing of negative values and the expansion. The statistics *µ*_*b*_(*f*) and *σ*_*b*_(*f*) are the mean and standard deviation of band f over the null branch’s permuted surfaces 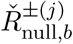, with *σ*_*b*_(*f*) floored at the 10th percentile over bands of the standard deviations above 0.1.

The pursuit deconvolves the standardized integrated reliance surface deterministically. Because every band of the moving-average pyramid is a box average, the smearing the pyramid imposes is known in advance and serves as a kernel with no free parameters. Reliance generated at source band k is smeared into observation band k^′^ at time lag Δt in proportion to the fraction of band k^′^’s support that falls inside k’s support when the supports are centered Δt apart,

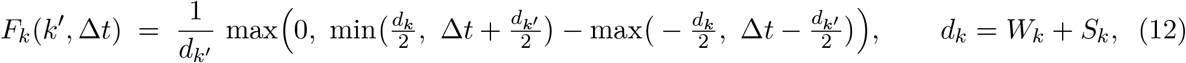

with *d*_*k*_ and 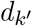 the support widths of the source and observation bands. Each *F*_*k*_ is the moving-average pyramid’s point-spread function for source band k. The kernel is determined entirely by the support widths, which are set as described in Section M.3.

Under this kernel, the pursuit iteratively identifies biphasic pairs of lobes, each lobe the reliance on one deflection. For each candidate lead lobe, the pursuit seeks a partner of opposite polarity in the same band, within a phase-shift window scaled to the band’s own timescale (0.5 to 1.5 support widths after the lead lobe’s time). Pairing lobes of opposite polarity within the phase-shift window is the assumption that the deconvolution makes about the response’s shape. The pairing does not fix the number of deflections, their latencies, or their amplitudes, and a response of three deflections is recovered as two pairs sharing the middle lobe.

The kernel is split into causal and anti-causal halves so that a lead lobe is deconvolved with the causal half and its partner with the anti-causal half. In the causal half, the rows *F*_*k*_(*k*^′^, Δ*t*) of the bands k^′^ other than k are zero before the lobe’s time (Δt < 0), and in the anti-causal half after the lobe’s time (Δt > 0). Each lobe thus explains reliance in the other bands only toward its partner, and a lead lobe explains none in those bands before its own time, since reliance explained there would make the lead lobe’s deflection appear to begin early. The row of band k, being the lobe itself, remains symmetric about Δt = 0 in both halves.

The pursuit starts from the standardized integrated reliance surface as its residual surface. Each iteration scores every band, time and polarity of the residual surface as a candidate lead lobe with the causal half of that band’s kernel and as a candidate partner with the anti-causal half. Each score is the residual surface’s cross-correlation with the half, divided by the norm of the half. Each lead lobe is paired with the highest-scoring candidate partner of opposite polarity within the lead lobe’s phase-shift window. Of all these pairs, the one whose two scores have the largest harmonic mean is selected. Because the harmonic mean is limited by the weaker score, the two lobes constrain each other (Section D.2), and a lead lobe whose candidate partners all score zero scores zero as a pair. The selected pair’s two halves of the kernel are then each fitted to the residual surface by least squares. Both halves are subtracted from the residual surface at one shrunk amplitude, which a soft-threshold shrinkage sets to the harmonic mean of the two fitted amplitudes less 0.25, and the residual surface is then clamped to be non-negative. The pursuit stops when the selected pair’s shrunk amplitude or the harmonic mean of its scores drops below 0.1, or after 100 iterations.

The deconvolved reliance surfaces 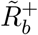 and 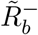 of block b come from the pair of standardized surfaces 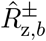 in three steps. The pursuit’s output is the portion of the standardized integrated reliance surface that the selected pairs explain, namely the standardized integrated reliance surface less its unexplained residual. A common-mode rejection then simultaneously subtracts each polarity’s pursuit output from the other, both subtractions using the pre-update values, and clamps the results at zero. Where both polarities’ pursuit outputs are positive at one band and time, only the excess of the stronger polarity remains (Section D.5). An asymmetric cross-band smoothing then spreads reliance three bands toward lower frequencies but only one toward higher, by a truncated Gaussian with a standard deviation of one band. Every step from the expansion onto the 1 ms grid through the cross-band smoothing is applied identically to the primary branch’s surfaces and to the null branch’s surfaces (Section M.9).

The smoothed deconvolved reliance surfaces 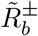 of a condition’s blocks are combined (Section M.2), and the combined deconvolved reliance surface of each polarity is denoted 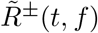,

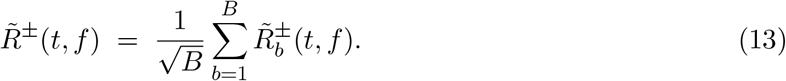

The same sum, with the same B, combines the null branch’s surfaces and each permutation of either branch. Permutation j of a combined surface sums each block’s own permutation j, which the blocks draw independently. The factor 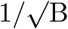 is Stouffer’s, and it cancels in the test, since *T*_crit_ and s come from surfaces combined by the same sum. We impose a causal prior on the SafeLens significance map, that no part of the response begins before the event. The prior is applied as a causal support mask *ℒ*(*t, f*), zero in every band before the event onset and one from the onset on. The cells at which *ℒ* is zero are the causal exclusion (Fig 1M). Because each band’s moving averages are centered (Section M.3), a sample shortly after the onset still averages data from before it. Section D.6 discusses the backward leakage of the centered averages that remains after the onset. The combined surface is thresholded against s · *T*_crit_, where *T*_crit_ is the critical value of the permutation test and the factor s ≥ 1 matches *T*_crit_ to the scale of the primary branch’s combined surface (Section M.9). The thresholded surface is multiplied by the causal support mask, giving the SafeLens significance map (Fig 1M),

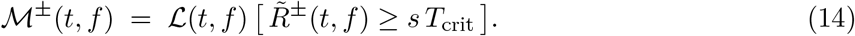

The bracket [P] is 1 where P holds and 0 elsewhere. For each polarity, collapsing *ℳ*^±^ across bands by a logical OR yields the one-dimensional set of time points at which SafeLens declares a polarity-specific event-related response.

### M.9 Null Construction and Family-Wise Error Control

Each polarity’s deconvolved reliance surface 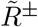 is tested at every time and frequency that the causal mask keeps (Eq (14)), and the family-wise error rate across those tests is controlled by a permutation test on the maximum of 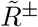 [23, 93]. The maximum runs over the same cells, every time and frequency of the epoch’s interior from the event onset on. The null distribution of that maximum is built from the null branch’s resting data, the faux event-locked data 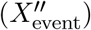 and the secondary asynchronous data 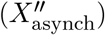, cut from the secondary recordings (Section M.1) to replicate the event-locked blocks and the blocks of primary asynchronous data alike. The chain that produced 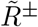 is run on the null branch’s data from forest training to the GLM, where their event-locked and resting labels are permuted, so no forest is retrained under a permutation. The surface of every such permutation is deconvolved and adds one maximum to the null distribution, which therefore needs no analytic form [23]. The family-wise control holds provided that, under the global null hypothesis *H*_0_ that the channel has no event-locked response, the observed maximum of 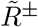 is exchangeable with the null maxima once the two branches share a scale (below and Section D.6).

Together the faux event-locked data 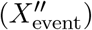 and the secondary asynchronous data 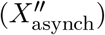 replicate the event-locked data and the primary asynchronous data block by block, and each of their blocks contains one trial for each trial of the block it replicates. The resting blocks between the event-locked blocks (the primary asynchronous data, 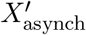, Fig 1A) have no events and so are cut into asynchronous trials, whose starts also begin the grids of the resting windows drawn from those blocks (Section M.3). The starts of the asynchronous trials of each resting block form a train whose intervals are drawn with replacement from the inter-event intervals of all the event-locked blocks. Drawn intervals are added until the next interval would make the train’s span exceed the block’s length minus 2c, where the clearance c is the smallest inter-event interval of the event-locked data, and the train is then centered in the block. In every dataset reported here the trains together contain more starts than the event-locked data have events, and starts are removed one at a time until the asynchronous trials number the same as the events. Each removal deletes the first or last start of the train in the resting block with the most starts and moves no other start, so the trimmed train is never re-centered and its intervals remain drawn intervals. The asynchronous trials are placed once for each subject, so the forests of every channel share them. With every block’s event times and trial starts in place, the blocks of 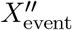 and 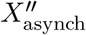 are cut in the chronological order of the blocks they replicate, in one forward pass through the secondary recordings, so no sample is used twice. The event times copied onto a block of 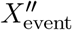 are its faux event times (Fig 1B). Each block is cut as one unbroken segment that spans its event times or trial starts at the spacing of the block it replicates, from c before the first to c after the last. A block that does not fit in what remains of the secondary recording being cut is cut whole from the next recording that can contain it, and it is shortened only if no later recording can.

Null forests trained on the faux event-locked data and the secondary asynchronous data are passed through the same ablation, GLM and deconvolution stages as the forests trained on the event-locked and primary asynchronous data. The GLM evaluates each branch, primary and null, on its true labels and under *N*_perm_ label permutations (reference 100). Each permutation exchanges labels only among the trials of each pair of an event-locked block and its resting block (Section M.2), so that labels are exchanged only between trials recorded close together in time. The null branch’s permutations give the null distribution of the maximum, and the primary branch’s permutations are used only to center that branch’s surfaces (Section M.8) and to compute the scale factor s (below). The null branch’s evaluation on its true labels is retained outside the null distribution as a record of the resting-state background the pipeline produces on resting iEEG.

The test statistic is the maximum of 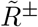 over both polarities and the tested cells *T* (every time and frequency of the epoch’s interior at which *ℒ*(*t, f*) = 1), so the null maxima and the observed maximum are taken over the same cells. Since each polarity’s surface is non-negative (Section M.8), one upper threshold protects both polarities against false positives. Each permutation contributes one such maximum, taken on the null branch’s permuted surface 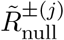 combined over blocks as 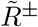 is (Eq (13)), giving an empirical null distribution {*M*_*j*_} (j = 1 to *N*_perm_). The threshold *T*_crit_ is the null maximum of ascending rank ⌈(*N*_perm_ + 1)(1 − *α*)⌉, which at a family-wise error rate *α* = 0.05 and *N*_perm_ = 100 is the 96th null maximum,

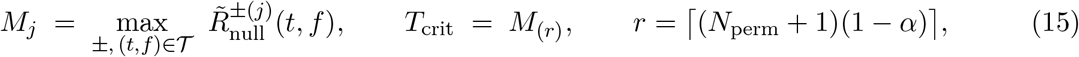

where 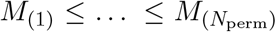 are the null maxima in ascending order. The + 1 in the rank counts the observed maximum of 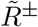 as one of *N*_perm_ + 1 maxima, so under *H*_0_ the probability that 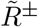 meets or exceeds *T*_crit_ anywhere in *T* is at most *α* [94].

This bound holds only if the primary branch’s deconvolved reliance surfaces 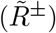 are on the null branch’s scale, since *T*_crit_ is taken from the null branch alone. Centering matches the two branches’ levels, and standardization divides both by the null branch’s spread, which puts them in the units of *T*_crit_ (Section M.8). A difference in spread, however, survives standardization, so if the primary branch’s integrated reliance surfaces 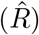 vary more widely than the null branch’s, its standardized 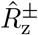 surfaces, and with them its 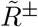 surfaces, are larger. The two branches’ forests are trained against resting data from different recordings (Section M.1), so the spread of their 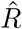 surfaces can differ. The primary branch’s 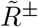 surface can then cross *T*_crit_ more often than *α* under *H*_0_. On each channel *T*_crit_ is therefore multiplied by a factor s, the scale ratio of the primary branch’s 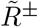 surfaces to the null branch’s, and a location is declared significant where 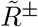 meets or exceeds s ·*T*_crit_ (Eq (14)). s is estimated from the two branches’ permuted surfaces, since under *H*_0_ each branch’s unpermuted surface is exchangeable with its permuted surfaces, which therefore show how large the branch’s surface is under *H*_0_.

A permutation changes only which trials of a block have the event-locked label, keeping their number, so about half of the block’s event-locked trials keep the label on average, and more or fewer in any one permutation. Where there is an event-related response, a permuted surface of the primary branch retains the difference between the mean response of the trials labeled event-locked and that of the trials labeled resting, which averages zero over the permutations but can be large in a single one. The permuted surfaces that contain most of this difference would enlarge s, and with it the threshold. To exclude the response from s, each permuted surface is summarized by its 25th-percentile cell over all bands and time points, taking the larger polarity at each cell, which excludes the surface’s peaks. These time points include those before the event onset, which the permutation test excludes (Eq (15)), so that s is estimated from more baseline than the tested cells alone would provide. s is the scale ratio of the two branches’ distributions of these summaries, estimated over their lower halves as the median ratio of their quantiles, primary over null, at the 2nd, 4th and so on to the 50th percentile. In estimating s from the parts of the surfaces that the response rarely reaches, we took an approach akin to genomic control, which estimates the inflation of association statistics from their median because few of them are non-null [95]. The factor is floored at 1, so that sampling noise in its estimate cannot lower the threshold below *T*_crit_ [96]. To test how the results depend on the summary percentile, we repeated the analysis with summary percentiles from the 10th to the 100th, and each held the false-positive rate on resting iEEG with no CCEP injected at or below nominal, while higher percentiles, which take in more of the response, resolved fewer deflections.

### M.10 Component Read-Out

The SafeLens significance map of Eq (14) is the final output of the core analytical framework described in this work. However, the validation in Section V.2 scores discrete components, each with a polarity and boundaries. We therefore describe here one last post-processing step that we call the component read-out, a heuristic that we use to collapse the SafeLens map into a definite number of such components. A user may instead analyze the time-by-frequency significance map directly or collapse it in another way, and Section D.5 discusses the limits of this specific read-out. The read-out starts from the significant time points of each polarity (Section M.8), and a time point significant in both polarities is assigned the polarity of the highest-frequency band significant at that time. Where that band is significant in both polarities, the time point is assigned the polarity with the larger weighted sum. A polarity’s weighted sum is its deconvolved reliance surface 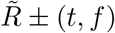 summed over frequency bands with each band weighted by the inverse of its support width, so that most weight goes to the highest-frequency bands, whose narrow support widths place a deflection most precisely in time. Each weight is also multiplied by the central difference of the logarithm of the support width at that band, which corrects the sum for the uneven spacing of the bands’ support widths on a logarithmic scale (Section M.3). Contiguous time points of one polarity form a component. To prevent over-fragmentation, components of one polarity are merged across gaps shorter than the 7 ms support width of the highest-frequency band (Section M.3), and components shorter than 2 ms (half the 5 ms knot spacing of Section M.7, rounded down) are dropped.

Because a component’s significance can begin and end away from the edges of its deflection, the read-out then refines each component with curves fitted to its polarity’s weighted sum. A component is first cut in two at the dip in its weighted sum between two deflections of one polarity, when the finest band significant at the dip is coarser than the finest bands significant on either side of it. Each component is then fitted with a split Gaussian (a Gaussian with separate widths before and after its peak). Up to five more split Gaussians are then added to the component’s curve, each where the weighted sum exceeds both that curve and the other polarity’s weighted sum by the largest area, as long as that area is larger than the 95th percentile of the areas by which the same polarity’s weighted sum exceeds the other’s in the null branch’s evaluation on its true labels (Section M.9). Two neighboring components, other than the two parts of a cut, share one boundary when the gap between them is shorter than half the support width of the wider of their dominant bands (a component’s dominant band is the band of its largest significant value), at the crossing of their curves if their polarities differ and at the gap’s middle if they do not. Every other edge moves to where the first split Gaussian’s cumulative area reaches 5% at an onset or 95% at an offset, within limits set by its neighbors and its significant bands. The remaining details of this heuristic, including its handling of edge cases, are in the script safelens_readout.m of the public SafeLens repository (Data and Code Availability).

### M.11 Per-Trial Scores and Single-Trial Analysis

The significance reported in this work is controlled family-wise over the times, frequencies and polarities of each channel’s deconvolved reliance surface 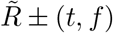, which pools the channel’s trials (Section M.9). No per-trial test is made. The SafeLens significance map nonetheless serves single-trial analysis by supplying the number of components, their spans and their polarities, which together form a prior that per-trial timing models may use. Section V.6 tests the SafeLens significance map as that prior against the common approach of fixed analysis windows and against CRP’s components, and Section R.6 reports the results. Single-trial analysis can also use the per-trial scores, the classifier score of every event-locked trial at every searchlight position (Section M.5). The per-trial scores are free of train-test leakage because everything the classifier learns, from the scaling of each feature to the trees, is learned without the scored trial’s block (Sections M.2 and M.4). A per-trial significance test on these scores is left to future work (Section D.3).

## V. Validation Design

### V.1 Simulated CCEPs and Ground Truth

Because ground truth is unknowable in empirical recordings, we validated SafeLens on simulated CCEPs injected into real resting iEEG, so that the background noise, autocorrelation, and non-stationarity were those of genuine brain activity while the event-locked response was known exactly. The injection was additive, each channel’s signal being its real resting trace plus a noise-free evoked template generated for that channel after each simulated pulse, so the resting trace supplied the noise against which SNR was defined and the template was the ground truth. The templates were injected into one resting recording of each of two subjects (168 and 184 channels, the two channel sets of the simulated-data figures). In each recording, event blocks of about 26 pulses at a fixed interval of 3.4 s alternated with resting blocks, as recommended for real data (Section M.0). The resting blocks supplied the primary asynchronous data for the event-versus-rest comparison, and the subjects’ other resting recordings supplied the secondary recordings for the null (Section M.1). The secondary recordings were further from the ideal of rest recorded close to the events (Section M.0), since they were recorded 3 to 7 hours after the injected recording, after clinical CCEP mapping and around a session of theta-burst stimulation (TBS), both of which can change resting activity, so the null data need not match the primary asynchronous data. The empirical null nonetheless held the false-positive rate on these data within the range expected at the nominal 5% (5.4%, Sections R.1 and D.6).

Each channel’s evoked template was generated by its own instance of David et al.’s neural-mass model of evoked responses [97], a Jansen–Rit cortical column [98] that received an extrinsic drive after each pulse [99]. The column had three populations: pyramidal cells, excitatory interneurons and inhibitory interneurons. The template was the pyramidal cells’ membrane potential, the excitatory minus the inhibitory postsynaptic potential they received. Published values set the connection strengths within the column and from the drive into it [97, 99] and the gain of the inhibitory synapses [97]. The gain of the excitatory synapses was drawn uniformly for each channel between 3.25 and 4 mV, from a stable to a near-critical column.

We adapted the model to CCEPs in four ways. First, the extrinsic drive was a train of input volleys of mostly alternating sign that widened up to 1 s, timed so that successive input peaks fell across the latency ranges of the early (N1), intermediate (positive) and later (N2) deflections characteristic of CCEPs. The train’s strength, which scaled every volley, was the lower of a rising and a falling curve. The rising curve was logistic, with its half height about 4 ms after the pulse plus 1 ms for every millimeter between the column and the stimulation site, a distance drawn for each channel. The falling curve decayed exponentially from the pulse to 1% of its starting value at a decay time drawn for each channel between 0.5 and 2 s, where it ended the train. Second, the pulse’s direct electrical effect was modeled as an alpha function peaking at 1 ms and cut at 10 ms, whose amplitude fell with the inverse square of the same distance. The alpha function drove the postsynaptic potentials of all three populations without passing through a synaptic gain. This input put a stimulation artifact at the start of every template, as the pulse does in a CCEP recording. The artifact was part of the ground truth but was not scored as a response, since Section V.2 excludes the first 15 ms after the pulse from the deflection and channel scores (Section M.1 describes how SafeLens handles the artifact). Third, the synaptic time constants rose within each trial so that early transmission was fast and late transmission slow, the excitatory constant from an AMPA-like to an NMDA-like regime and the inhibitory constant from a GABAA-like to a GABAB-like regime. Fourth, the extrinsic drive into each population was split into a part added directly and a part multiplied by the pyramidal cells’ firing. The multiplied part grew from 5% to 35% of the drive in step with the rise of the time constants, shifting the drive from direct input toward modulation of the column’s own activity.

Trial-to-trial variability was manipulated explicitly by jittering the generative parameters. Each recording received templates in a zero-jitter and a high-jitter condition, which gave four datasets and allowed SafeLens, CRP and the z-scored detector to be compared between datasets with and without that variability (Section R.5). A subject’s two datasets shared the same resting background, channels and pulse times, but each channel’s parameters were drawn independently in the two conditions from the same ranges, so the conditions were compared as unpaired samples of templates. In the high-jitter condition the excitatory synaptic gain, the decay time of the drive, the timing of every volley and each time constant’s starting value, final value and rise duration were jittered on every trial. Each volley’s timing was set from the volley before it, so the timing changes accumulated along the train and the later deflections varied most from trial to trial. The jitter level was set as high as it could be while the later deflections still repeated across trials, since at higher levels the timing of successive deflections would overlap so much from trial to trial that the deflections lost their boundaries. Each pulse’s template was drawn from 500 trials simulated for its channel, and the resulting latency jitter of the high-jitter condition was measured on those 500 trials. On each trial, each deflection of the trial-averaged template (Section V.2), a rise or a fall, was matched to the rise or fall of the same kind whose steepest point lay nearest the deflection’s own steepest point in the average. The deflection’s latency jitter was the standard deviation of that point’s time across trials, as a percentage of the deflection’s duration. Its median was 10% (interquartile range 7–14%) for deflections starting before 100 ms, 14% (10–19%) at 100 to 250 ms, 38% (22–60%) at 250 to 500 ms and 64% (43–95%) at 500 to 1500 ms (S6 Table). For 849 of the 1,347 adjacent pairs of deflections starting after 250 ms, the two deflections’ latencies overlapped across trials, the later one’s 5th percentile falling before the earlier one’s 95th.

Across channels, template amplitude was swept over a wide range of signal-to-noise ratios, so that sensitivity could be characterized as a function of SNR. Within each variability condition, all 352 channels of the two recordings were ranked together by the SNR of their unscaled templates and assigned in rank order the 352 target SNRs of a grid evenly spaced in decibels from −20 to +20 dB. Each channel’s template was then scaled by one amplitude multiplier that brought its SNR to its target. A template’s SNR was the mean of the decibel values of its single-trial SNRs. A trial’s SNR was the RMS of that trial’s template over a window centered on the largest absolute peak of the trial-averaged template, divided by the RMS of that trial’s detrended resting trace over a window twice as wide with the same center. On each trial the scaled template was then divided into lobes at its zero crossings, and each lobe was set to zero (a noise floor) unless it and one of its neighboring lobes reached a single-trial SNR of −50 dB over their joint span. These single-trial SNRs differ from the SNRs by which Figs 2 to 4 bin channels and deflections, which are taken on trial averages (the next paragraph and Section V.2) and can lie outside the −20 to +20 dB range. Each channel’s ground-truth response window was read from the local SNR of its template against the resting trace it was added to, both averaged over all the held-out trials of the cross-validation (Sections M.2 and V.3). The local SNR at each time after the pulse was the largest of the segmental SNRs [100], the ratios in decibels of the template’s standard deviation to the resting trace’s, over windows of 5 to 500 ms ending at that time. The response window ran from the first to the last time after 15 ms at which the local SNR exceeded −15 dB. Its largest local SNR was the channel SNR by which Fig 2 bins the channels.

**Fig 2.**
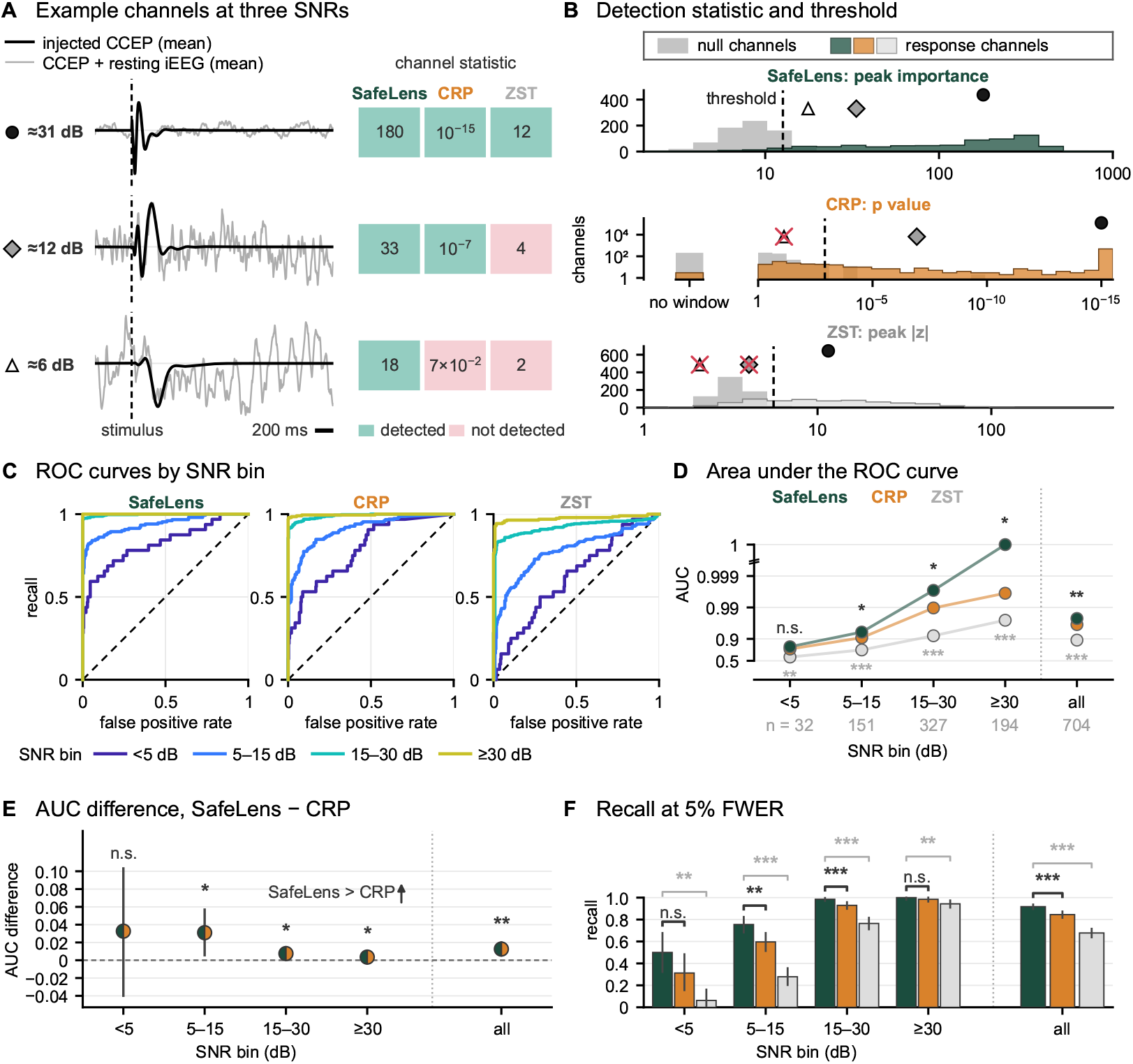
Channel-level detection of simulated CCEPs by SafeLens, CRP and ZST. Pooled over the 704 channels of the four simulated datasets (the 352 recorded channels of two subjects, each under the zero-jitter and high-jitter conditions), with the channel as the unit of analysis. **A**. The traces of three example channels, each with its channel statistic (Section V.2) from SafeLens, canonical response parameterization (CRP) and the z-scored trial-averaging detector (ZST), the statistics plotted in (B). Each cell is green where its method detects the channel at the threshold used in (F) and pink where it does not. **B**. All three methods’ statistics on the channels with an injected CCEP (response channels) and on the same channels’ resting iEEG with no CCEP injected (null channels), with mixed colors where the two overlap. Symbols mark the example channels of (A), with a red cross where the method does not detect the channel. CRP’s and ZST’s dashed lines are their thresholds pooled over the two subjects. CRP’s p values are floored at 10^−15^, with the channels for which CRP returns no response window in a separate bar. **C**. Receiver-operating characteristic (ROC) curves of SafeLens, CRP and ZST, one per channel SNR bin. Each curve is computed from the bin’s response channels and all null channels. Dashed diagonals mark chance. **D**. Area under the ROC curve (AUC) per SNR bin and over all channels, with the number of response channels (n) under each bin. **E**. The AUC difference, SafeLens minus CRP, per SNR bin and over all channels, with its 95% interval. Values above zero, where the arrow points, mean that SafeLens had the higher AUC. **F**. Recall at the 5% family-wise error rate (FWER) per SNR bin and over all channels, each method at its own threshold (Sections M.9 and V.3). The unit of tests and intervals is the recorded channel (both datasets). Intervals are bootstraps over recorded channels, in (E) with their null channels and all three methods drawn together. Stars in (D) test SafeLens against CRP (black, above the points) and against ZST (gray, below them) by the bootstrap of (E), the black ones repeated in (E), and brackets in (F) compare pairs by an exact sign-flip test of each one’s recall difference. The tests of (D) and of (F) are each Holm-corrected over the ten comparisons of SafeLens with CRP and with ZST that the panel draws (* p < 0.05, ** p < 0.01, *** p < 0.001, n.s. not significant). Gray marks involve ZST.

### V.2 Detection and Resolution Metrics

SafeLens, CRP and the z-scored detector were scored against the noise-free templates (Section V.1) at two levels, the templates’ individual deflections and the whole channel. Each channel’s noise-free templates, which varied from trial to trial in the high-jitter condition, were averaged over all the held-out trials of the cross-validation (Section V.1). This averaged template was split into deflections where the slope of its 5 ms moving average changed sign. Each deflection was thus a single rise or fall, and its polarity was positive for a rise and negative for a fall. Each deflection’s component SNR was the ratio in decibels of the averaged template’s RMS over the deflection to the RMS over the same span of the residual noise, the average of the same trials’ simulated recordings minus the averaged template. Deflections starting in the first 15 ms after the pulse, the post-stimulus exclusion band that contains the stimulation artifact, were not scored. They were left out for a fair comparison with CRP, which begins its analysis 15 ms after the pulse to limit contamination by the artifact [46]. To check that leaving them out did not favor SafeLens, we also scored them in the four simulated datasets and found that SafeLens resolved 66% of them against 51% of the later deflections, so leaving them out lowered its recall.

A method’s significant region was the set of time points at which it reported a significant response, and the method resolved a deflection when its significant region covered at least half of the deflection’s variation, the total absolute change of the averaged template g across the deflection. Covering a steep segment thus counted for more than covering a flat one, and for deflection i the criterion was

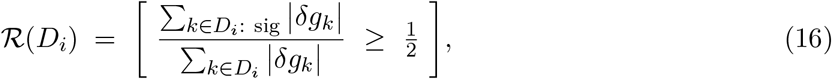

where *D*_*i*_ is the set of k for which samples k − 1 and k both lie in deflection i, *δg*_*k*_ = *g*_*k*_ − *g*_*k*−1_, and sig restricts the sum to the changes with at least one of their two samples in the significant region. Resolving a deflection also required matching its polarity. Only SafeLens reports the polarity of each component it detects (Section M.10), so only components correctly matching the deflection’s polarity were counted as detections. For CRP, whose significant region was the segment it marks as significant (between *τ* B and *τ* R, Section V.3), this requirement was far more lenient. A straight line fitted to CRP’s canonical response shape across the deflection only had to slope in the deflection’s direction. The z-scored detector was held to the same polarity requirement as SafeLens, with the polarity of each of its components read from the order of the peaks of its z-score (Section V.3).

Deflections with a component SNR at or below a floor of −15 dB were not scored, and a channel’s response ended at its first pair of consecutive deflections at or below the floor after its first deflection above it, every later deflection counting as below the floor. A deflection below this floor was nonetheless scored, for all three methods, when any one of them resolved it with a component of the deflection’s polarity that matched both of the deflection’s edges to within 25% of the deflection’s duration. The exception was made so that a clear detection below the floor counted for SafeLens or CRP and never against that method as a false positive (Section R.1). This exception added 20 deflections to the 3945 above the floor, and a method’s recall was the fraction of these 3965 deflections that it resolved.

The boundary error was a second deflection-level measure. It was computed for each method on the deflections it resolved, and each pair of methods was compared on the deflections both resolved (Fig 4 and S1 Fig). CRP’s components were the rising and falling parts of its canonical response shape within the segment it marks as significant (Section V.3). Each deflection a method resolved was paired with that method’s component of the same polarity that overlapped it longest, and an onset error and an offset error were measured between the pair’s edges. Each boundary between two adjacent deflections had a tolerance band, running from the point at which the earlier deflection had completed 97.5% of its variation to the point at which the later one had completed 2.5% of its own. Each error was zero when the component’s edge lay inside the band at the deflection’s corresponding edge, and otherwise was the distance from the component’s edge to the nearer end of the band, as a percentage of the deflection’s duration.

At the channel level, each method gave each channel two values, one on its response and one on rest. Each was the largest value of the statistic the method thresholded (Sections M.9 and V.3). The response value was computed from the channel’s held-out trials over the whole epoch after the post-stimulus exclusion band, for all three methods. The rest value was the largest value of the same statistic over the same epoch on the channel’s resting iEEG with no CCEP injected (Sections V.1 and R.1). Each recorded channel appears in both jitter conditions with the same resting iEEG, so its rest value appears twice among the 704 rest values. Channels were binned by channel SNR (Section V.1) in four bins (below 5, 5 to 15, 15 to 30, and 30 dB or more), and each bin’s receiver-operating characteristic (ROC) was computed from the response values of its channels against the rest values of all 704 channels. Each bin’s result is reported as its area under the curve and the fraction of its channels detected at the threshold each method uses throughout (Fig 2D and F). S2 Fig shows a component-level ROC computed on segments of each channel’s response.

False positives were counted in three further ways. First, each method was run at the threshold it used throughout (Sections M.9 and V.3) on the resting iEEG that underlies the simulated datasets, with no CCEP injected, and every channel on which it reported a response there counted as a false positive (Section R.1). These recordings shared no data with the secondary recordings on which each method’s threshold was calibrated, so each threshold was tested on data it had not been set on. Second, a simulated channel had a false positive when a response that a method reported there after the exclusion band overlapped no deflection of the averaged template at any SNR, including those in the exclusion band. For SafeLens the rate expected by chance is its rate in the first count (5.4%, 19 of 352 channels) scaled by the share of time points that no deflection covers. Deflections cover 33.5% of the time points after the exclusion band, averaged over the 704 simulated channels, so SafeLens’s expected rate is about 3.6% of channels (66.5% of 5.4%). Third, the components that each method reported on the simulated datasets were checked against every deflection of the averaged template at any SNR, including those in the exclusion band. A component was false if it overlapped with no deflection. It was also false if it overlapped a deflection starting after the exclusion band and the component’s declared polarity disagreed with the sign of the averaged template’s net change from the component’s onset to its offset. False components were counted as a percentage of the components each method reported (Fig 3C, Section R.3).

**Fig 3.**
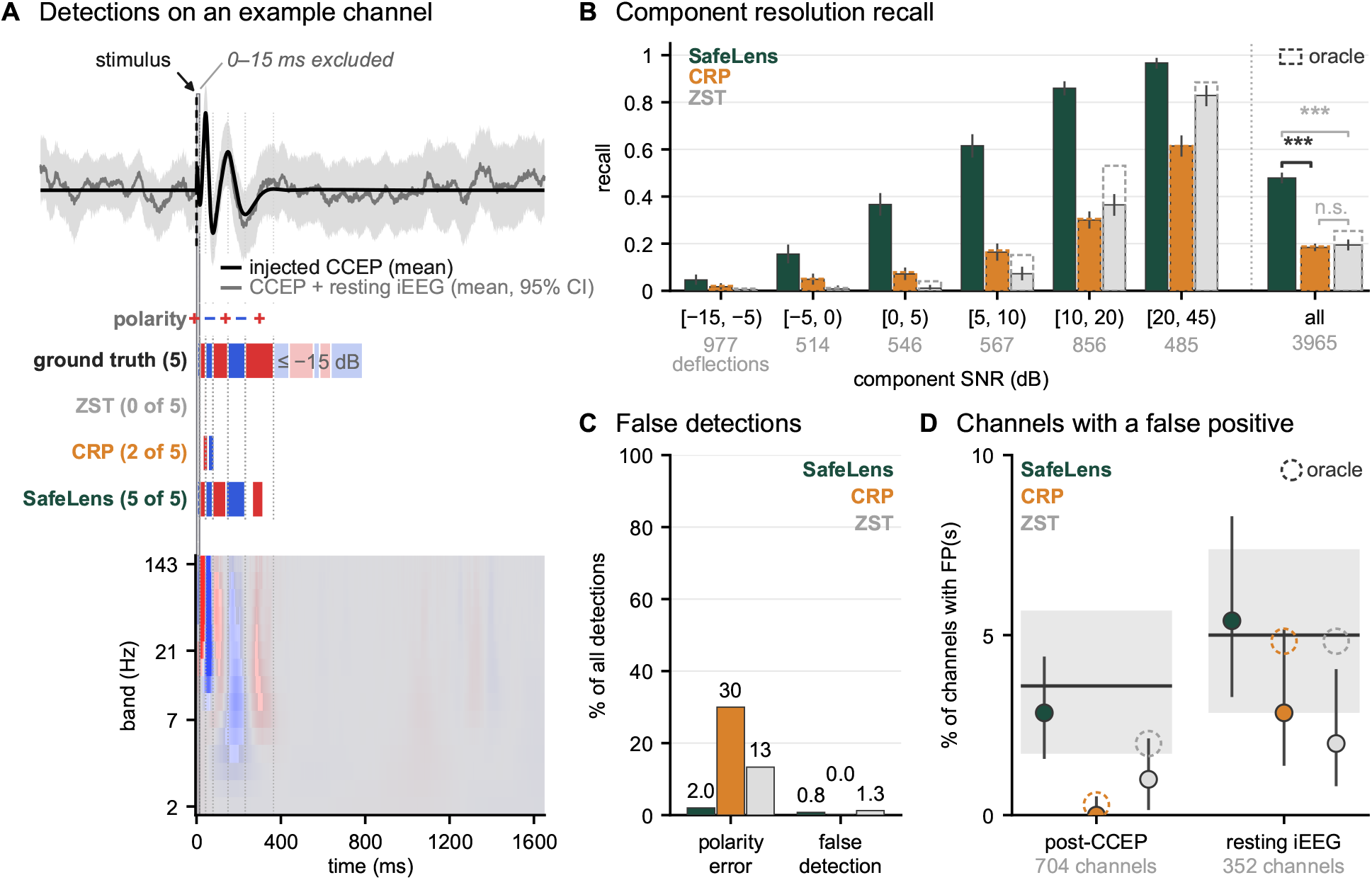
Component-level sensitivity and specificity of SafeLens, CRP and ZST. **A**. An example channel and the detections of SafeLens, canonical response parameterization (CRP) and the z-scored trial-averaging detector (ZST). From the top are the injected CCEP’s mean (the ground truth) and the recorded mean over all trials, baselined over 500 to 5 ms before the stimulus, with a simultaneous 95% confidence interval over all available time points (mean ± 3.68 SEM), the polarity of each evaluated ground-truth deflection, the ground-truth deflections (a response ends at its first pair of deflections at or below −15 dB, and those after it are faded and are evaluated, for all three methods, only where at least one of the three methods precisely identified both of a deflection’s edges, Section V.2), and each method’s detections, red for positive (rising) and blue for negative (falling). Dotted lines mark the evaluated deflections’ edges. The count beside the ground-truth row is its number of evaluated deflections, and the count beside each method’s row is the number of those the method resolves. At the bottom is SafeLens’s deconvolved field over frequency band and time (Section M.8), in the same colors, with a translucent veil over the cells outside the SafeLens map. The hatched band marks the first 15 ms after the stimulus, which are excluded from the evaluation in (A) and (B), so a ground-truth deflection that starts in them is not evaluated. A deflection is resolved when at least half of its total signal variation lies inside a significant region of the method with matching polarity (Section V.2). **B**. Sensitivity, as the fraction of the ground-truth deflections of the four simulated datasets that each method resolves per component SNR bin (Section V.2) and over all deflections (“all” adds 20 outside the bins). Over the 352 recorded channels (both datasets), error bars are 95% bootstrap intervals and brackets on “all” are sign-flip permutation tests of each channel’s recall difference (Section V.4), Holm-corrected over the three pairs (*** p < 0.001, n.s. not significant; gray with ZST). **C**. The percentage of the components reported by SafeLens (2848), CRP (2840) and ZST (2138) that are false, by wrong polarity or as a false detection overlapping no ground-truth deflection. **D**. The percentage of channels with false positives (FPs, Section V.2). On the left: post-CCEP FPs on the channels with CCEPs injected, and on the right: FPs from running each method on resting iEEG with no injected CCEP. Intervals (95%) are exact for resting channels and a recorded-channel bootstrap post-CCEP (exact at 0%). Dark lines mark the chance expectation, 5% for the resting channels. For the channels with CCEPs injected it is SafeLens’s rate on the resting iEEG (5.4%), scaled by the share of time that no ground-truth deflection covers (Section V.2). Gray bands are the central 95% ranges of the binomial percentages that a test at these rates gives by chance on 352 channels. Each method is at its own secondary-data threshold (Section R.1). Dashed: oracle thresholds, set on the resting data under the injected responses (unavailable in practice); marks test solid values.

### V.3 Benchmark Methods and Fairness Controls

SafeLens was compared with CRP [46] and with a z-scored detector (ZST) that thresholds the trial average [36, 69]. ZST represents the most common practice, since most CCEP studies that describe their detection method threshold the trial average [58]. CRP is the data-driven method closest to SafeLens, since it learns a response shape and its duration from each pair’s trials [46]. Like SafeLens, ZST and CRP use the trials of only one stimulating and recording pair to decide which time points of that pair’s response are significant (its significant region, Section V.2).

For each candidate duration measured from the pulse, CRP computes the scalar projection of each trial of a pair onto each of the pair’s other trials over that duration (a semi-normalized inner product) [46]. The response duration *τ* R is the candidate duration that maximizes the mean projection, the canonical response shape is the first principal component of the trials truncated to *τ* R, and the extraction significance p is the p-value of a one-sided one-sample t-test of whether the projections at *τ* R exceed zero [46]. For responses that begin some time after the pulse, CRP’s authors propose a search for the onset but do not apply it to CCEPs [46]. Our implementation adds that search, taking the onset *τ* B as the start time that maximizes the mean projection recomputed from that start time to *τ* R. CRP’s response window runs from *τ* B to *τ* R, and CRP’s score is 1 − p at every time point inside that window and zero elsewhere.

ZST thresholds the trial average after z-scoring it [36, 69]. Each trial is baseline-corrected by subtracting its mean over the interval from 500 to 5 ms before the pulse, the corrected trials are averaged, and the average is z-scored against the mean and standard deviation of its values over that interval. The absolute z-score at each time point after the post-stimulus exclusion band (the first 15 ms, Section V.2) is compared with a threshold, and ZST outputs the time points above it, which for the evaluation were divided at the peaks of the z-score into components, each with the polarity given by the order of its peaks (Section V.2). As is discussed in the next paragraph, the threshold was calibrated for each dataset on resting data in place of the original studies’ fixed threshold of 6 standard deviations [36, 69].

For a fair comparison, the three methods analyzed the same event-locked trials, verified to match trial by trial in the simulated datasets and in the patients’ recordings. Each benchmark’s threshold was calibrated, as SafeLens’s is, on the secondary recordings (Section M.1 and M.9). For each channel, each benchmark was run once on resting epochs drawn from those data without replacement under a fixed per-channel seed, as many epochs as the first cross-validation fold of SafeLens’s null holds out as rest trials (Section M.9). For each simulated dataset or patient, a benchmark’s threshold was the 95th percentile over the channels of each channel’s maximum score over time on those epochs. The benchmarks’ thresholds thus held the family-wise error at *α* = 0.05 on average over a dataset’s or patient’s channels, a weaker guarantee than SafeLens’s *α* = 0.05 on each channel (Section M.9). As a reference unavailable in practice, Figs 3B and 3D also show, as dashed outlines, each benchmark at the threshold set by the same 95th percentile on each of the two resting recordings that underlie the simulated datasets, with no CCEP injected (Section R.1). Every response on those recordings is false, so these thresholds give each benchmark a response on 17 of their 352 channels by construction, against the 19 that SafeLens reported at its own threshold. The methods also differed in filtering. In the simulated datasets and the patients’ recordings alike, CRP received the recordings unfiltered, as in its published form [46], ZST received them high-pass filtered at 1 Hz, and SafeLens received them unfiltered, since its features are blind to a constant offset and robust to slow trends (Section M.1 and M.3).

CRP returns one canonical shape and one response window for each pair and divides neither into deflections with a polarity, so it defines no component-level output [46]. Scoring it on deflections therefore exceeds its intended design scope. CRP’s authors nonetheless expect the shape to represent the response’s structure, and they show that it captures the N1/N2 shape where one is present [46]. The sign of the canonical shape was first set to the one whose centered correlation with the pair’s trial average over CRP’s response window was higher, since the sign of a principal component is arbitrary. CRP was credited with a deflection under the lenient polarity requirement of Section V.2, which asks only that a straight line fitted to this signed shape across the deflection slope in the deflection’s direction. For the measures of Section V.2 that need a component, such as the boundary error and the count of false components, components were cut from the signed shape. The signed shape was smoothed by a 25 ms moving average and split into monotonic excursions at the sign changes of its slope, and the part of each excursion inside CRP’s response window was a CRP component. Each component’s polarity was positive for a rise and negative for a fall, as for the ground-truth deflections. The 25 ms smoothing is wider than the 5 ms used for the ground-truth deflections (Section V.2) and was chosen to favor CRP. CRP’s shape is estimated from noisy trials, and a narrower smoothing was overly aggressive in fragmenting the components, while a wider smoothing reduced the accuracy of higher frequency components.

### V.4 Statistical Inference Across Methods

The three methods were scored on the same simulated channels and deflections (Section V.2), so each comparison of two methods was paired. These channels are not independent, since each of the 352 channels recorded from the two subjects appears twice among the 704 simulated channels, once in each jitter condition (Section V.1). A recorded channel’s two versions and all the deflections on them share its resting iEEG, so the tests grouped them by recorded channel. Channel recall was compared by an exact sign-flip test of each recorded channel’s difference in recall between two methods, and its 95% intervals were computed by a bootstrap over recorded channels with 20,000 draws. Deflection recall was compared by an exact sign-flip permutation test over recorded channels of each channel’s recall difference, using the clustered McNemar statistic [101]. The difference in the area under the ROC curve was tested within each SNR bin and over all channels by a bootstrap with 20,000 draws. Each draw resampled whole recorded channels with their response and rest values for all three methods. For channel recall and for the area under the curve separately, the p-values of the ten comparisons of SafeLens with CRP and with ZST in the four SNR bins and over all channels were adjusted by Holm’s procedure [102]. CRP and ZST were not compared with each other. For deflection recall, the p-values of the three pairs of methods were adjusted by Holm’s procedure. To weight every recorded channel equally in deflection recall, a sign test also compared the fraction of each channel’s deflections that SafeLens resolved with the fraction CRP resolved (Section R.3). Boundary errors (Section V.2) were compared between SafeLens and CRP, and between SafeLens and ZST, each pair over the deflections both resolved (Fig 4 and S1 Fig, Section R.4). On the deflections both resolved, a sign-flip permutation test within each SNR bin and each duration bin compared each recorded channel’s mean paired difference in boundary error (the mean of the onset and offset errors) with zero (Fig 4C, D). The spread of each method’s signed onset and offset errors, measured as the range between their 10th and 90th percentiles, was compared by permuting the two methods’ errors within each recorded channel (Fig 4E). Mean boundary errors were compared three times (Fig 4B). On the deflections both resolved, a sign-flip permutation test over recorded channels compared SafeLens with CRP. On the deflections only SafeLens resolved, a permutation of the two sets’ labels within each recorded channel compared SafeLens’s error with its error on the deflections both resolved, and a bootstrap over recorded channels compared it with CRP’s error on the deflections both resolved weighted to the SNR of those only SafeLens resolved (Section R.4). The same sign-flip test compared SafeLens with ZST on the deflections both resolved. Within each panel of Fig 4, the p-values were adjusted by Bonferroni’s correction, except for the comparison with ZST. Fig 3 makes no test within an SNR bin, and its results per bin are shown only to describe them. The lowest or highest SNR bin was merged into its neighbor when it contained fewer than 5% of a figure panel’s channels or deflections. The one exception is Fig 2’s lowest bin, below 5 dB, which contains 32 of the 704 channels (4.5%) and was kept to show ZST close to chance (AUC 0.62): there, SafeLens detects 16 of the 32 channels, CRP 10 and ZST 2. Had it been merged into the 5 to 15 dB bin, as the rule would require, SafeLens’s recall and AUC in the merged bin would still be significantly above CRP’s (71.0% against 54.6%, Holm p = 0.0006; 0.919 against 0.887, Holm p = 0.03) and ZST’s (24.0% and 0.748, both Holm p ≤ 0.0004), each metric Holm-corrected over the eight comparisons of that layout.

**Fig 4.**
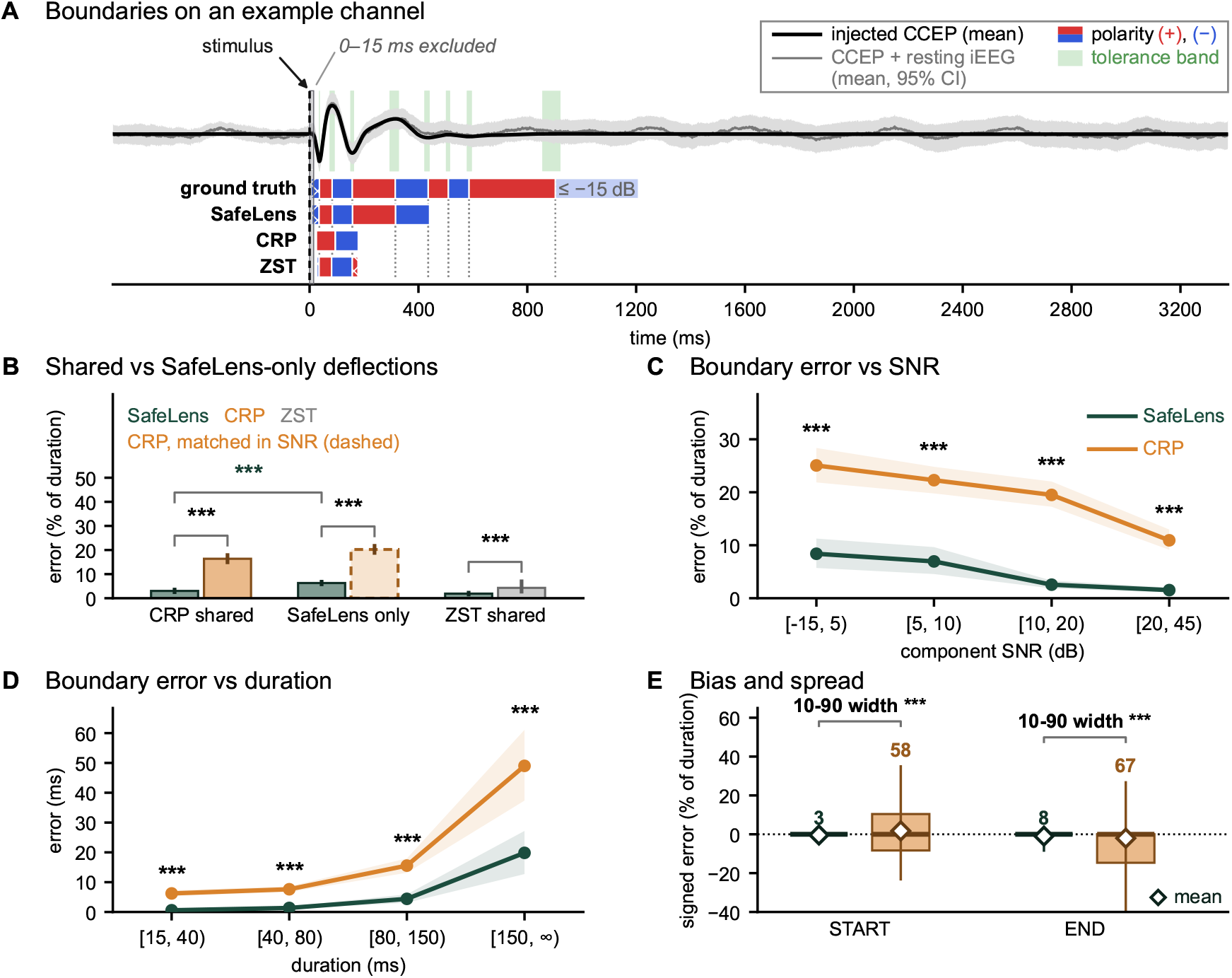
Boundary error of SafeLens and CRP on the simulated deflections. **A**. An example channel, with the tolerance band of each boundary of an evaluated deflection in the range shown (Section V.2). Its traces are the injected CCEP’s mean and the mean over all trials of the resting iEEG with the CCEP injected, baselined as in Fig 3, with a simultaneous 95% confidence interval (the mean ± 3.56 SEM, set as in Fig 3). Below the traces are the ground-truth deflections (faded and marked at ≤−15 dB) and the components of SafeLens, canonical response parameterization (CRP) and ZST, red for positive (rising) and blue for negative (falling), with dotted lines at the boundaries of the evaluated deflections. Crosshatching marks a ground-truth deflection that starts inside the excluded 0 to 15 ms and is not evaluated, and any component that is excluded for not having qualified as a valid detection (Section V.2). **B**. Mean boundary error with 95% bootstrap intervals over recorded channels, for the 642 deflections SafeLens and CRP both resolve, the 1244 of the 1250 only SafeLens resolved that have a boundary error, and the 745 SafeLens and ZST both resolve. A deflection’s boundary error is the mean of its absolute onset and offset errors, each as a percentage of its duration (Section V.2). The dashed bar is CRP’s mean error on the deflections both resolve, reweighted so that each component-SNR bin (−15 to 5, 5 to 10, 10 to 20 and 20 to 45 dB) counts as often as in the SafeLens-only deflections. Brackets test over recorded channels (Bonferroni over the three brackets without ZST; *** p < 0.001): SafeLens against CRP or ZST by sign-flip (shared) and against CRP by bootstrap (dashed bar), and SafeLens’s two bars (green) by permuting labels within channel; the SafeLens-only deflections are weaker (median component SNR 11.1 vs 18.4 dB). ZST appears in (B), and S1 Fig repeats (C to E) for SafeLens and ZST. **C**. Boundary error against component SNR for the 642 deflections with a boundary error from both methods (n = 43, 79, 232 and 288 from the lowest bin to the highest). Lines are means with 95% bootstrap bands over recorded channels, and each bin has a sign-flip test of each channel’s mean error difference between SafeLens and CRP, Bonferroni-corrected over bins. **D**. The same error in milliseconds against ground-truth duration (n = 174, 295, 142 and 31), with bands and tests as in (C). **E**. Signed error at each onset (START) and offset (END) of the deflections in (C), negative when the edge falls before its tolerance band and zero inside it. Boxes in the colors of (C) span the 25th to 75th percentiles with a line at the median, and whiskers span the 10th to 90th percentiles. The number above each whisker is that span in percentage points, and brackets compare the spans by swapping the methods’ errors within channels, Bonferroni-corrected over the two.

### V.5 Human iEEG Application

After the same Laplacian re-referencing as the simulated data (Section M.1), SafeLens was applied to single-pulse CCEP recordings of eight patients with drug-resistant epilepsy, implanted with SEEG to localize the seizure onset zone. In each subject’s session, single pulses (7 mA, 0.2 ms pulse width) were delivered through one pair of contacts in blocks of 27 pulses at an inter-pulse interval of 3.4 s [103], eleven to fourteen blocks per subject (276 to 378 pulses, two blocks of one subject with fewer than 27). The resting recordings between the blocks were the primary asynchronous data, and those before the first block or after the last were the secondary recordings (Section M.1). Some gaps between the CCEP and resting blocks included TBS, whose effects in this cohort are reported in [103] and are not analyzed here. The pulses of every block, before and after TBS, were pooled into one SafeLens significance map per channel (Section M.0).

Of the 1352 recorded channels, the analysis kept those whose contact lay in gray matter in the anatomical parcellation. It excluded the stimulating contacts, the contacts that the clinical localization marked for exclusion, and the channels whose post-stimulus trace was an amplifier recovery or a drift that did not return to baseline between pulses. A channel was excluded as primarily an amplifier recovery artifact when its trial-averaged trace, smoothed by a 25 ms moving mean, never reversed direction between 15 and 500 ms after the pulse. A channel was excluded as primarily drift artifact when the mean of its trial-averaged trace from 2500 to 2900 ms after the pulse lay more than 50 µV from the trace’s mean from 500 to 5 ms before the pulse. Eleven further channels with drift that neither criterion detected were excluded on inspection. The selection left 616 channels, 54 to 96 per subject.

On these channels, SafeLens ran with seven folds of whole blocks (Section M.2). CRP and the z-scored detector received resting epochs drawn from the secondary recordings (Section V.3) and every trial that SafeLens’s seven folds held out, except one subject’s last trial, whose epoch ran past the end of that subject’s recording. CRP’s input was unfiltered, as in the simulated validation. The z-scored detector’s input was high-pass filtered at 1 Hz, as in the simulated validation, since at lower corners it reported large monotonic late excursions that neither SafeLens nor CRP, both more robust to slow drift, reported.

Since real recordings have no ground truth, the three methods were compared on three things each reports across the 616 channels, whether a channel responds, how long its response lasts and how late its last significant time point lies. A channel was responsive for SafeLens when its significance map contained at least one significant component, and for a benchmark when at least one post-stimulus time point was significant. SafeLens’s yield of responsive channels was tested against that of CRP, the primary benchmark (Section V.3), by an exact sign-flip test over patients of each patient’s difference in the percentage of its channels that responded. A response’s duration was the longest run of significant time points on a responsive channel, with runs separated by gaps of 100 ms or less joined, from its first to its last significant time point, within its significant components for SafeLens. The durations of SafeLens and CRP were compared by the Mann–Whitney U test, since they came from different sets of responsive channels.

For each method, how late the cohort’s responses extended was tested against chance. A channel’s last significant time point was the latest post-stimulus time point that the method declared significant on it, whether or not it was contiguous with the earlier ones. At each post-stimulus time, the fraction of a subject’s analyzed channels whose last significant time point lay at or beyond that time was averaged over the eight subjects with equal weight, giving the method’s cohort curve. Its chance threshold came from 2000 null replicates of the cohort in which each channel was made falsely responsive with probability 0.05, the nominal channel-level false-positive rate of all three methods (Sections M.9 and V.3). Each falsely responsive channel was given one spurious run of significant time points, with an onset uniform over the post-stimulus epoch and a length drawn from the method’s own observed lengths. The threshold was the pointwise quantile of the null curves, raised until at most 5% of them crossed it anywhere along the time axis, so that a crossing was corrected family-wise over time. A method’s extent beyond chance was the end of the first run of times at which its cohort curve lay above that threshold.

### V.6 Single Trial Analysis

Per-trial timing models need a prior giving the number of components in a response and their approximate spans [11, 66, 67], which the SafeLens map supplies (Section M.11). We tested the SafeLens map as that prior on the two high-jitter simulated datasets (Section V.1), the only data in which each deflection’s true time on every trial is known and varies. A deflection’s latency was the time of its maximum for a rise or its minimum for a fall in the average of the injected noise-free templates. A deflection’s duration was the time from its onset to its offset in that average. A deflection’s true time on a trial was the time of its maximum for a rise or its minimum for a fall in the trial’s template, searched within half the deflection’s duration on either side of its latency. A deflection’s latency jitter on the injected trials was the standard deviation of these true times as a percentage of its duration. Of the 1,738 deflections scored in these datasets (Section V.2), 1,539 had a latency jitter of 10% or more. The other 199 were left out, since a deflection whose timing barely varies leaves little per-trial timing to recover. A deflection’s single-trial SNR was the ratio in decibels of the mean square of its noise-free templates to that of the resting traces they were added to, over its time points on all trials. The analyzed deflections were the 926 of the 1,539 (on 322 channels) whose single-trial SNR was at least −25 dB, a floor set from the ideal reference (below).

The SafeLens map was compared with the standard analysis windows, from 10 to 50 ms after the pulse for the N1 [34–36, 62, 69] and from 50 to 300 ms for the N2 [34, 35, 62], and with CRP’s components. The SafeLens and CRP priors gave each deflection the span of the component it was paired with in scoring (Section V.2), or no span when the deflection was not resolved. The standard windows gave a deflection the window that contained its latency, or no span when its latency lay outside both windows. Each span was read by differentially variable component analysis (dVCA) [14], which estimates the span’s waveform from the trials and shifts each trial to match the waveform within a limit of half the span’s duration either way. For a window this limit is the half-window limit, half the window’s width. The windows were also read at a best case, with the limit narrowed to half the true duration of the longest of the window’s deflections with a latency jitter of 10% or more (a median limit of 50 ms). No analysis of real data could know this duration. Each window was also read by the conventional measure of latency, the time of each trial’s largest peak inside the window. ZST, the common practice in detection (Section V.3), was not used as a prior, since the common practice in timing is the standard windows and their largest peaks. As an ideal reference, dVCA also read each deflection’s true span, from its onset to its offset.

Since 897 of the 926 analyzed deflections had a single-trial SNR below 0 dB, the deflections were scored on super-trials, averages of trials of similar Woody lag [11]. A trial’s Woody lag for a span was the shift at which it correlated best with the average of all the channel’s trials over that span, computed from the recorded trials alone. Super-trials of 1, 2, 3, 4, 5, 6, 8 or 13 trials were formed within each event block (Section V.1), separately for each prior’s spans. Each deflection was scored at the smallest of these sizes at which averaging would bring its single-trial SNR to 0 dB, or at 13 trials when none could, and at that size under every prior. Each span was read by dVCA on its super-trials, and each window also by the largest peak on the same super-trials, each reading giving one series of times per span at every size. A deflection’s score was the Pearson correlation across the super-trials of its own size between its span’s series and the true times of the super-trials’ averaged templates. One series could serve all the deflections in one window or in one component, whereas a per-trial timing model is meant to time each deflection separately. Each span was therefore credited to the deflection its series followed best, and the other deflections the span served had no time of their own and scored 0, as did each deflection to which its prior gave no span. A correlation was large at 0.5 or more, medium from 0.3 and small from 0.1, and a deflection with a correlation below 0.1 was not tracked [104]. The priors were also compared on the 202 matched deflections, those that both SafeLens and CRP resolved and whose latency lay inside a standard window. These were scored without crediting, so that a window’s series counted for every matched deflection within the window. The deflections that shared a window had correlated true times in the simulation (a median of 0.73 between pairs), which favors the windows, since one shift per super-trial can then fit them all.

The approaches (each a prior with the way its spans were read) were compared by a two-sided sign-flip permutation test over recorded channels of each channel’s mean difference in correlation, since the deflections of one channel are not independent. The SafeLens map read by dVCA was compared with every other approach on the analyzed deflections and with every other approach and the ideal reference on the matched deflections. The SafeLens map was also compared with whichever of CRP’s components and the windows’ three approaches had the highest mean correlation in each of nine single-trial SNR bins and again in each of four latency bins. The same comparison was made at each fixed super-trial size, with every analyzed deflection scored at that size. The comparisons by SNR and by latency used all 1,539 deflections, with the spans credited separately within the analyzed deflections and within the 613 below −25 dB. The −25 dB floor was set from the ideal reference and so from none of the compared priors. The ideal reference’s mean correlation was 0.02 for the deflections below the floor and at least 0.1 (a small correlation [104]) in every SNR bin from −25 dB up. The tests were Bonferroni-corrected within each panel of Fig 5, and confidence intervals on the mean correlations of the matched deflections came from 2,000 resamplings of whole channels.

**Fig 5.**
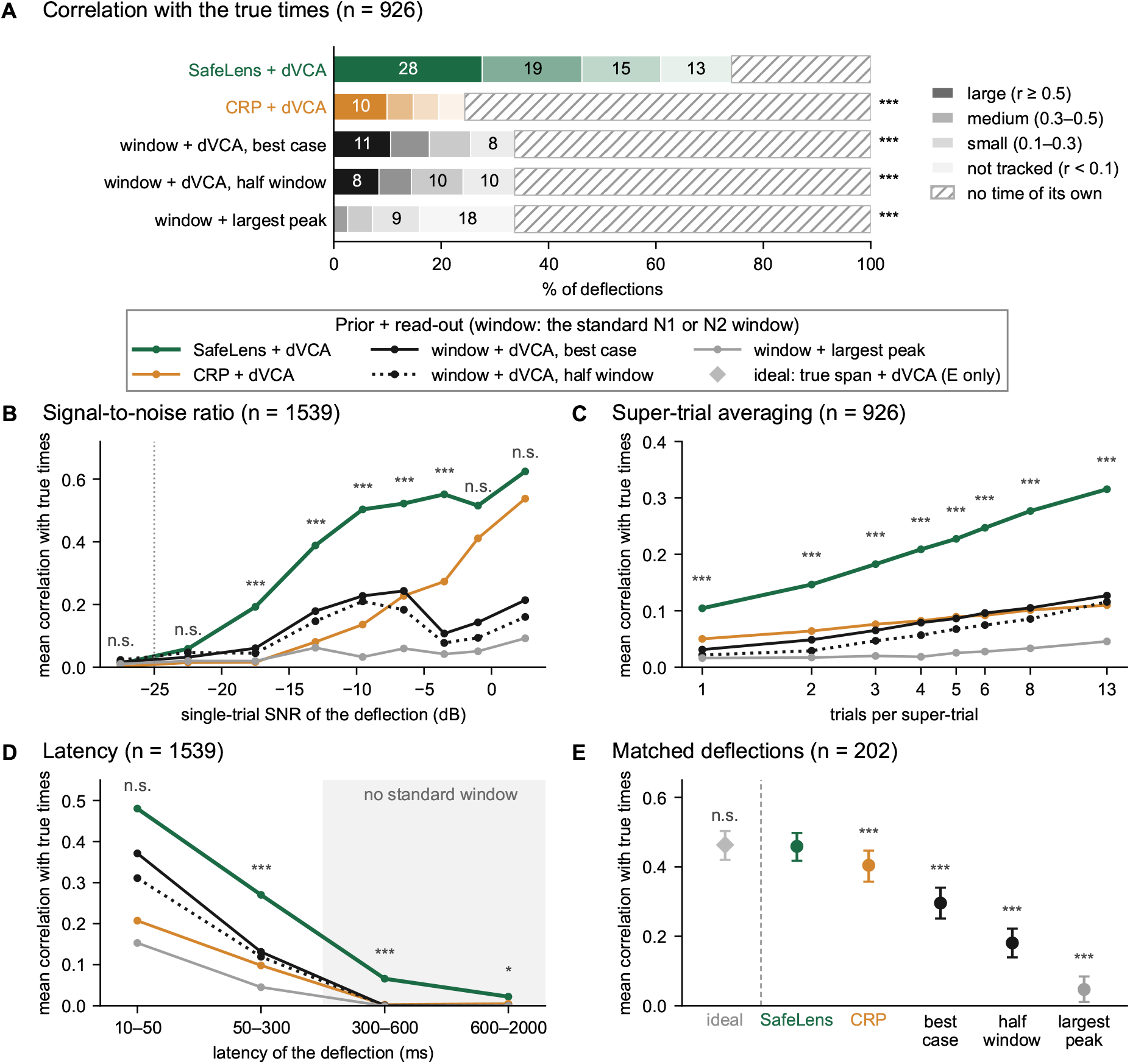
Per-trial timing of simulated deflections with SafeLens, CRP and standard windows as priors. The deflections, each with a latency jitter of 10% or more, are from the two high-jitter simulated datasets. Each approach reads the spans of a prior by differentially variable component analysis (dVCA) or by the largest peak (Section V.6). The two window variants limit each trial’s dVCA shift to half the window’s width (half window) or to half the true duration of its longest deflection (best case). Each deflection is assessed by the correlation across super-trials (averages of trials) between its estimated and true times. A response ends at its first pair of deflections at or below −15 dB, and the deflections after it are not evaluated. **A**. Percentage of the analyzed deflections, those with a single-trial SNR of −25 dB or more, by the size of their correlation [104]. Darker tints of each approach’s color mark larger correlations. Hatching marks deflections with no time of their own, given no span by their prior or sharing a span whose times were credited to another deflection. **B**. Mean correlation over all deflections by single-trial SNR, with the −25 dB floor dotted. **C**. Mean correlation over the analyzed deflections by the number of trials per super-trial. **D**. As (B), by latency. **E**. Mean correlation over the matched deflections, those that SafeLens and canonical response parameterization (CRP) both resolved with a latency inside a standard window, with 95% confidence intervals from resampling channels. Marks give sign-flip permutation tests over recorded channels of each channel’s mean difference in correlation, Bonferroni-corrected within each panel (* p < 0.05, *** p < 0.001, n.s. not significant). Each mark compares SafeLens + dVCA with the approach of its row in (A) or its column in (E), and in (B to D) with the best other approach at that point.

## R. Results

### R.1 False-positive rate on resting iEEG and post-CCEP

We first asked how often each of the three methods (SafeLens, CRP and ZST, Section V.3) reports a response in resting iEEG with no injected CCEP. The pool of resting data set aside for the null (the secondary recordings, Section M.1 and M.9) could not serve as this test, since every method’s threshold is calibrated on that pool and so flags about 5% of its channels by construction (Section V.3). We therefore ran each method, with no CCEP injected, on the resting iEEG that underlies the simulated datasets and shares no data with that pool (Section V.1). On these 352 channels SafeLens reported a response on 19 (5.4%), CRP on 10 (2.8%) and ZST on 7 (2.0%), each within or below the central 95% range of counts that a test calibrated at 5% gives by chance (10 to 26 of 352, Fig 3D). Since each of SafeLens’s forests is trained on a 500 ms searchlight window (Section M.3), an injected CCEP could raise SafeLens’s map up to 500 ms from the CCEP, at times the CCEP’s deflections do not cover. We therefore also tested whether an injected CCEP raises SafeLens’s false positives, counting a reported response as false when it overlaps none of the CCEP’s deflections. On the 704 channels of the four simulated datasets (the two subjects in two jitter conditions each), SafeLens reported such a false positive on 20 channels (2.8%, Fig 3D). Since 66.5% of the time from 15 ms after the pulse lies outside every deflection, about 3.6% of channels were expected to have one by chance (66.5% of SafeLens’s resting rate of 5.4%, Section V.2). SafeLens’s 20 of 704 channels (2.8%) lie within the central 95% range around that rate (1.7% to 5.7%, 6 to 20 of the 352 recorded channels). In summary, every method’s false-positive rate on resting iEEG lay within or below the range expected of a test at the nominal 5%, and an injected CCEP did not raise SafeLens’s false positives above chance.

### R.2 SafeLens accurately detects responsive channels

We next focused on the coarsest level of detection: whether each of the three methods detects that a channel has any event-related response at all. Each channel’s statistic was the maximum of SafeLens’s or ZST’s detection statistic after the 15 ms exclusion band, and for CRP its extraction significance p (Sections V.2 and V.3, Fig 2A). For the ROC, the positives are the channels with an injected CCEP and the negatives are the same channels’ resting iEEG with no CCEP injected, on which false positives were counted in Section R.1. Recall was measured at the threshold each method uses throughout, which SafeLens sets for each channel and each benchmark for each subject (Sections M.9 and V.3). Pooled over the 704 channels of the four simulated datasets, the area under the ROC curve was 0.978 for SafeLens, 0.965 for CRP and 0.891 for ZST (Fig 2D). SafeLens’s AUC exceeded CRP’s by 0.013 (95% interval 0.005 to 0.021, Holm p = 0.009, Fig 2E) and exceeded ZST’s with Holm p ≤ 0.0005. Recall at 5% FWER was 91.8% for SafeLens, 84.5% for CRP and 67.8% for ZST (Fig 2F), and SafeLens’s recall exceeded CRP’s (Holm p = 1.5 × 10^−6^) and ZST’s (Holm p < 10^−10^). Of the 79 channels that only one of SafeLens and CRP detected, SafeLens detected 65 (82%) and CRP 14. Stratified by SNR bin (Fig 2F), SafeLens’s recall exceeded CRP’s in every bin and significantly at 5 to 15 dB, 75.5% against 59.6% (Holm p = 0.0049), and at 15 to 30 dB, 98.5% against 93.0% (Holm p = 4.1 × 10^−4^). The margins of 50.0% against 31.3% below 5 dB (Holm p = 0.078) and 100% against 98.5% at 30 dB or more (Holm p = 0.25) were not significant. SafeLens’s AUC exceeded CRP’s significantly at 5 to 15 dB (0.939 against 0.908, Holm p = 0.049), at 15 to 30 dB (0.997 against 0.990, Holm p = 0.013) and at 30 dB or more (1.000 against 0.996, Holm p = 0.049, Fig 2D). The difference at 30 dB or more was 0.0035 (95% interval 0.0001 to 0.0101, Fig 2E). Below 5 dB the two AUCs did not differ significantly (0.822 against 0.789, Holm p = 0.37). ZST’s recall was 6.3%, 27.8%, 76.5% and 94.3% across the same four bins, and SafeLens exceeded ZST in every bin in recall (Holm p ≤ 0.0049) and in AUC (Holm p ≤ 0.0066). In summary, SafeLens’s recall was higher than CRP’s in every bin and significantly higher at 5 to 15 dB, at 15 to 30 dB and over all channels, and significantly higher than ZST’s in every bin.

### R.3 SafeLens recovers more than twice the deflections with the correct polarity

Once we had established how the three methods differ in channel-level CCEP detection, we asked how well each of them recovers each of the individual deflections that make up a simulated CCEP. We consider a ground-truth deflection resolved when a method marks as significant, with the correct polarity, the time points over which at least half of the deflection’s amplitude change occurs (Section V.2, Fig 3A). We included a ground-truth deflection in this analysis if it either reached the −15 dB inclusion floor as a whole against the noise remaining in the averaged data before its channel’s response ended at its first pair of deflections at or below that floor (3945 deflections), or if one of the three methods resolved it precisely at both edges (Section V.2). The resulting 3965 deflections are scored identically for all three methods, with ZST’s polarity read from the order of its z-score peaks (Section V.3). SafeLens resolved 47.9% of CCEP deflections with the correct polarity, against 18.4% for CRP and 19.4% for ZST. SafeLens’s recall exceeded CRP’s in every SNR bin (Fig 3B). Recall was 37% against 7% at 0 to 5 dB, 62% against 16% at 5 to 10 dB and 86% against 30% at 10 to 20 dB, and in the highest bin SafeLens resolved 97% against CRP’s 61%. Over all 3965 deflections, with the deflections clustered by recorded channel, SafeLens resolved significantly more than CRP and than ZST (sign-flip permutation tests over the 352 recorded channels, Holm-corrected p = 3 × 10^−6^ for each, the smallest attainable with 10^6^ permutations, Fig 3B).

Conversely, specificity is the fraction of the components reported by each of the three methods that is false, either by a polarity that disagrees with its overlapping ground truth’s net direction across the component’s own span or by overlapping with no ground-truth deflection at all (Fig 3C). Of SafeLens’s 2848 reported CCEP components, 2.8% were false, 2.0% by polarity and 0.8% by overlapping with no deflection. Of CRP’s 2840 components 30.0% were false (30.0% and 0.0%), and of ZST’s 2138 components 14.6% were false (13.3% and 1.3%).

SafeLens’s advantage in recall does not depend on pooling multiple deflections per channel. With each channel scored by the fraction of its deflections a method resolved, SafeLens exceeded CRP in 315 of the 352 recorded channels and CRP exceeded SafeLens in 13 (24 tied). SafeLens also resolved more deflections than CRP within each subject’s two simulated datasets (46.8% against 17.6% and 48.8% against 19.1%). The median per-channel recall was 50% for SafeLens against 17% for CRP (sign-flip permutation test over the 352 recorded channels of each channel’s recall difference, Holm-corrected over the three pairs of methods, p = 3 × 10^−6^, the smallest attainable with 10^6^ permutations), and a channel-resampling bootstrap estimated the pooled recall gap at 29 percentage points (95% CI 28 to 31). The deflections that SafeLens exclusively resolved were predominantly low SNR deflections (Fig 3B). In the simulated data, the deflections only SafeLens resolved are also the late ones, since component SNR decreased with latency (Spearman *ρ* = −0.60, median 14 dB before 50 ms against −5 dB beyond 200 ms). In summary, SafeLens resolved 48% of the ground-truth deflections against 18% for CRP and 19% for ZST, and the deflections it added were predominantly at low SNR and late in the response.

### R.4 SafeLens recovers deflection boundaries more accurately

We next asked whether SafeLens recovers more deflections by placing its boundaries less carefully. To answer this question, we measured boundary error against a symmetric tolerance band as a percentage of each deflection’s own duration. The tolerance band spans the flat, low-information interval on either side of a true boundary, so an edge within it scores zero error and error grows with distance beyond it (Fig 4A). A deflection’s error was the mean of its onset and offset errors, and of the 648 CCEP deflections that both SafeLens and CRP resolved, 642 had this error from both methods. For these 642, SafeLens’s median error was 0.0% of duration against 15.3% for CRP, and its mean 3.0% against 16.4% (sign-flip test over recorded channels, Bonferroni-corrected p < 0.001, Fig 4B). Across the eight per-bin tests of Fig 4C and D (four SNR bins and four duration bins), SafeLens’s mean error was the lower in all eight and significantly lower in all eight (sign-flip tests over recorded channels, Bonferroni-corrected within each panel). SafeLens’s onsets fell early more often than late (14.0% against 7.0%), with a mean signed error of −0.4% of duration. Its offsets fell early more often than late (15.6% against 9.5%) and by about as much (a mean of 16.1% of duration against 16.5%), so its mean signed offset error was −0.9%. CRP’s mean signed errors were near zero, +1.7% at onset and −2.0% at offset, but CRP’s signed errors spread far more widely than SafeLens’s. The 10th-to-90th percentile range of the signed error was 2.6 percentage points for SafeLens against 58.1 for CRP at onset and 8.4 against 67.1 at offset (permutation swapping the methods’ errors within recorded channels, Bonferroni-corrected p < 0.001, Fig 4E).

Over all the deflections each method resolved, mean boundary error was 5.2% of duration for SafeLens and 17.7% for CRP (1891 and 723 deflections with a boundary error, Section V.2). Of the 1250 deflections that only SafeLens resolved, 1244 had a boundary error. These 1244 deflections had lower SNR than the 642 deflections both methods resolved, a median of 11.1 against 18.4 dB (interquartile ranges 5.4 to 16.7 and 11.9 to 24.7 dB). SafeLens’s mean boundary error on them was 6.3% of duration against 3.0% on the 642 (permutation of the set label within recorded channels, Bonferroni-corrected p = 1.5 × 10^−4^, Fig 4B). CRP resolved none of the 1244, so its error on them was estimated from its mean error on the 642 in each SNR bin, weighted by that bin’s share of the 1244. The estimate was 20.2% against SafeLens’s 6.3% (Bonferroni-corrected p < 0.001, Fig 4B). In summary, SafeLens identifies more deflections while also recovering their boundaries more accurately. The deflections only it resolves have a mean boundary error of 6.3% of duration against 3.0% for the deflections both methods resolve, and CRP’s estimated error at their SNR is 20.2%.

Fig 4B also compares SafeLens with ZST, on the 745 deflections those two both resolved, where SafeLens’s mean boundary error was 1.9% of duration against 4.3% for ZST (sign-flip test over 227 recorded channels, p = 1.2 × 10^−4^). For visual simplicity, Fig 4C to E show SafeLens and CRP only, and S1 Fig shows the same panels for SafeLens and ZST on these 745 deflections. There, SafeLens’s error was significantly lower in one SNR bin and one duration bin, and the 10th-to-90th percentile range of its signed offset error was narrower, 1.2 percentage points against 13.5 (Bonferroni-corrected p = 0.013, 0.025 and p < 0.001). The range of the signed onset error did not differ between the two methods, and ZST’s error was significantly lower than SafeLens’s in no bin. SafeLens’s margins over ZST were smaller than over CRP because ZST resolved almost no deflections below 5 dB (Fig 3B), so only 6 of these 745 deflections had an SNR below 5 dB, against 43 of the 642 shared with CRP, while on the deflections of 20 dB or more, over half of the 745, both methods’ errors were small (1.2% and 2.1% of duration, S1 Fig A).

### R.5 Robustness to trial-to-trial variability

CRP allows only the CCEP’s amplitude to vary across trials [46], so its recall is expected to drop when the CCEP also varies in latency and morphology. We therefore tested whether each of the three methods’ recall dropped under the high-jitter condition, whose CCEP varied strongly across trials in latency, amplitude and morphology (Section V.1, panels A and B in S3 Fig). Each method’s recall on the high-jitter deflections was compared with its recall on zero-jitter deflections matched to them in onset, duration and SNR. CRP’s recall dropped from 23.1% on the matched zero-jitter deflections to 18.9% on the 1,736 high-jitter deflections (−4.2 percentage points, Bonferroni-corrected p = 0.0015, panel C in S3 Fig). SafeLens’s recall dropped from 53.1% to 52.2% and ZST’s from 22.8% to 21.8%, and neither drop was significant (Bonferroni-corrected p = 1 for each). The difference between CRP’s drop and SafeLens’s drop was not significant (Bonferroni-corrected p = 0.23).

Jitter could also make SafeLens’s and CRP’s boundaries less accurate on the deflections that both still resolve. Boundary error (Section R.4) was compared on the 642 deflections (344 with zero jitter and 298 with high jitter) for which both SafeLens and CRP had a boundary error (panel D in S3 Fig). SafeLens’s mean boundary error went from 3.4% of the deflection’s duration with zero jitter to 2.6% with high jitter, and CRP’s from 16.0% to 16.8%. Neither change was significant (Bonferroni-corrected over the two methods, p = 0.22 and 1.0). In both conditions SafeLens’s mean boundary error was lower than CRP’s (corrected p < 0.001 for each). In summary, of the three methods only CRP’s recall dropped significantly under high jitter, and neither SafeLens’s nor CRP’s mean boundary error changed significantly.

### R.6 SafeLens map as a prior for per-trial timing models

We next asked how closely a per-trial timing model follows the true times of the simulated deflections when the SafeLens map supplies its prior, compared with the standard analysis windows from which per-trial latencies are usually read (Section V.6). With the SafeLens map as the prior of the per-trial timing model dVCA, 27.6% of the 926 analyzed deflections had a large correlation with their true times (Fig 5A). The shares were 10.6% with the standard windows at their best case, 8.4% with the half-window limit, 2.6% when the windows were read by each trial’s largest peak and 9.9% with CRP’s components. With the SafeLens map, 60.9% of the deflections were tracked (a correlation of 0.1 or more), against 25.5% with the best-case windows and 19.6% with CRP’s components. The approaches differed most in the share of deflections left with no time of their own (hatched in Fig 5A). SafeLens had not resolved 25.3% and CRP 74.9%, and under the windows’ three approaches 39.2% shared a window with a deflection that the window’s series followed better and 27.1% lay outside both windows. Over all 926 deflections the mean correlation with the true times was 0.28 with the SafeLens map and 0.09 with CRP’s components. The windows had 0.11 at their best case, 0.10 with the half-window limit and 0.04 with the largest peak. The SafeLens map’s mean correlation per channel was higher than CRP’s on 247 of the 322 recorded channels (lower on 34) and higher than the best-case windows’ on 249 (lower on 59). The SafeLens map’s differences from all four other approaches were significant after Bonferroni correction, by sign-flip permutation tests over the recorded channels of each channel’s mean difference in correlation (Bonferroni-corrected p = 4 × 10^−6^ for each, the smallest attainable with 10^6^ random sign flips). The SafeLens map led the best-case windows in each high-jitter simulated dataset (27.1% against 10.6% and 28.1% against 10.6% with a large correlation). The SafeLens map also led the best-case windows with the deflections scored without crediting (27.6% against 17.8% with a large correlation) and with the windows scored on the SafeLens super-trials (27.6% against 8.5%).

The SafeLens map had a higher mean correlation than CRP’s components and the windows at every fixed number of trials per super-trial, rising from 0.10 on single trials to 0.32 at 13 trials against at most 0.13 (Fig 5C). At each number of trials the difference from the best of CRP’s components and the windows was significant after correction (sign-flip permutation tests over the 322 recorded channels, Bonferroni-corrected p = 8 × 10^−6^ at each number). Over all 1,539 deflections by single-trial SNR, the SafeLens map exceeded the best of CRP’s components and the windows significantly from −20 to −2 dB, with mean correlations of 0.19 to 0.55 against 0.06 to 0.27 (Fig 5B). Above −2 dB the SafeLens map had 0.52 to 0.62 and CRP’s components 0.41 to 0.54, with no difference significant after correction. Below −20 dB none of the five approaches exceeded 0.06. By latency, the SafeLens map led the best-case windows significantly from 50 to 300 ms (0.27 against 0.13) and did not differ significantly from them on the deflections from 10 to 50 ms, on 20 recorded channels (0.48 against 0.37, Fig 5D). After 300 ms, where no standard window lies, the SafeLens map exceeded CRP’s components significantly from 300 to 600 ms (0.066 against 0.002) and from 600 to 2,000 ms (0.022 against 0.005). Of all 1,539 deflections, the 716 after 300 ms included 29 with a large correlation under the SafeLens map and 1 under CRP’s components.

Because the priors gave different shares of the deflections a span, they were also compared on the 202 matched deflections, to which every prior gave one (Fig 5E). There the mean correlation was 0.46 with the ideal reference (each deflection’s true span), while the windows had 0.30 at their best case, 0.181 with the half-window limit and 0.047 with the largest peak. Each of the windows’ three approaches was significantly lower than the SafeLens map after correction (sign-flip permutation tests over the 159 recorded channels, Bonferroni-corrected p = 5 × 10^−6^ for each). CRP’s components had 0.404 and were also significantly lower than the SafeLens map after correction (Bonferroni-corrected p = 1.4 × 10^−4^), while the ideal reference did not differ significantly from it. The SafeLens map’s lead over CRP’s components in large correlations (Fig 5A) came almost wholly from the deflections that CRP did not resolve. Of the 926 analyzed deflections, 256 had a large correlation with the SafeLens map and 92 with CRP’s components. Of the 694 that CRP did not resolve, and to which its components therefore gave no span, 153 had a large correlation with the SafeLens map and none with CRP’s components. These 694 were weaker and later than the 232 that CRP resolved (median single-trial SNR −17.6 against −10.0 dB, median latency 254 against 123 ms). In summary, the timing model followed the true times more closely with the SafeLens map as its prior than with the standard windows, on all the analyzed deflections and on the matched ones. It also tracked more deflections than with CRP’s components, and followed the matched ones more closely.

### R.7 Illustrative application to human iEEG

To complement the ground-truth simulation, we applied SafeLens across eight subjects from an empirical single-pulse iEEG CCEP dataset (Fig 6). Because real recordings have no ground truth, detection accuracy cannot be scored, and the three methods are compared on what each reports across the same 616 gray-matter channels (Section V.5). With all three methods run on the same trials, SafeLens classified a mean of 68.0% of each patient’s channels as responsive (413 of 616 channels, 67.0%), against 58.1% for CRP (344, 55.8%) and 51.6% for ZST (301, 48.9%) (Section V.5, Fig 6A). SafeLens classified more channels as responsive than CRP in 6 of the 8 patients and fewer in none (exact sign-flip test over patients, p = 0.031, Fig 6A). Of the 105 channels on which SafeLens and CRP disagreed, 87 were responsive by SafeLens alone and 18 by CRP alone. The response per responsive channel, with gaps of 100 ms or less joined, had a median duration of 337 ms for SafeLens versus 175 ms for CRP (Mann–Whitney p < 10^−10^) and 162 ms for ZST (Fig 6B). The fraction of channels significant at each time point peaked within the first 200 ms for all three methods and declined most slowly for SafeLens (Fig 6C). To test how late after the stimulus significance extends beyond chance across the cohort, the eight subjects were given equal weight (Section V.5). At each post-stimulus time the fraction of a subject’s channels significant at or after that time was compared with a chance threshold corrected family-wise across the time axis (Section V.5). SafeLens’s fraction remained above the chance threshold out to 1040 ms, CRP’s to 455 ms and ZST’s to 570 ms (Fig 6D). The median offset of significance per responsive channel was 434 ms for SafeLens versus 244 ms for CRP and 264 ms for ZST. Fig 6E and F show one subject’s channels grouped by anatomical region and ordered by distance from the stimulation site, first as SafeLens’s polarity-resolved components and then colored by the method detecting at each time point. In summary, SafeLens classified more channels as responsive than either benchmark across the eight neurosurgical patients, and the fraction of its channels significant at or after a given time remained above chance to 1040 ms after stimulation, against 455 ms for CRP.

**Fig 6.**
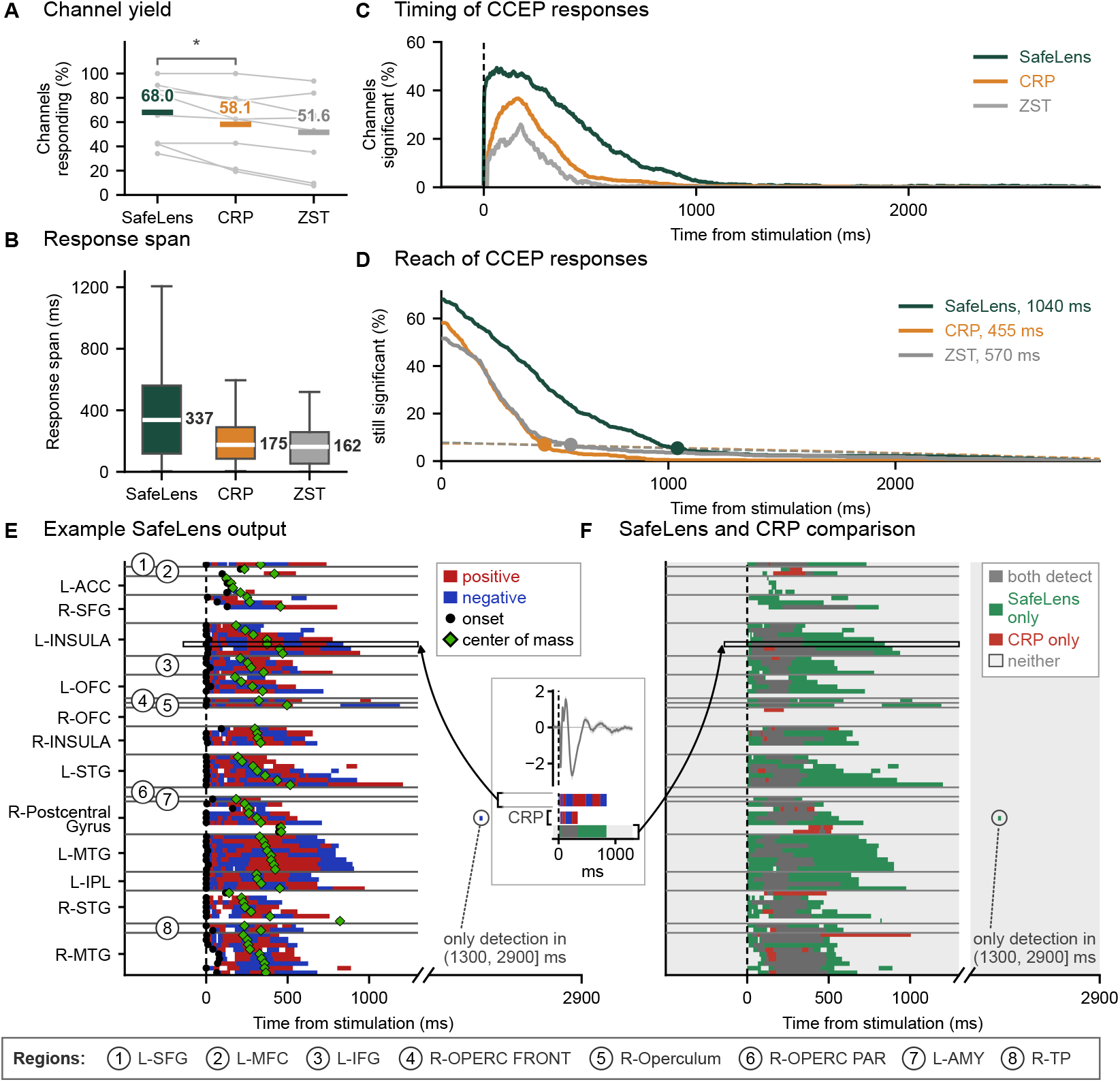
Application of SafeLens to human iEEG CCEPs (eight subjects, 616 gray-matter channels). SafeLens, canonical response parameterization (CRP) and the z-scored trial-averaging detector (ZST) were run on the same trials, with ZST high-pass filtered at 1 Hz (Section V.5). **A**. The percentage of each patient’s channels with any detected response, one gray line per patient, with each method’s mean over the eight patients marked and printed. SafeLens detects more responding channels than CRP in 6 of 8 patients and fewer in none (exact sign-flip test over patients, * p < 0.05). **B**. Response span per responsive channel, gaps ≤ 100 ms joined (n = 413, 344, 301). Boxes span the quartiles with the median (white line) printed beside them, and whiskers extend to 1.5 times the interquartile range with points beyond them not drawn. **C**. Timing of CCEP responses, as the percentage of channels significant at each time after stimulation. **D**. Reach of CCEP responses beyond chance. The curves give the percentage of each subject’s channels whose last significant time point lies at or beyond time t, averaged over the eight subjects with equal weight. Each method’s chance threshold (dashed, the three nearly coinciding) is set so that at most 5% of the null replicates exceed it at any time (Section V.5). The threshold thus corrects each crossing family-wise across the whole time axis. Dots mark the end of each method’s first contiguous run above its chance threshold, at the time given in the legend. **E**. One patient’s SafeLens components on channels grouped by anatomical region, regions ordered by distance from the stimulation site and channels within one by center of mass, nearest and soonest at the top. Each channel’s onset and center of mass are those of its whole response. Gray lines separate the regions. Eight small regions are numbered beside their rows and named in the key below (E) and (F). The time axis of (E) and (F) is compressed after 1,300 ms, where the only detection is circled. The inset shows one channel’s trial average with its SafeLens components (upper bar), CRP’s components (middle bar) and its row of (F) (lower bar), and the arrows point to that channel’s outlined rows in (E) and (F). **F**. The same channels in the same order, colored at each time point by which of SafeLens and CRP detect a response, dark gray for both, green for SafeLens only, red for CRP only and light gray for neither.

## D. Discussion

### D.1 Summary of findings

SafeLens is a statistical machine learning framework designed to recover evoked response deflections that conventionally evade detection due to their low amplitude or trial variability. It extracts the presence, boundaries, and polarity of these components under strict family-wise error control, unconstrained by fixed analysis windows, predefined signal features, or assumptions of trial invariance. Here, we use SafeLens to decompose single-channel CCEPs into separately tested deflections, each with its polarity and boundaries. Although canonical response shapes and trial averages can be partitioned post hoc into candidate deflections (Section V.3), to our knowledge, no prior CCEP method yields a statistically controlled decision on the presence, boundaries, and polarity of each deflection [58].

When evaluated against existing methods on both simulated ground truth and clinical intracranial recordings, SafeLens demonstrated significant advantages in detecting and characterizing evoked responses. We report seven main findings:

#### 1) False-positive control

SafeLens’s false positives stayed within the range expected of a test at the nominal 5% level. On the resting recordings that underlie the simulated datasets, with no CCEP injected, SafeLens reported a response on 5.4% of the 352 channels. Across the 704 simulated channels with injected CCEPs (the 352 channels in each of two jitter conditions), SafeLens reported a false positive overlapping no ground-truth deflection on 2.8% of channels, within the range expected by chance over the response-free portion of the epoch (Section R.1).

#### 2) Channel detection

With each method at its own threshold for a 5% family-wise error rate, SafeLens detected 91.8% of the simulated channels with a CCEP, against 84.5% for CRP (Holm-corrected p = 1.5 × 10^−6^) and 67.8% for the z-scored detector (Holm-corrected p < 10^−10^, Section R.2, Fig 2). Across all thresholds, the area under the ROC curve was 0.978 for SafeLens, significantly exceeding both 0.965 for CRP (Holm-corrected p = 0.009) and 0.891 for the z-scored detector (Holm-corrected p ≤ 0.0005).

#### 3) Component recall

Of the 3965 simulated deflections evaluated, SafeLens resolved 48% with the correct polarity against 18% for CRP (Holm-corrected p = 3 × 10^−6^, Section R.3, Fig 3). The z-scored detector resolved 19%. SafeLens resolved more deflections than CRP in every range of SNR, with the widest margins between 5 and 20 dB. Of the components each method reported, only 2.8% of SafeLens’s were false (having the incorrect polarity or overlapping no deflection), compared to 30.0% of CRP’s and 14.6% of the z-scored detector’s.

#### 4) Boundary accuracy

On the 642 shared deflections, for which both SafeLens and CRP had a boundary error, SafeLens’s mean boundary error was 3.0% (median 0.0%) of the deflection’s own duration, against 16.4% (median 15.3%) for CRP (Bonferroni-corrected p < 0.001, Section R.4, Fig 4). The 1250 deflections that only SafeLens resolved were weaker than the shared ones (median SNR 11.1 against 18.4 dB across the 1244 evaluated for boundary error). On those deflections, SafeLens’s mean boundary error was 6.3% (median 2.2%), well below CRP’s estimated mean of 20.2% at their SNR (Bonferroni-corrected p < 0.001).

#### 5) Robustness to trial-to-trial variability

SafeLens’s recall did not change significantly between the zero-jitter condition and the high-jitter condition, whose trials varied in latency, amplitude and morphology (53.1% to 52.2% of deflections matched across the two conditions, not significant after correction). The z-scored detector’s recall did not change significantly either (22.8% to 21.8%, not significant after correction), while CRP’s fell from 23.1% to 18.9% (corrected p = 0.0015, Section R.5, S3 Fig). On the shared deflections, SafeLens’s mean boundary error was 3.4% in the zero-jitter condition and 2.6% in the high-jitter condition, while CRP’s went from 16.0% to 16.8%. Neither SafeLens’s nor CRP’s mean boundary error changed significantly with jitter (Section R.5).

#### 6) The SafeLens map as a prior

To estimate each trial’s timing in the simulated high-jitter datasets, we supplied the spans of the SafeLens map’s components as the prior of a per-trial timing model (Section R.6, Fig 5). With the SafeLens map as its prior, the model’s estimated times had a large correlation (r ≥ 0.5) with the true times for 28% of the 926 analyzed deflections, against 11% with the standard N1 and N2 windows (even at their best case, where each trial’s shift was limited to half the true duration of the window’s longest deflection) and 10% with CRP’s components. This advantage held even on the matched deflections, to which all three priors gave a span, where the mean correlation under the SafeLens map significantly exceeded that under the best-case windows (Bonferroni-corrected p = 5 × 10^−6^) and under CRP’s components (Bonferroni-corrected p = 1.4 × 10^−4^), and did not differ significantly from that under each deflection’s true span. By contrast, the conventional measure of latency, each trial’s largest peak inside a standard window, gave a mean correlation of only 0.047 on the same matched deflections. These results support using the SafeLens map in place of the standard windows or CRP’s components as the prior of a per-trial timing model (Section D.3).

#### 7) Detection of long-duration CCEPs in eight patients

Across the 616 gray-matter channels recorded in eight patients, SafeLens classified 67.0% as responsive, compared to 55.8% for CRP and 48.9% for the z-scored detector, detecting more responsive channels than CRP in six of the eight patients and fewer in none (exact sign-flip test over patients, p = 0.031). A response’s duration on a responsive channel, the longest run of significant time points with gaps of 100 ms or less joined (Section V.5), had a median of 337 ms for SafeLens, against 175 ms for CRP (p < 10^−10^) and 162 ms for the z-scored detector. At the cohort level, the share of channels with a significant time point at or after a given time stayed above chance until 1040 ms for SafeLens, against 455 ms for CRP and 570 ms for the z-scored detector (Section R.7, Fig 6).

Taken together, these findings demonstrate that SafeLens resolves an extended, highly variable response structure that remains largely invisible to standard benchmarks. Because this map isolates robust and weak deflections independently at each channel under strict statistical control, it supplies a rigorously denoised prior for subsequent analyses, including network connectivity, comparisons across experimental conditions, and per-trial characterization (Section D.3). We next detail the five data-driven mechanisms responsible for this performance gap (Section D.2).

### D.2 Why SafeLens resolved more deflections than CRP and the z-scored detector

We benchmarked SafeLens against the z-scored detector and CRP. The z-scored detector represents the most common practice, since most CCEP studies that describe their detection method threshold the trial average [58]. CRP is the data-driven method closest to SafeLens, since it learns a response’s shape and duration from the trials of each stimulating and recording pair [46]. Its authors note that CRP cannot parameterize the timing of particular features, such as the latency of the first peak or trough, although that latency could still be measured on the shape [46]. They also expect the shape to represent the response’s structure, and they show that it captures the N1/N2 shape where one is present [46]. We therefore evaluated CRP’s shape within its response window against the simulated deflections, as the closest available baseline for resolving them (Section V.3). Because CRP’s parameterization stops at the shape as a whole, characterizing deflection timings from that shape goes beyond CRP’s design, so a shortfall there is outside of the primary scope of CRP. Its polarity was judged by a lenient test, which requires only that a line fitted to CRP’s shape across the deflection rise for a rising deflection and fall for a falling one (Sections V.2 and V.3). Its shape was cut into components only to measure boundary error and false components, with a smoothing chosen to favor CRP.

SafeLens resolved more than twice as many simulated deflections as CRP and the z-scored detector (Section R.3). The two benchmarks each test one statistic chosen in advance, the trial average against its pre-pulse interval and the projections between trials against zero, and use rest only to set a threshold (Section V.3). SafeLens trains its classifiers to separate trials from rest without presupposing which feature contains the response, and its lead comes from five properties of that design. Its random forests learn which rises and falls in any band and which nonlinear combinations of them separate single trials from far more windows of asynchronous rest, so a weak deflection that recurs across the trials can separate the classes. Its reliance field is read from forests trained separately at each searchlight position and tested at each time and frequency for each polarity, so it resolves where each deflection lies and whether it rises or falls. Its null distribution comes from the same chain run on set-aside rest, preventing any response leakage that would otherwise contaminate a null distribution generated via trial permutation. Its forests restrict the features each split may try, which limits how far large deflections mask weak ones, and aggregating over random draws reduces the variance that the draws add. Each held-out trial is classified against rest on its own, so trials need not have an identical waveform, a robustness that matters even without jitter because the background varies from trial to trial. Together they let SafeLens find weak and late deflections that the benchmarks’ statistics miss, beside larger deflections and across varying trials, and test them against a null free of the response.

#### Random forests trained against rest

Part of SafeLens’s sensitivity to weak deflections is attributable to its forests, which learn how single trials differ from rest in any band and any combination of features. At each searchlight position a class-balanced random forest is trained on the deflection features of event-locked and resting windows, and every held-out window is thereby transformed into a probability of event against rest (Sections M.3 to M.5). That probability addresses directly the question an analysis of evoked responses sets out to answer: whether activity after an event differs from activity without one. Each tree splits on one feature at a time, conditioned on the splits above it, so the forests can capture nonlinear relationships and interactions among features without their being specified in advance [80, 86]. A response whose evidence lies in a conjunction of features or in any one of several can therefore separate the classes, while each benchmark fixes in advance how its statistic combines the voltages (Section V.3). Because rest is asynchronous with the events, every resting window can serve as a sample of rest at every position (Section M.3). Across their trees the forests therefore learn how each feature varies without an event from far more rest windows than there are event trials. A split on a feature compares the trials’ values with the resting distribution of the same feature, so each band is weighed against its own background. A brief deflection is measured in the faster bands, where the background of field potentials is expected to have less power, so it can separate the classes while its amplitude is small against the background summed over all bands. Both benchmarks test the broadband signal, in which the background that such a deflection has to exceed includes every band (Section V.3). The deflections that SafeLens resolved and CRP did not were weaker, with a median SNR of 11.1 dB against 18.4 dB for those both resolved (Section R.4).

#### Reliance field tested at each time, band and polarity

SafeLens’s deconvolved reliance surface is tested at each time and frequency for each polarity (Section M.9), so SafeLens can report where each deflection lies and its polarity. Resolving a deflection requires both, since a deflection counts as resolved only when a significant region of matching polarity covers at least half of its amplitude change (Section V.2). Testing at that resolution costs power, since SafeLens’s family-wise control spans every time after the event and every band of a channel for both polarities, where CRP makes one decision per channel within one response window (Section V.3). SafeLens nonetheless detected more of the channels with a CCEP than CRP did (91.8% against 84.5%) and resolved more than twice as many deflections (Sections R.2 and R.3). To detect more channels while paying that cost, SafeLens’s test at each time, band and polarity must be more powerful than CRP’s single test of a channel. The polarity that resolving requires comes from measuring rises and falls separately, and consecutive deflections alternate between the two (Section V.2). The ablation sign-flips features against the direction their forest learned, so the reliance it measures is specific to rises or to falls (Section M.6). A rise and the fall that follows it are therefore read out as separate components, with their shared boundary where their fitted curves cross (Section M.10). SafeLens’s mean boundary error was 3.0% of a deflection’s duration against 16.4% for CRP (Section R.4). Each searchlight position has its own forest and window, so a deflection more than one window after a large early response is learned by forests whose windows exclude that response, and the response’s features cannot mask it there (Section M.3). CRP’s window ends where the profile of its projections peaks [46], so a late deflection after a quiet interval extends it only if it lifts the profile above its earlier peak. The deflections that SafeLens resolved and CRP did not were mostly late ones, though in this validation late deflections were also weak (Section R.3).

#### A null distribution from set-aside rest

The benchmarks’ fixed statistics can be thresholded on rest directly (Section V.3), but a map made by a learned and nonlinear chain has no null distribution in closed form. SafeLens generates one by a permutation test [23], which it applies to the chain run on set-aside rest with faux event times (Section M.9). It permutes the labels 100 times at the reliance-surface model without retraining the forests. The trials’ map is thereby tested against real rest transformed by the same steps. Permuting the labels of the trials themselves would leave part of the response in many permuted surfaces, whose maxima would then raise the threshold that the response has to cross. The response can affect the threshold only through the factor s, which matches the two branches’ scales and can only raise the threshold (Section M.9). SafeLens resolved 1,350 of the 3,701 simulated deflections when s was estimated from the maximum of each permuted map, which often falls on the response. It resolved 1,890 when s was estimated from each map’s 25th-percentile cell, as in the reference configuration (Section M.9). The false-positive rate on rest stayed at or below nominal under every summary tested. With the threshold taken instead from the trials’ own permuted surfaces, which contain part of the response, SafeLens resolved 1,292/3,701 deflections against 1,890/3,701 with the set-aside null (Section M.9). These 1,292 still exceed CRP’s 806 (p < 10^−10^), so the other four properties produce a lead of their own, which the set-aside null widens by 598 deflections. Rest recorded hours from the trials can serve for the null once centering, standardization and the scaling factor s bring the two branches to one level and scale (Sections M.8 and M.9). The family-wise control then holds provided that, with no response present, the trials’ maximum is exchangeable with the null maxima (Section M.9). Each permutation also exchanges labels only between an event-locked block and the resting block beside it, so slow changes of state are common to both. In the two subjects whose recordings underlie the simulated datasets, the null rest was recorded 3 to 7 hours after the trials, and SafeLens’s false-positive rate on rest was 5.4% (Sections V.1 and R.1). Its threshold also holds the family-wise error at 5% on each channel, where the benchmarks’ thresholds hold it on average over a dataset’s channels, so the lead does not come from a looser threshold (Sections V.3 and R.2). Neither benchmark’s published method calibrates its threshold over channels (Section V.3), and that calibration was added here for a fair comparison, since a Bonferroni correction over the z-scored detector’s correlated time points would have been conservative [23]. To our knowledge, no earlier method for evoked responses in intracranial recordings sets its family-wise threshold by training a second set of classifiers on resting data set aside from the comparison.

#### Random feature subsets at each tree node and ensemble averaging

The forests, the ablation and the reliance-surface model each aggregate over random draws, and the candidates drawn for each split limit how far large deflections mask weak ones while the aggregation reduces the variance that the draws add. Each tree is grown on a class-balanced bootstrap with one block set aside and sees a random subspace of a third of the features, of which each split tries 28 (Section M.5). The integrated reliance surface reads what the forests relied on, so a weak deflection reaches it only if the trees split on its features (Sections M.6 and M.7). A tree that tried every feature at every split would tend to split on the most discriminative ones, those of the largest deflection or of the stimulation artifact, and a feature of moderate effect would be masked [80]. At a split whose candidates include none of those features, the tree splits on the most discriminative of the others, which can be a weaker deflection’s feature. The forest’s reliance can thereby spread over the weaker deflections as well as the largest. Restricting the features each split tries also decorrelates the trees [80], which lowers the bound on a forest’s error for trees of a given strength [78]. Each interior time is read from the four forests whose windows cover it, and overlapping bands measure each deflection in more than one band (Sections M.3 and M.8). The reliance-surface model combines the four forests’ estimates and shrinks them where their votes cancel, and the deconvolution combines the bands (Sections M.7 and M.8). The model also averages 50 fits on resampled trials and trees, while each ablation scan draws its own mask (Sections M.6 and M.7). The null branch makes draws of the same kind, so its maps contain the noise that remains after the averaging, as the trials’ map does (Sections M.6, M.7 and M.9). The third of the features that each tree sees and the 28 candidates per split (the square root of the number of features [81]) are reference defaults applied unchanged to every dataset (Sections M.0 and M.5). Fewer features per tree would leave more trees without the most discriminative features, but each tree could capture fewer interactions and the forest would need more trees to cover every feature, while more features per tree would do the reverse.

#### Single trials classified against rest

The forests classify each held-out trial’s window against rest on its own, so SafeLens needs no identical waveform across the trials (Section M.5). That matters in both conditions, since even in the zero-jitter datasets each trial contained the injected response on a different segment of ongoing activity, which can shift where its deflections appear to begin and end (Section V.1). The resting windows come from the same recordings (Section V.1), so the forests learn from rest how much the background alone changes each feature and classify each trial’s window against that variation. In the high-jitter datasets the response itself also varied in latency, amplitude and shape (Sections V.1 and R.5). A tree can assign a deflection at any of several latencies to the event class, so a trial whose deflection has shifted can still raise its window’s probability of event against rest. The evidence of the shifted trials can thereby add up over the times and bands that the deflection covers across them. SafeLens’s recall did not change significantly under high jitter while CRP’s fell, and neither method’s boundary error changed significantly after correction (Section R.5). The z-scored detector’s recall did not change significantly either, but it resolved mostly the deflections of highest SNR (Fig 3B), which were mostly early ones that varied least across trials (Sections R.3 and V.1). CRP’s recall fell even against zero-jitter deflections matched in SNR (Section R.5), which points to CRP’s model, in which only the response’s amplitude varies across trials [46]. CRP’s window is also expected to shorten when the later deflections vary, since they then add less to the projections between trials that set its end, an effect not measured here. The difference between CRP’s change in recall and SafeLens’s change was not significant after correction, but this test is expected to be insensitive. Its two conditions contain different deflections, matched only in onset, duration and SNR. SafeLens’s change also includes the later and weaker deflections, which vary most across trials (Section V.1) and which CRP seldom resolved (Section R.3). A design that follows the same deflections through both conditions would compare the two changes directly.

### D.3 The SafeLens significance map as a prior for single-trial and between-condition analyses

Per-trial timing models require the number of components and their approximate positions as a prior [11, 66, 67], and the SafeLens map supplies both under statistical control. In Section R.6, the SafeLens map was tested as that prior on simulated CCEPs with strong trial-to-trial variability. With the SafeLens map as its prior, a per-trial timing model followed the true times with a large correlation for 28% of the analyzed deflections, against 11% with the standard N1 and N2 windows at their best case. With the conventional measure of latency, the time of each trial’s largest peak in a window, the share was 3%. A standard window spans several deflections, so a timing model given the window fits one series of times to all of them, and each trial’s largest peak in the window follows whichever deflection is largest on that trial (Section V.6). The windows fell behind even on the matched deflections, to which every prior gave a span. There their mean correlation was 0.30 at their best case and 0.047 by the largest peak, against 0.46 with each deflection’s true span (Section R.6). The test favored the windows, whose best case used a shift limit set from the true durations and whose deflections had correlated timing (Section V.6). With the half-window limit, which recorded data allow, the windows’ share was 8% of the analyzed deflections and their mean correlation on the matched ones 0.181 (Section R.6). CRP could supply part of that prior by cutting its canonical shape into components. On the matched deflections the timing model’s mean correlation with CRP’s components (0.404) was significantly lower than that with the SafeLens map, and CRP had not resolved 74.9% of the 926 analyzed deflections, against 25.3% for SafeLens (Section R.6). Those components are also not tested individually, since CRP’s significance test is made once per channel [46], and 30.0% of them were false in the simulation (Section R.3). The SafeLens map therefore supplies the tested prior that per-trial timing models have lacked, although its latency fidelity was measured only on simulated data, where each trial’s true latency is known (Section V.6).

The deflections SafeLens recovers beyond the reach of the two benchmarks are predominantly the weak and late ones, and it is an open question whether those are also the deflections that change most when an experiment modulates the response. The motor evoked potential shows that one response can have both a saturated and an unsaturated measure at once: during voluntary contraction, its amplitude reaches a ceiling across a range of stimulus intensities over which its duration continues to increase linearly [105]. The strong deflections of a CCEP might likewise saturate as the motor evoked potential’s amplitude does while the CCEP’s weak deflections keep changing by a larger fraction of their own size, and that has not been tested. The available intracranial plasticity work appears at first to show that larger responses change more, reporting a correlation of 0.427 between the logarithm of baseline response amplitude and the standardized size of the induced change [106]. That result is nonetheless compatible with every channel changing by the same fraction of its own response. A standardized effect size divides the change by the trial-to-trial measurement noise, leaving the response’s own amplitude out of the denominator, so a large-amplitude channel returns a larger effect size at identical proportional change. A more recent study in this line excluded weakly responding channels from its plasticity analysis by design, on the stated ground that a change in a weak signal cannot be separated from drift [107]. Testing the question therefore requires expressing change as a fraction of each channel’s own baseline response and resolving the weak components in the first place. The present work supplies the second requirement, the resolved weak components.

The recordings analyzed here offer a test of that question. Their sessions delivered TBS between some blocks of single pulses (Section V.5), and a study of how TBS changes a CCEP needs to resolve subtle changes in its morphology. Each SafeLens significance map in this paper was built from the trials of every block and condition, pooled against rest (Section M.0). Because SafeLens assumes neither a preset analysis window nor trial invariance, a deflection present in only some of the pooled trials can still enter that significance map and be tested like any other, whereas an average over all the trials would dilute it. Conditions such as the blocks before and after TBS can then be compared by per-trial timing models that take the SafeLens significance map as their prior (Section R.6). They could also be contrasted directly within the permutation test that thresholds each deconvolved reliance surface, by building a surface for each condition against its own rest and comparing the two surfaces against a null that exchanges the trials’ condition labels. In future work, we intend to apply SafeLens to these recordings in this way to resolve how TBS changes the CCEP. Whether that null separates a difference between two responses from a difference between two resting states, and whether pooling the conditions raises sensitivity over analyzing each alone, remain to be tested.

Further out, inference at the level of the individual trial becomes available with a per-trial test alone, because the per-trial probabilities are already free of train-test leakage (Section M.11). Such inference would relate a response’s trial-by-trial presence, latency, and amplitude to brain state, task demand, or tissue pathology. That extension follows from the architecture without having been a design goal. The extension remains unevaluated here, and nothing reported above depends on it. Where a response appears on only some trials [55, 56], the same per-trial test could select the responding trials and pass only those to the later steps of SafeLens, which could raise the sensitivity of those steps. The selection would be valid only if the null’s trials were selected in the same way, and neither the selection nor that check was tested here.

### D.4 CCEP duration in patient data

In the eight patients, SafeLens detected responses later after the pulse than the end of the accepted duration of a CCEP (Section R.7). That duration spans an early N1 peaking within roughly the first 50 ms and an N2 over the following few hundred milliseconds [34–36], and the longest responses characterized across a population extend to about half a second [68]. Recording time did not set that duration, since most studies leave at least a second between pulses [48, 58] and the corpus on which CRP was developed used inter-stimulus intervals of 3 to 7 s [46]. On the eight subjects’ recordings, the two benchmarks reproduced that duration, with the share of channels still responding above chance only until 455 ms for CRP and 570 ms for the z-scored detector. SafeLens’s share stayed above chance until 1040 ms (Section R.7). The three methods analyzed the same event-locked trials, so the difference between their extents comes from the detectors, each with its own filter and threshold (Sections V.3 and V.5). The accepted duration of a CCEP is therefore a lower bound set by the sensitivity of the detectors the field has used.

Three other explanations of SafeLens’s longer extent are unlikely. The first is spurious detection. The chance threshold of the cohort test already allows for a spurious run of significant time points, anywhere after the pulse, on each channel with probability 0.05 (Section V.5). SafeLens’s false-positive rate on the resting recordings that underlie the simulated datasets was 5.4%, within the range expected of a test at that 5% (Section R.1). The second is that the evidence for a response can spread past the response’s end, up to one searchlight window (Section R.1). The simulation would have shown such a spread, since the extent of every simulated deflection was known. A reported response that overlapped none of the CCEP’s deflections counted as a false positive, and an injected CCEP did not raise SafeLens’s false positives above chance (Section R.1). A spread joined to a deflection would instead have delayed that deflection’s offset, yet SafeLens’s offsets fell early more often than late on the deflections that both it and CRP resolved (Section R.4). The bound of one window, 500 ms, also overstates the spread that a late response can produce. A deflection enters the features of a later time only in the bands whose moving averages still take in part of it, and in a share that shrinks the further that time lies past the deflection’s end (Section M.3), so its evidence extends far past it only if the deflection is large. For spread to account for SafeLens’s whole margin of 470 ms over the z-scored detector, responses ending near that detector’s reach of 570 ms would have had to spread their evidence nearly a whole window further, and responses that late, at the limit of what the trial average resolves, are weak. The third is theta-burst stimulation (TBS), which the sessions delivered between some blocks of single pulses (Section V.5) and which can raise or lower cortical excitability [106, 107], so a rise could prolong the activity a pulse evokes. All three detectors received the same data, which was a mix of pre- and post-TBS CCEPs and resting-state recordings, so this should not have introduced a bias between detectors, and the comparison methods still estimate response durations the field has reported from recordings without TBS. Contrasting the blocks before and after TBS would measure any lengthening directly (Section D.3).

The late structure in the CCEP that SafeLens resolves is unlikely to be epileptiform, although the delayed responses that occur between 100 ms and 1 s after a pulse have historically been read as pathological [55, 56]. Delayed responses are associated with the seizure-onset zone [55], which in focal epilepsy is a limited region [108]. SafeLens components starting after 300 ms were, by contrast, spread over 26 of the 31 anatomical regions sampled and over 3 to 13 distinct regions within every individual subject. No subject’s largest single region included more than a third of that subject’s channels with a component starting after 300 ms. The late deflections also do not form a separate population of channels, since they appear within channels that the comparison methods already classify as responsive. Stimulus-evoked epileptiform events are moreover rare at the electrode-pair level. In a cohort of 29 patients, afterdischarges followed stimulation of 0.75% of electrode pairs outside the seizure-onset zone and 2.5% within it, and the difference did not reach significance [109]. By contrast, the stimulated pair of every one of the eight subjects evoked components starting after 300 ms. The late structure is therefore more plausibly part of the evoked response than an epileptiform event, in line with the late components reported under physiological conditions [110].

### D.5 SafeLens assumptions

SafeLens’s features are differences of moving averages, each the local rise or fall of the signal in one of 15 frequency bands, from 143 Hz down to 2.2 Hz (Section M.3). They are therefore blind to a constant offset, so a sustained shift in level registers only near where it begins and ends. The widest band is bounded by the 500 ms window of each searchlight position, and the window therefore sets the longest single deflection the features can represent. The forests then learn from the trials what separates an event-locked window from rest in these features (Section M.5). The later steps make two assumptions about the response’s shape. The first is that each deflection has a partner of opposite polarity on a similar timescale. The ablation that measures what the forests rely on tests rises and falls in alternating pairs. The deconvolution then pairs each deflection with one of opposite polarity in the same frequency band, following within a delay scaled to that band (Sections M.6 and M.8). An evoked potential alternates between rises and falls, so the pairing fits it. A deflection without such a partner cannot be paired and may be under-detected, as a step in voltage that does not reverse would be. The second is the causal prior, that no part of the response begins before the event (Section M.8). Neither assumption fixes the number of deflections, their amplitudes or their exact shapes, and only the second restricts their latencies, to the time after the event.

The component read-out, which turns the SafeLens significance map into the components evaluated in this paper, adds conventions of its own (Section M.10). The significance map keeps each frequency band separately, but the read-out collapses the bands and gives each time point one polarity. It also merges components of one polarity separated by gaps shorter than 7 ms, and drops components shorter than 2 ms. Two deflections that coincide in time but differ in frequency band (such as a sharp deflection riding on a slower one) can therefore both enter the significance map, while the read-out cannot report both whole. It would likely report them as one component if they share a polarity, and as the sharper component flanked by two parts of the slower one if they do not. The simulated responses placed their deflections in sequence, so this case is untested, and a user who expects coinciding deflections can analyze the SafeLens significance map band by band.

Analyzing each channel independently costs SafeLens statistical power, and in return its results impose no model of how the channels relate. The cost is the power that pooling across channels can supply, since methods that exploit a component’s consistent topography across contacts detect responses that a single-channel analysis misses [24, 25, 27]. Pooling of that kind requires a model of how the channels relate, and part of any coordinated structure a pooling method then reports comes from that model. For CCEPs, how the channels’ responses relate is the connectivity the experiment sets out to measure, so a pooling model would assume part of what the experiment measures. SafeLens uses no such model, so network summaries built from its significance maps (such as the raster of one subject’s components in Fig 6E) contain no structure imposed by a model of how the channels relate. These summaries still contain the false positives that each channel’s 5% family-wise error rate allows, and any covariation that volume conduction puts into the recordings. Pooling draws its power from responses that covary across channels, and the simulated responses were injected into each channel independently, so how much power pooling would add could not be measured here.

### D.6 Limitations and future directions

This study had multiple limitations. First, accuracy against ground truth could be measured only on simulated CCEPs, and how well each result translates to real CCEPs depends on what that result measures. The false-positive rate on resting recordings does not depend on the simulation, as it was measured on real resting iEEG with nothing injected (Section R.1). Channel detection needs only a response that recurs across trials and departs from rest, so it should translate to real CCEPs.

On the recordings of the eight patients, the three methods ranked in the same order as in channel detection on the simulation, with SafeLens calling the most channels responsive and the z-scored detector the fewest (Sections R.2 and R.7). Of the channels that only one of SafeLens and CRP called responsive, SafeLens called 87 and CRP 18 (Section R.7), and of the simulated channels that only one of them detected, SafeLens detected 65 and CRP 14 (Section R.2). Component recall and boundary accuracy depend most on the simulation, since they are measured on the deflections of responses generated by a neural-mass model of a cortical column [97–99]. SafeLens uses none of the model’s particular shapes, only the properties set out in Section D.5, and the model’s responses share those properties with CCEPs. They were also added to real resting recordings, so their background contained the noise and rhythms of real iEEG. The additive injection assumes that an evoked response adds to ongoing activity, whereas a real CCEP depends on the state of the tissue it arrives in [44, 47–52]. The model also couples later latencies to slower components, so its latency-spectrum gradient is partly built in and no spectral result is reported here. A real CCEP response that departs from the simulated ones may be resolved less accurately than the simulation measured. A late component absent from some trials [55, 56] is one example, and widening the simulation to include it is among the next steps below.

Second, the simulation used the resting recordings of only two subjects, so its 704 channels and 3965 deflections share two resting backgrounds. The tests of Section V.4 treat the channels of one subject as independent, so their p-values may be too small. The advantage over CRP nonetheless held within each subject, with component recall of 46.8% against 17.6% in one and 48.8% against 19.1% in the other (Section R.3). The eight patients add further subjects to the comparison of the three methods, though without ground truth (Section R.7).

Third, SafeLens’s control of false positives in real CCEP data depends on how closely the resting data set aside for the null (the secondary recordings, Section M.1) resemble the rest recorded between CCEP blocks. In the two subjects whose recordings underlie the simulated datasets, the secondary asynchronous data were recorded 3 to 7 hours after the recording into which the CCEPs were injected, after clinical CCEP mapping, and before and after TBS (Section V.1). The false-positive rate was still 5.4%, within the range expected at its nominal 5% (Section R.1). In the eight patients, they were the rest recorded before the first block of pulses or after the last (Section V.5). The threshold is also multiplied by a factor s, never below 1, which rises when the recording’s deconvolved reliance surfaces vary more widely than the null’s (Section M.9). Resting data from a different state, such as sleep in place of wakefulness, were not tested. Because s raises the threshold when the recording varies more widely than the null and never lowers it, we expect such data to cost sensitivity before they raise the false-positive rate. The factor s can also grow with a strong response, part of which survives the permutations of the recording from which s is estimated, and a larger s likewise costs sensitivity without raising the false-positive rate. Beyond that match of scales, the family-wise control rests on two exchangeability assumptions (Section M.9). The recording’s maximum must be exchangeable with the null’s maxima when no response is present, which the false-positive rate measured on rest supports (Section R.1). The null’s maxima come from exchanging the event-locked and resting labels of trials only within each pair of an event-locked block and its resting block, so those labels need be exchangeable only among trials recorded close together in time [93].

Fourth, the simulated stimulation artifact was a smooth pulse of 1 ms rise and 10 ms duration (Section V.1), whereas a real electrical pulse can produce a much larger artifact. SafeLens measures each frequency band with moving averages centered on each time point, wider for the slower bands. A large, sharp excursion therefore spreads into the slower bands up to half an average’s width before it begins (Section M.3), and by as much after it ends. The causal mask removes what spreads to before the pulse (Section M.8), and on the simulated responses SafeLens’s onsets had a median signed error of zero (Section R.4). An excursion much larger than the simulated responses, such as a large stimulation artifact or an unusually strong early deflection, may still add reliance in the slower bands just after the pulse and so place a response’s first onset early. Simulating realistic artifacts is among the next steps.

Fifth, while SafeLens runs offline, it analyzes each channel on its own and requires a GPU to run efficiently. Its channels can be divided across GPU nodes (Section M.0), and it also runs without a GPU, more slowly. The 704-channel simulated validation took ~62 hours of machine time on the workstation described in S4 Table, and a whole-brain, multi-subject study takes days. SafeLens was also tested only at the trial counts of these datasets, thirteen or fourteen blocks of about 26 pulses in the simulation (Sections M.2 and V.1) and 276 to 378 pulses per channel in the patients (Section V.5). How its sensitivity falls with fewer trials is among the next steps.

Several of these limitations lead naturally to the next steps. The application to eight patients can be extended to a larger and more heterogeneous cohort and to other stimulation paradigms, so that the late structure SafeLens resolved can be characterized across datasets. The simulation can be widened to the resting recordings of more subjects, to responses absent from some trials or covarying across contacts, to deflections that coincide in time, to stimulation artifacts of realistic size and to fewer trials. Covariance across contacts would also allow a comparison with methods that pool across channels (Section D.5). Within SafeLens, varying each part of the design named in Section D.2 would measure its contribution. Examples are a permutation null built from the recording itself, trees that see every feature and a significance map made without the deconvolution, each compared with the full pipeline. The resting null permutes the labels of windows that null forests trained once have classified (Section M.9), and its false-positive rate on rest held within the range expected at the nominal 5% (5.4%, Section R.1). Retraining the forests under each permutation would check directly that training them once leaves the null unchanged. The deconvolution’s pairing could also be generalized to deflections that have no partner of opposite polarity (Section D.5).

## Conclusions

SafeLens trains a random forest at every searchlight position with resampling at every level of training, reads out what the forests rely on by ablation without retraining them, and runs a resting null through the entire pipeline. Together these three techniques put each reported deflection under the channel’s family-wise error control, and the analysis remains practical on one workstation (Section M.0). A CCEP can therefore be reported as a set of separately tested deflections, each with its presence, extent, and polarity under that control, which to our knowledge no existing CCEP method provides [58]. On simulated CCEPs, SafeLens resolved 47.9% of the 3,965 evaluated ground-truth deflections with the correct polarity against 18.4% for CRP, and 2.8% of its reported components were false against 30.0% of CRP’s (Section R.3). SafeLens placed boundaries at a mean error of 3.0% of the deflection’s own duration against 16.4% for CRP, and its advantages in recall and boundary accuracy held when trials varied (Sections R.4 and R.5). With the SafeLens map as its prior, a per-trial timing model followed the true times with a large correlation for 28% of the analyzed simulated deflections, against at most 11% with the standard N1 and N2 windows (Section R.6). In the recordings of eight subjects, the response structure SafeLens resolved remained above chance at the cohort level to 1040 ms after stimulation, against 455 ms for CRP on the same recordings (Section R.7). The duration the field has accepted for a CCEP therefore reflects the sensitivity of the detectors used to measure it (Section D.4).

Beyond CCEPs, SafeLens is designed for an event that repeats at known times with responses that end before the next event, given a recording that supplies one time series per site and enough resting data to build its null (Sections M.0 and M.1). Nothing in its construction is therefore specific to CCEPs or to electrophysiology, although SafeLens has been tested only on intracranial recordings. A site may be an electrode contact, an imaging pixel or a voxel. TMS-EEG, in which a magnetic pulse evokes a response recorded at scalp electrodes, is an immediate candidate for SafeLens. The null construction also transfers to any nonlinear detector without a closed-form null, since a complex processing chain run in full through an extensive empirical resting null becomes family-wise-error-controllable, provided that null is exchangeable (Section M.9).

## Data and Code Availability

SafeLens runs in MATLAB and requires the Parallel Computing Toolbox, the Statistics and Machine Learning Toolbox and a CUDA-capable GPU, on which feature extraction, the ablation scans and the largest linear solves run. The SafeLens code will be made publicly available under an open-source license no later than this study’s publication in a peer-reviewed journal. The data supporting the findings of this study are available from the corresponding author upon reasonable request, subject to approval by the Stanford University IRB.

## Ethics Statement

Eight patients with medically intractable epilepsy underwent a surgical procedure at Stanford University Hospital to implant intracranial recording contacts within brain parenchyma (stereo-EEG). Contacts were placed in accordance with clinical need to localize epileptic regions. All patients gave their written informed consent for the surgical procedure and to participate in brain stimulation research during their hospital stay. Research was approved by the Stanford University IRB, as described previously for this cohort [103]. The simulated datasets (Section V.1) were built on resting recordings of two of these patients.

## Author Contributions

**Conceptualization**: Christopher B. Minasi, Christopher C. Cline, Corey J. Keller; **Data curation**: Christopher B. Minasi; **Formal analysis**: Christopher B. Minasi; **Funding acquisition**: Corey J. Keller; **Investigation**: Christopher B. Minasi, Umair Hassan; **Methodology**: Christopher B. Minasi, Christopher C. Cline, Corey J. Keller; **Project administration**: Christopher B. Minasi, Christopher C. Cline, Corey J. Keller; **Resources**: Corey J. Keller; **Software**: Christopher B. Minasi; **Supervision**: Christopher C. Cline, Corey J. Keller; **Validation**: Christopher B. Minasi; **Visualization**: Christopher B. Minasi; **Writing – original draft**: Christopher B. Minasi; **Writing – review & editing**: Christopher B. Minasi, Christopher C. Cline, Corey J. Keller.

## Acknowledgments

We thank Josef Parvizi for his contribution to patient recruitment, imaging analysis including the anatomical labeling of the recording contacts, and the facilitation of research time with the neurosurgical patients at Stanford University. We also thank the patients who took part in this research.

