## Supporting information: S1-S3 Figs and S1-S6 Tables for "SafeLens: A Statistically Controlled Nonlinear Machine Learning Framework for Characterizing Hidden Dynamics in Event-Locked Neural Activity"

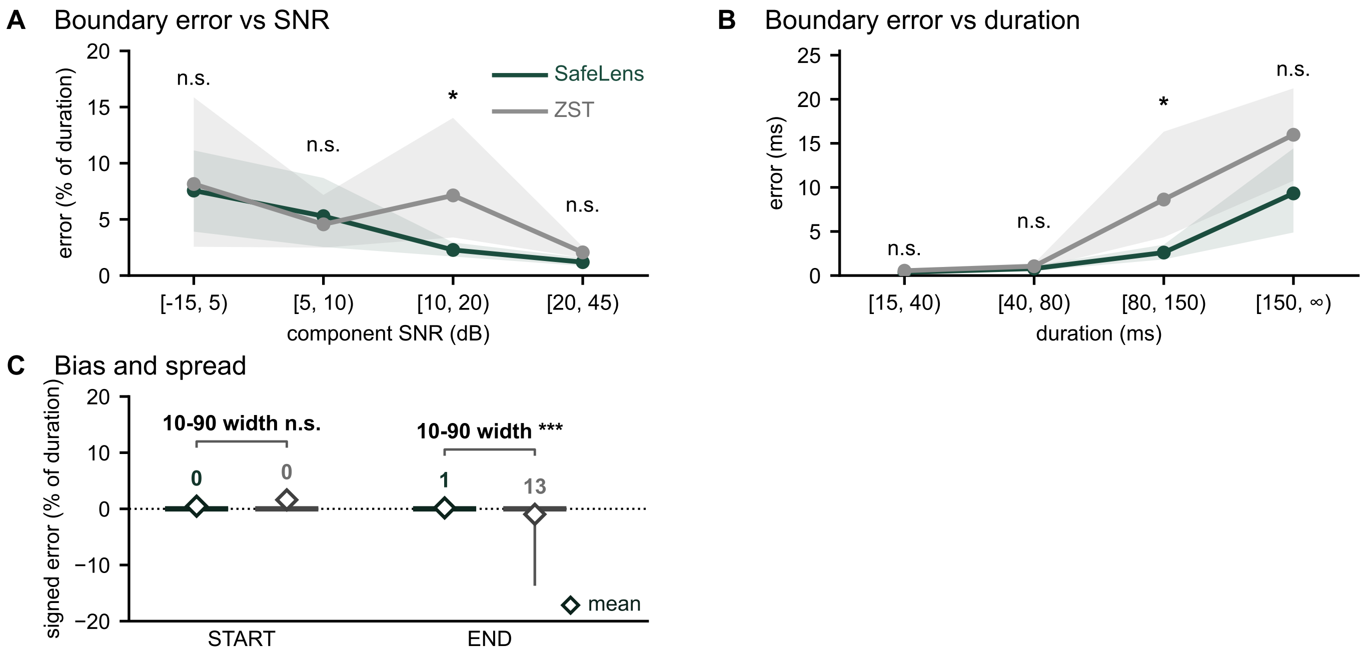


**S1 Fig. Boundary error of SafeLens and ZST on the simulated deflections.** The panels repeat Fig 4C to E for SafeLens and the z-scored trial-averaging detector (ZST), on the 745 deflections both resolved (Section V.2). **A.** Boundary error against component SNR (n = 6, 39, 300 and 400 from the lowest bin to the highest). Lines are means with 95% bootstrap bands over recorded channels, and each bin has a sign-flip test of each channel's mean error difference between SafeLens and ZST, Bonferroni-corrected over bins (* $p < 0.05$, *** $p < 0.001$; n.s., not significant). **B.** The same error in milliseconds against ground-truth duration (n = 141, 330, 224 and 50), with bands and tests as in (A). **C.** Signed error at each onset (START) and offset (END), negative when the edge falls before its tolerance band and zero inside it. Boxes span the 25th to 75th percentiles with a line at the median, and whiskers span the 10th to 90th percentiles. The number above each whisker is that span in percentage points, and brackets compare the spans by swapping the methods' errors within recorded channels, Bonferroni-corrected over the two.

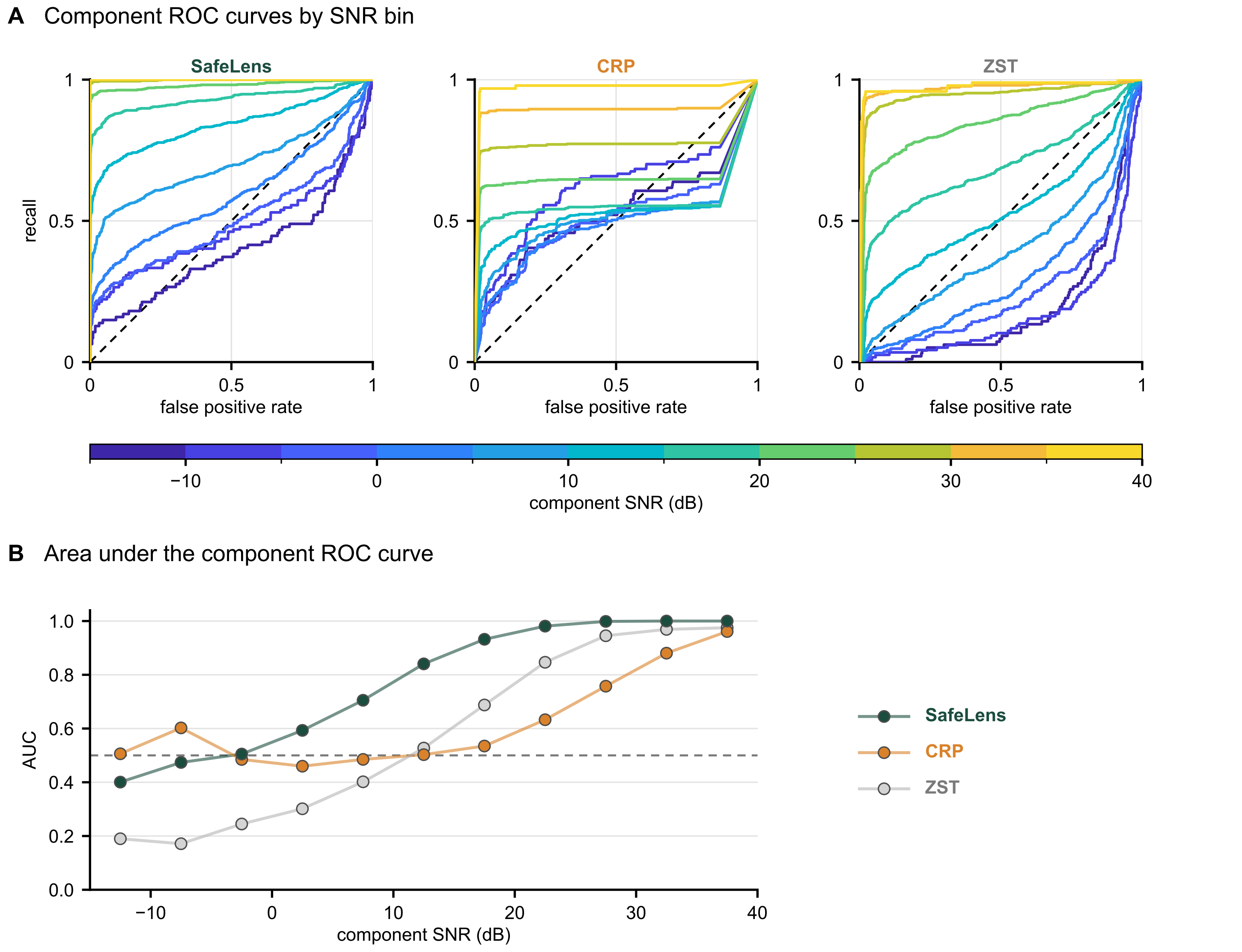


**S2 Fig. Component-level ROC.** SafeLens is compared with canonical response parameterization (CRP) and the z-scored trial-averaging detector (ZST). Pooled over the 704 channels of the four simulated datasets with an injected CCEP (response channels) and the same channels' secondary recordings at the faux event times (null channels), with the component as the unit of analysis. A component is a segment of an injected response, cut at the minimum of the local SNR between neighboring peaks, and its SNR is the largest local SNR inside it (Section V.1). Each method's value for a component is the largest value of its statistic inside it. The segments are cut from the ground-truth response window for all three methods. The negatives are the largest value on each null channel and the largest positive value in each segment of a response channel with a local SNR below −30 dB. **A.** Receiver-operating characteristic (ROC) curves of each method, one per 5 dB bin of component SNR. The lowest bin contains every component below −10 dB and is drawn from −15 to −10 dB, and the eight components above 40 dB are left out. SafeLens and CRP have 4,435 components and ZST 4,437, with 1,610, 830 and 1,610 negatives. CRP's statistic is zero outside its own response window (Section V.3), so all the components and negatives outside that window share the value zero. No threshold separates tied values, so each of CRP's curves ends in a straight line to the top right. **B.** Area under the ROC curve (AUC) per bin, counting a tie between a component and a negative as a miss, so that CRP's AUC lies below the area under its curve in (A). Dashed lines mark chance, the diagonal in (A) and 0.5 in (B).

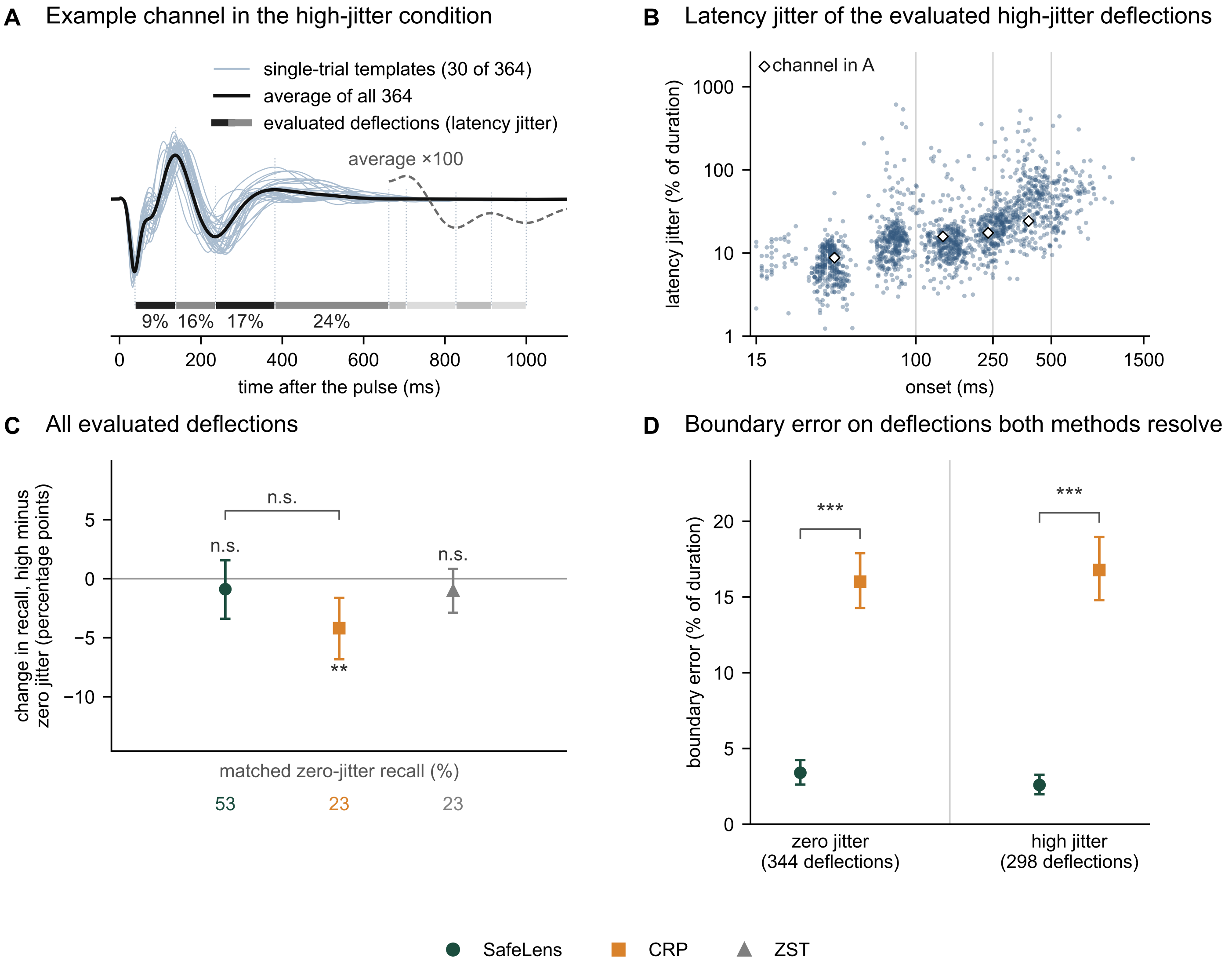


**S3 Fig. Recall and boundary error of SafeLens, CRP and ZST with zero and high jitter.** SafeLens is compared with canonical response parameterization (CRP) and the z-scored trial-averaging detector (ZST). **A.** An example channel in the high-jitter condition, typical in latency jitter. Thin lines are noise-free templates injected at its pulses. Bars in alternating shades and dotted lines mark the evaluated deflections and their edges. The dashed line is the average, magnified, from the fifth evaluated deflection on. Where the single trials are visible, each bar is labeled with the deflection's latency jitter, the standard deviation of its latency across trials as a percentage of its duration. Deflections starting within 15 ms of the pulse are not evaluated. A response ends at its first pair of deflections at or below −15 dB, and those after it (faded bars) are not evaluated. **B.** The 1,743 evaluated high-jitter deflections with a defined latency jitter. Vertical lines mark the edges of the onset bands used for matching. White diamonds mark the deflections of the channel in (A). **C.** Each method's change in recall from zero-jitter deflections matched on onset, duration and SNR to the 1,736 high-jitter deflections. **D.** Mean boundary error of SafeLens and CRP on the deflections that both resolved (Section V.2). Error bars in (C) and (D) are uncorrected 95% bootstrap intervals over recorded channels. In (C) each method's change is tested against zero, and the bracket compares CRP's change with SafeLens's, by a permutation that swaps each recorded channel's zero- and high-jitter deflections; in (D) the brackets compare SafeLens and CRP by a sign-flip test over recorded channels; all Bonferroni-corrected (*** $p < 0.001$, ** $p < 0.01$, n.s. not significant).

| Group | Modality (PubMed search term) | Records | Share |
|---|---|---|---|
| **Electrical and magnetic** | `"Electroencephalography"[MeSH] NOT "Electrocorticography"[MeSH]` | 16,011 | |
| | `"Magnetoencephalography"[MeSH]` | 1,063 | |
| | `"Electrocorticography"[MeSH] OR "intracranial EEG"[tiab] OR "stereo-EEG"[tiab] OR "stereoelectroencephalography"[tiab] OR ECoG[tiab]` | 719 | |
| | `"single-unit"[tiab] OR "single unit recording"[tiab] OR "microelectrode"[tiab] OR "spike train"[tiab]` | 748 | |
| | **subtotal** | **18,541** | **56.0%** |
| **Hemodynamic** | `"functional magnetic resonance"[tiab] OR fMRI[tiab] OR "BOLD"[tiab]` | 4,072 | |
| | `"Spectroscopy, Near-Infrared"[MeSH] OR fNIRS[tiab] OR "near-infrared spectroscopy"[tiab]` | 646 | |
| | **subtotal** | **4,718** | **14.2%** |
| **Optical** | `"calcium imaging"[tiab] OR "Ca2+ imaging"[tiab] OR GCaMP[tiab]` | 691 | |
| | `"fiber photometry"[tiab] OR "fiber photometry"[tiab]` | 115 | |
| | **subtotal** | **806** | **2.4%** |
| **Effector and autonomic** | `"Evoked Potentials, Motor"[MeSH] OR "motor evoked potential"[tiab] OR "Electromyography"[MeSH]` | 8,141 | |
| | `pupillometry[tiab] OR "pupil dilation"[tiab] OR "pupillary response"[tiab]` | 383 | |
| | `"Galvanic Skin Response"[MeSH] OR "skin conductance"[tiab] OR "electrodermal"[tiab]` | 318 | |
| | **subtotal** | **8,842** | **26.7%** |
| **Four groups** | **combined** | **32,907** | **99.3%** |
| **Set aside** | `"Positron-Emission Tomography"[MeSH]` | 122 | |
| | `"functional ultrasound"[tiab] OR "fUS imaging"[tiab]` | 42 | |
| | `"voltage imaging"[tiab] OR "genetically encoded voltage"[tiab] OR "dLight"[tiab] OR "GRAB sensor"[tiab] OR "neurotransmitter sensor"[tiab]` | 61 | |
| | **subtotal** | **225** | **0.7%** |
| **Total** | | **33,132** | **100%** |

**S1 Table. Prevalence of recording modalities in the event-locked literature.** Counts supporting the Introduction's statement that the event-locked design spans the field's recording modalities, at more than thirty-three thousand publications indexed since 2015. Each modality term shown was combined with a common event-locked design term, ("evoked potentials"[MeSH] OR evoked[tiab] OR "event-related"[tiab] OR "event related"[tiab] OR "task-related"[tiab] OR "stimulus-locked"[tiab]), and restricted to publication years 2015 to 2025. Counts were retrieved from the PubMed E-utilities interface on 29 July 2026 and are reproducible by re-running the queries. Two properties of the count should be read alongside it. First, the modality terms are not mutually exclusive, so a study using two modalities contributes to both rows, and the shares are shares of the summed record count rather than of a partition into distinct articles. Second, no species restriction was applied, because the optical methods in particular are used predominantly in animal work. Restricting the same queries to humans[MeSH] raises the four-group share from 99.3% to 99.6% and does not change the conclusion. The count is offered as an order-of-magnitude justification for which modalities the Introduction treats in detail, and not as a bibliometric result in its own right. In the search terms, [MeSH] matches the articles indexed under that Medical Subject Headings term or a narrower one, and [tiab] the articles whose title, abstract or author keywords contain the term.

| Step | Quantity | Value |
|---|---|---|
| **Corpus** | panel research articles, 2015–2025 | 39,155 |
| | matching event-design vocabulary | 17,780 (45.4%) |
| **Validation sample** | articles drawn across the decade | 100 |
| | labeled event-locked (physiological recording) | 64 |
| | labeled event-based, behavioral read-out only | 3 |
| | labeled non-event | 33 |
| | vocabulary match, checked mechanically | 46 |
| **Classifier calibration** | sensitivity, any event design | 43/67=0.64 |
| | specificity | 30/33=0.91 |
| **Corrected prevalence** | **any event or trial design** | **66% (95% CI 51–81%)** |
| | physiological event-locked recordings | 63% |

**S2 Table. Prevalence of event-locked designs in core systems and cognitive neuroscience journals.** Counts supporting the Introduction's statement that event-locked studies make up roughly two thirds of the research articles in the field's core journals. Inclusion. The corpus is every research article indexed 2015–2025 in eight journals whose stated scope is systems, cognitive, or behavioral neuroscience with primary in-vivo recordings, chosen to span the field's sub-communities: J. Neurosci., Nat. Neurosci., Neuron, NeuroImage, J. Cogn. Neurosci., Psychophysiology, J. Neurophysiol., and Cereb. Cortex (journal article[pt], excluding reviews, comments, editorials, and errata). Defining the corpus by journal scope rather than by topic keywords avoids circularity: no search term related to the quantity being measured enters the denominator. Exclusion. Venues whose scope is primarily cellular and molecular neuroscience (in-vitro and culture work), disease-centered clinical research, or reviews fall outside the panel by construction; within the panel, articles on resting-state activity, anatomy, molecular mechanisms, or theory without an event design are not excluded in advance but are counted by the classification itself. Estimate. An article counts as event-locked when it analyzes a recorded physiological or neural signal relative to repeated discrete events, the design of Introduction paragraph 1. A title/abstract vocabulary match (task, stimulus/stimuli, cue(s), trial(s), paradigm, evoked, event-related, stimulus-locked, ERP(s)) flags 45.4% of the corpus, but vocabulary is not design: in a validation sample of 100 articles drawn in five slabs across the decade and labeled against the definition above, a third of true event-design articles use none of these words (sensitivity 0.64, largely animal-systems phrasing such as responses to whisker deflection), while false positives are few (specificity 0.91). Correcting the corpus-wide rate for the measured classifier error, $(0.454 + 0.909 - 1)/(0.642 + 0.909 - 1) = 0.66$, gives 66% (95% CI 51–81% by the delta method), in agreement with the sample's own direct rate (67/100). The sample's mechanical vocabulary rate (46/100) matches the corpus rate (45.4%), the representativeness check this sampling permits. Counts were retrieved from the PubMed E-utilities interface on 14 August 2026 and are reproducible by re-running the queries. The estimate is offered as an order-of-magnitude statement of how much of systems and cognitive neuroscience the event-locked design organizes, and not as a bibliometric result in its own right: sample labels were assigned from abstracts by rubric, and the panel's flagship journals include cellular systems work, which lowers the share relative to a strictly cognitive panel.

| Stage | Parameter | Reference value |
|---|---|---|
| **Deflection pyramid (M.3)** | per-step growth cap $\Delta_{\max}$ | 50 samples |
| **Reliance GLM (M.7)** | time-knot spacing | $\max(5 \text{ samples}, \min_k S_k) =$ 5 ms |
| **Standardization before the deconvolution (M.8)** | null noise-floor percentile | 10th |
| | noise-floor $\varepsilon$ | 0.1 |
| **Cross-band smoothing (M.8)** | cross-band Gaussian $\sigma$ | 1.0 |
| **Read-out grid (M.6–M.8)** | shift-decimation factor | 1 (reliance grid = training grid) |

**S3 Table. Secondary reference parameters.** Fixed constants of the reference configuration that are set in the released code. $\Delta_{\max}$ is tied to the reference window $\omega = 500$ samples and must be edited if $\omega$ changes. The Monte-Carlo counts ($N_{\text{trees}}$, $N_{\text{scan}}$, GLM bags, $N_{\text{perm}}$) are stated in the main text.

| Stage | Time per channel | Limiting resource | Parameters that alter run time |
|---|---|---|---|
| **Forest training (M.5)** | **One lane**. 13.8 min<br>**11 lanes**. 1.44 min | CPU cores | Linearly in each: pulses, searchlight positions and trees<br>Unaltered: permutations |
| **Phase-barcode ablation (M.6)** | **One lane**. 2.2 min<br>**11 lanes**. 0.18 min | GPU compute | Linearly in each: pulses, searchlight positions, trees and ablation scans ($N_{\text{scans}}$)<br>Unaltered: permutations and sampling rate |
| **Reliance-surface GLM and deconvolution with their null (M.7–M.9)** | **One lane**. 11.9 min<br>**11 lanes**. 3.65 min | host memory | Linearly, through the solve's largest array (S5 Table): pulses<br>Sublinearly, 2.1 times as long for 101 evaluations (the unpermuted labels and 100 permutations) as for one: permutations<br>Linearly, two solves each (primary and null branch): blocks |
| **Whole pipeline** | **One lane**. 29.9 min<br>**11 lanes**. 5.30 min | | |

**S4 Table. Measured compute time per channel and the resource that limits each stage.** Time per channel at the reference configuration (500 ms searchlight window, the 33 searchlight positions that a 3.406 s inter-pulse interval admits, 1 kHz sampling, $N_{\text{trees}} = 300$, $N_{\text{scans}} = 300$, $N_{\text{perms}} = 100$, and seven-fold cross-validation, each fold holding out a run of adjacent blocks), measured on one workstation with 32 physical cores, 128 GB of host memory and one NVIDIA RTX 3090 GPU (24 GB). A lane is one MATLAB process with four parallel workers, which runs one stage on a list of channels from one subject. Channels are processed independently of one another, so the numbers of channels and subjects alter the total time but not the time per channel. For one lane, the table gives the time of one channel of a simulated dataset (14 blocks of about 26 pulses) processed alone, each stage in one lane in turn, with the start-up of MATLAB and its workers. For 11 lanes, the table gives the machine time per channel of the final run on the eight patients (651 channels, 276 to 378 pulses per channel): its 57.5 hours of wall-clock time, with 11 lanes running on average and up to 17 at once, divided by its 651 channels, and each stage's share of that time. The whole-pipeline row also includes combining the blocks' maps into one map per channel (1.5 min for the channel processed alone and 0.03 min per channel in the final run) and, for the channel processed alone, 0.5 min between the stages. The limiting resource is the one that sets a stage's throughput. Forest training runs in parallel across the CPU cores, so its throughput follows the machine's core count. The GLM and deconvolution solves run on the GPU, but host memory limits how many of their lanes run at once: each solve needs about 1.9 GB of GPU memory and a lane runs one solve at a time, and throughout the final run the scheduler's estimates allowed 3 to 14 of these lanes at a time by host memory and 17 or more by GPU memory. S5 Table gives the host memory (RAM) and GPU memory per lane.

| Stage | Host memory per lane | GPU memory per lane | Parameters that alter memory consumption |
|---|---|---|---|
| **Forest training (M.5)** | 4.9 GB | 0.7 GB | Linearly in each: searchlight positions and trials<br>Nearly linearly: sampling rate (halving the sampling rate leaves 0.49 of the memory) |
| **Phase-barcode ablation (M.6)** | 4.6 GB | 1.0 GB | Linearly: trials<br>Unaltered: permutations and sampling rate |
| **Reliance-surface GLM and deconvolution with their null (M.7–M.9)** | 6.8 GB | 2.8 GB | Linearly in each, through the solve's largest array: ablation scans ($N_{\text{scans}}$), searchlight positions, trials per block and evaluations |
| **Whole pipeline** | **One lane**. 9.7 GB<br>**16 lanes**. 79 GB | **One lane**. 2.8 GB | |

**S5 Table. Memory per lane.** Each stage's peak host memory and peak GPU memory, measured on the workstation of S4 Table with one channel of a simulated dataset processed alone, each stage in one lane in turn. A lane's host memory is the private working set of its MATLAB process and four parallel workers, the memory they use and share with no other process, so each further lane adds that amount. A lane's GPU memory is the card's memory in use above what was in use before the run. Channels are processed independently, so a lane's memory does not grow with the number of channels it processes. The whole-pipeline row gives all the pipeline's processes together. For the channel processed alone, its host memory is the sum of the processes' working sets: the GLM and deconvolution lane's 6.8 GB, about 1 GB for the scheduler (the MATLAB process that starts the lanes) and about 2 GB that the processes share, such as program code. For 16 lanes, it is the most host memory in use that the scheduler recorded during the final run on the eight patients (S4 Table), less the 24.4 GB in use before the run. The 16 lanes were 7 training, 8 GLM and deconvolution and 1 ablation lane, and their 79 GB is less than the sum of their peaks, 93 GB, since the lanes do not all peak at once. The largest array of the GLM solve contains one value for each combination of ablation scan, searchlight position, held-out trial and evaluation (the unpermuted labels and each permutation) of one block, so its size grows linearly with each of these numbers. The resource guide distributed with the code gives the size of every allocation and the line of code that makes it.

| Deflection starts | Deflections | Latency jitter, mean ± SD | Median [IQR] | 10th–90th percentile | Above 50%, n (%) | Above 100%, n (%) | Adjacent pairs with overlapping latencies, k of n (%, 95% CI) |
|---|---|---|---|---|---|---|---|
| **15–100 ms** | 744 | 16% ± 36% | 10% [7–14%] | 5–20% | 27 (3.6%) | 12 (1.6%) | 22 of 392 (5.6%, 3.6–8.4) |
| **100–250 ms** | 502 | 20% ± 35% | 14% [10–19%] | 8–28% | 20 (4.0%) | 10 (2.0%) | 42 of 501 (8.4%, 6.1–11.2) |
| **250–500 ms** | 672 | 52% ± 64% | 38% [22–60%] | 15–97% | 247 (36.8%) | 63 (9.4%) | 284 of 667 (42.6%, 38.8–46.4) |
| **500–1500 ms** | 722 | 105% ± 196% | 64% [43–95%] | 29–164% | 473 (65.5%) | 163 (22.6%) | 564 of 679 (83.1%, 80.0–85.8) |

**S6 Table. Latency jitter of the simulated deflections in the high-jitter condition.** A deflection's latency jitter is the standard deviation, across the 500 trials simulated for its channel, of the time of the deflection's steepest point, as a percentage of the deflection's duration in the trial average (Section V.1). The deflections are the high-jitter condition's ground-truth deflections starting 15 ms or more after the pulse, a rise or a fall each, whose 500 trials' average changes in the deflection's direction over its span. There are 2,641 in 352 channels, and the one starting after 1,500 ms is left out of the rows. On each trial, a deflection is matched to the rise or fall of the same kind whose steepest point lies nearest its steepest point in the trial average. The 200 of the library's 250,000 trials whose template exceeded the single-precision maximum are left out, since they were injected as zeros. Above 50% and above 100% count the deflections whose latency jitter exceeds half or all of their duration. An adjacent pair of deflections overlaps when the later deflection's 5th latency percentile precedes the earlier deflection's 95th, so that on some trials the later deflection comes before the earlier one does on others. Each pair is counted in the row of its later deflection. Intervals are exact (Clopper–Pearson). In the zero-jitter condition every deflection's latency jitter is 0% (all 565 deflections of the 50 channels checked).
